# A Unifying Statistical Framework to Discover Disease Genes from GWAS

**DOI:** 10.1101/2022.04.28.489887

**Authors:** Justin N.J. McManus, Robert J. Lovelett, Daniel Lowengrub, Sarah Christensen

## Abstract

Genome-wide association studies (GWAS) identify genomic loci associated with complex traits, but it remains an open challenge to identify the genes underlying the association signals. Here, we extend the equations of statistical fine-mapping, to compute the probability that each gene in the human genome is targeted by a causal variant, given a particular trait. Our computations are enabled by several key innovations. First, we partition the genome into optimal linkage disequilibrium blocks, enabling genome-wide detection of trait-associated genes. Second, we unveil a comprehensive mapping that associates genetic variants to the target genes they affect. The combined performance of the map on high-throughput functional genomics and eQTL datasets supersedes the state of the art. Lastly, we describe an algorithm which learns, directly from GWAS data, how to incorporate prior knowledge into the statistical computations, significantly improving their accuracy. We validate each component of the statistical framework individually and in combination. Among methods to identify genes targeted by causal variants, this paradigm rediscovers an unprecedented proportion of known disease genes. Moreover, it establishes human genetics support for many genes previously implicated only by clinical or preclinical evidence, and it discovers an abundance of novel disease genes with compelling biological rationale.

## INTRODUCTION

Genome-wide association studies (GWAS) have identified ~10^5^ genomic loci associated with complex traits across the spectrum of human disease and physiology (Price et al., 2015; Visscher et al., 2017; Buniello et al., 2019; Loos, 2020). If we could identify the specific genetic polymorphisms driving each association (i.e., the ‘causal variants’), and link those to the genes they affect, we would uncover the genetic architecture of complex traits and disease. In particular, the ability to identify genes targeted by causal variants would revolutionize drug development and unveil the molecular mechanisms underlying phenotypic variation. Indeed, sophisticated computational strategies already generate hypotheses about which genes may underlie GWAS associations, and in some cases can directly identify genes targeted by causal variants (Raychaudhuri et al., 2009; Rossin et al., 2011; Pers et al., 2015; Taşan et al., 2015; Greene et al., 2015; Giambartolomei et al., 2014; de Leeuw et al., 2015; Gamazon et al., 2015; Gusev et al., 2016; Zhu et al., 2016; Hormozdiari et al., 2016; Barbeira et al., 2018; Mancuso et al., 2019; Nasser et al., 2021).

Nevertheless, the discovery of trait-associated genes from GWAS has been hindered by several compounding challenges, which impede the identification of causal variants and their target genes. First, GWA studies carry a heavy multiple comparisons burden. At current sample sizes, many causal variants induce association signals below the threshold for genome-wide significance (Pe’er et al., 2008; Yang et al., 2015; Visscher et al., 2017). Then, even when a genome-wide-significant (GWS) association is discovered, it is often difficult to pinpoint the causative genetic variant(s). Nearby variants exhibit linkage disequilibrium (LD), which is to say their alleles do not assort independently (Gabriel et al., 2002a; Wall and Pritchard, 2003; Slatkin, 2008). The resulting correlations between their alleles (Gabriel et al., 2002b; Berisa and Pickrell, 2016) create corresponding correlations in their GWAS signals. LD therefore entangles the association signals from nearby variants, making it difficult to tease apart the causal polymorphisms, which drive each association, from the correlated ‘passenger’ variants that have no effect on the trait (Eilbeck et al., 2017; Schaid et al., 2018). And then, even when a causal variant can be identified, ~90% of these are non-coding (Maurano et al., 2012); it is typically unknown which genes they affect (Elkon and Agami, 2017).

A convergence of mathematical and experimental advances has now made these problems tractable. Statistical fine-mapping algorithms have established the theory to dissect causal variants from complicated GWAS associations (Servin and Stephens, 2007; Hormozdiari et al., 2014; Chen et al., 2015, 2016; Pickrell, 2014; Kichaev et al., 2014; Kichaev and Pasaniuc, 2015; Kichaev et al., 2017; Benner et al., 2016; Wang et al., 2020; Weissbrod et al., 2020). In addition, functional genomics technologies have uncovered the mechanisms of gene regulation (Andersson et al., 2014; Rao et al., 2014; Elkon and Agami, 2017; The ENCODE Project Consortium et al., 2020; Meuleman et al., 2020; GTEx Consortium, 2020), thereby paving the way for computational methods to link variants to the genes they affect. Here, we combine these advances into a novel statistical framework, which tackles the major problems of GWAS analysis head-on.

We posit that the most direct approach to discover trait-associated genes is to uncover the causal variants that induce GWAS associations, and then identify the ‘target’ genes affected by those variants. We implement this idea in a probabilistic setting that does not require literally pinpointing the causal polymorphisms (which is usually not possible at current GWAS sample sizes). In particular, we extend the equations of statistical fine-mapping from the domain of *variants* to *genes*, by incorporating a comprehensive mapping (i.e., lookup table) from variants to the target genes they affect. We thereby compute the posterior probability of association (PPA) for genes; namely, the probability that genes are targeted by a causal variant. Furthermore, we introduce an optimal partition of the human genome into mutually uncorrelated LD blocks, which enables genome-wide application of the fine-mapping equations. For a given complex trait, we can therefore compute the probability of association for every gene in the genome, making it possible to identify disease genes implicated by sub-GWS associations.

## RESULTS

### LD Blocks in the Human Genome

In order to express the genome-wide fine-mapping equations in a computationally tractable form, we leverage the statistical structure of the human genome. Linkage disequilibrium (LD) induces correlations between the alleles of nearby variants. LD follows a well-known spatial pattern, occurring in discrete intervals (blocks) separated by recombination hotspots (Berisa and Pickrell, 2016). We can therefore define a partition of the human genome into contiguous, non-overlapping, mutually uncorrelated LD blocks that collectively span the genome. Each block circumscribes a set of mutually correlated alleles that are uncorrelated with sites in other blocks. The problem of finding such a partition can be formulated mathematically, and its exact solution is given by the global optimum of a graph clustering problem (Figure S1). We present an algorithm to efficiently obtain the solution to this LD partition problem (see the *Methods*). At the global optimum, variants tend to co-cluster into the same LD block if and only if their sample correlation passes a *χ*^2^ statistical test, at a significance level *α_N_* which reflects the sample size *N*. We compare the LD blocks discovered here to previous studies of LD in the human genome (Note S1).

#### LD Blocks Encompass Correlations between Alleles

To delineate the LD blocks, we first downloaded phased genotype data from Phase 3 of the 1000 Genomes Project (1KGP; The 1000 Genomes Project Consortium, 2015). This release contains 2,504 individuals from five continental super populations: African (AFR), East Asian (EAS), European (EUR), South Asian (SAS), and Admixed American (AMR). We computed the LD partition in each super population separately, by applying the graph clustering algorithm to the sample correlations measured in each ancestry. We summarize the results from the EUR population (Figure 1), which is the ancestry of the GWAS cohorts we analyzed in this work. We also provide corresponding results for the other unadmixed super populations (Figures S2 – S5).

**Figure 1.**
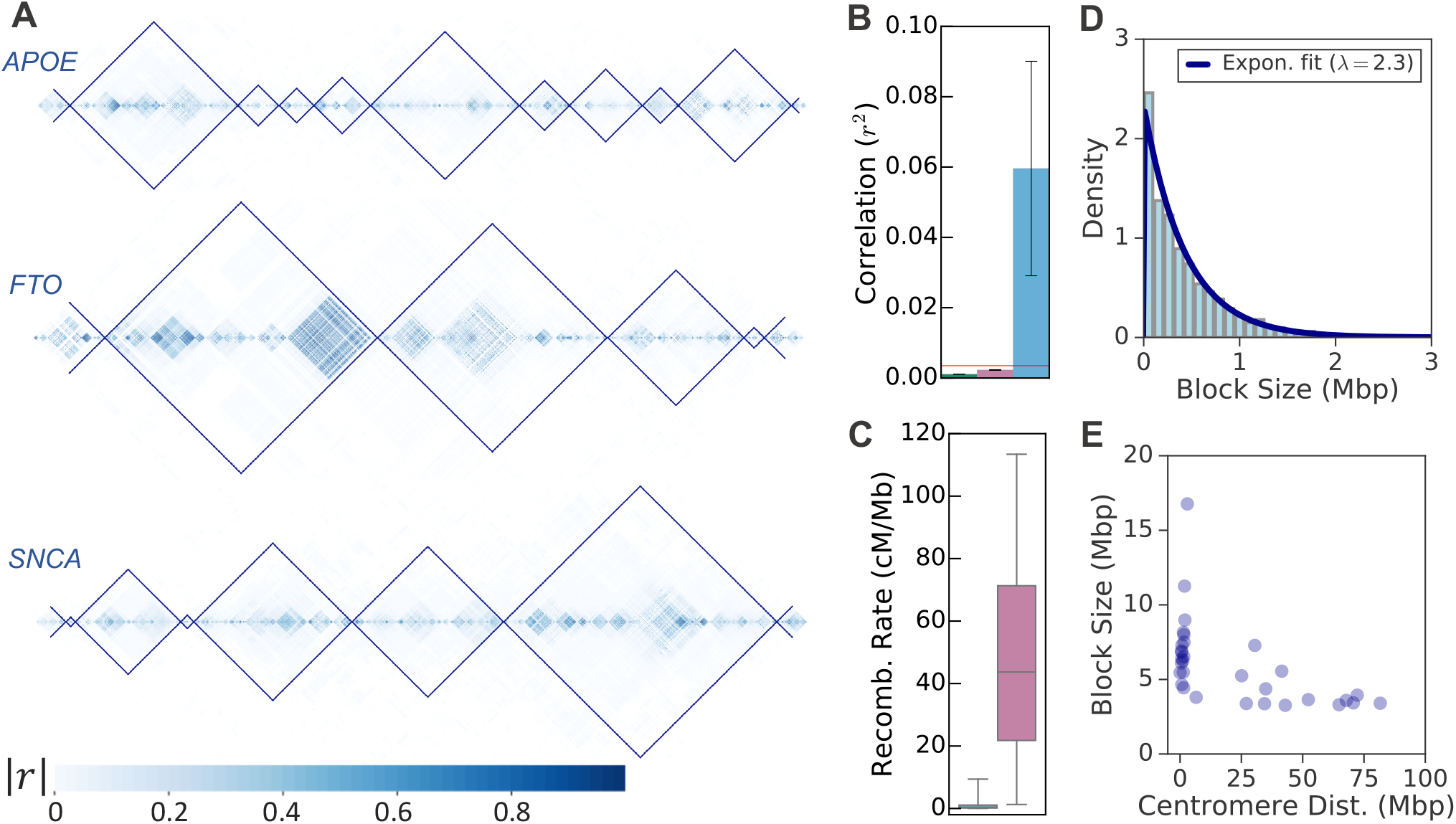
The LD partition in Europeans. (A) Heatmaps visualize the magnitude along the diagonal of the correlation matrix computed from the 1KGP EUR super population. (The matrix has been rotated by 45° counterclockwise, so that the diagonal runs horizontally.) Each heatmap is subset to a prominent locus in human genetics: *APOE* (chr19:44,000,000–48,000,000), *FTO* (chr16:52,500,000–54,500,000), and *SNCA* (chr4:89,000,000–95,000,000). (Coordinates are with respect to GRCh37.) Thin lines delineate the LD blocks in this population. The LD boundaries trace the edges of the block diagonal structure of the underlying correlation matrix. Regions of strong correlation are never split across blocks, which encompass both strong and weak correlations. (B) The mean squared correlation (± standard deviation) between pairs of variants within the same block (sky blue), in adjacent blocks (reddish-purple), or in non-adjacent blocks (green, barely visible). The red line indicates the minimum correlation between two variants we consider to be in LD, 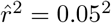, after correcting for bias in the sample correlation coefficient. (C) Box plot of the recombination rates at the boundaries between LD blocks (right, reddish-purple) and within blocks (left, sky blue, barely visible). The boxes span the interquartile range (middle 50%) of each distribution of recombination rates; the whiskers extend to the 5th and 95th percentiles. Recombination rates are markedly higher at the boundaries between LD blocks compared to within blocks. (D) The distribution of LD block sizes in Europeans follows an exponential with rate parameter λ = 2.3, consistent with a random (Poisson-distributed) occurrence of recombination hotspots throughout the genome. Block sizes in the top 0.5% have been removed. (E) The block sizes trimmed from panel D, as a function of their distance to the centromere. The largest blocks tend to reside near the centromere, where recombination is repressed (Nambiar and Smith, 2016).

The spatial pattern of LD in each super population is described by a correlation matrix (Figure S1). Our LD blocks hew to the block diagonal structure of this matrix, delineating the discrete regions of high correlation that occur along its diagonal (Figure 1A). Across loci, regions of high correlation are wholly contained within blocks; we observe little correlation between blocks. We computed the mean and standard deviation of the sample correlation, *r*^2^, between variants in the same block; between variants in neighboring blocks; and between long-range variants in nonadjacent blocks (Figures 1B and S2A). As expected, the average correlation within blocks is substantially larger than between neighboring or distant blocks. Moreover, in all populations, the mean correlation across blocks is less than the minimum correlation we consider to be non-negligible, by more than two standard deviations.

#### LD Blocks Reflect Molecular Mechanisms and Historical Demographics

The LD blocks are further validated by their consistency with the molecular mechanisms of recombination, and with human demographics. For instance, recombination rates are dramatically elevated at the boundaries between blocks, indicating that block endpoints coincide with recombination hotspots (Figures 1C and S2B). The consensus hotspot motif CCNC-CNTNNCCNC occurs at the same rate within the LD boundaries as it does in recombination hotspots (Table S2; Myers et al., 2008). Further still, the distribution of LD block sizes is well-fit by the exponential distribution after removing large outliers (Figures 1D and S3A), consistent with a random (Poisson) distribution of hotspot locations throughout the genome. The largest LD blocks, which deviate from the exponential distribution, tend to occur near a centromere (Figures 1E and S3B), where recombination is known to be strongly repressed (Nambiar and Smith, 2016). Finally, we find that the LD blocks in the AMR super population are substantially larger than in the unadmixed populations (Table S1). The large AMR blocks presumably reflect the recent history of admixture in AMR (The 1000 Genomes Project Consortium, 2015) and the very long-range ‘admixture LD’ it should induce (Nei and Li, 1973; Loh et al., 2013). (We validated that the LD block sizes reflect the true LD in the underlying population, not the sample size of each 1KGP cohort. See Figure S4 and Table S3.)

#### Universal LD Blocks

Population-specific LD blocks enable the genome-wide application of the fine-mapping model to cohorts of homogeneous ancestry. In order to analyze multi-ancestry GWAS data in the same way, we require a partition of the genome into universal LD blocks that are statistically uncorrelated in all populations under study. We therefore developed an algorithm (see the *Methods*) to identify common LD block boundaries that are shared across super populations, at a resolution of 2 kbp, which is the approximate width of recombination hotspots (Wall and Pritchard, 2003; Paigen and Petkov, 2010). Applying this method to the four unadmixed super populations (AFR, EAS, EUR, and SAS), we discover the existence of a universally valid LD partition, in the sense that the universal blocks span the genome and are mutually uncorrelated in all four populations (Figure S5). We found a total of 2,326 such blocks across all chromosomes, with a median length of ~ 600 kbp (Table S4). The existence of a universal LD partition holds even when the AMR super population is included in the consensus, although the long-range LD in AMR necessitates a coarser partition.

### The Variant-to-Gene Map

The LD blocks enable a divide-and-conquer approach to compute the probability that each variant in the human genome is causal. In order to compute the probability of association for all *genes*, we additionally require a mapping from putative causal variants to the target genes they affect. We built such a mapping from the union of three data sources: 1.) the Ensembl Variant Effect Predictor (VEP; McLaren et al., 2016); 2.) the Genotype-Tissue Expression (GTEx) project (GTEx Consortium, 2020); and 3.) a comprehensive map of *cis*-regulatory elements (promoters and enhancers) to their target genes. First, we used the functional consequences from the VEP to link variants in untranslated regions (UTRs), splice sites, or exons to the corresponding gene (see the *Methods*). Second, GTEx has pinpointed variants that mediate the expression of specific genes in particular tissues; we include these variant-to-gene associations in the map. Lastly, we built machine learning (ML) models of enhancer and promoter activity. The enhancer-promoter interactions we infer from these models constitute the majority (88%) of the variant-to-gene links in the map (see below).

#### Machine Learning of Enhancer-Promoter Interactions

Consider a DNA element in open chromatin (i.e., a DNase-I hypersensitive site, DHS), comprising a potential *cis*-regulatory element (Andersson et al., 2014). If that element is an enhancer for a particular promoter, then its activity (chromatin accessibility) should predict the promoter activity, and it should at least occasionally come into physical proximity with the promoter (see the *Methods* for the justification of these propositions). Therefore, it should be possible to identify the enhancers for a promoter, by finding the physically interacting regulatory elements that best predict the promoter activity. To test this hypothesis, we built a ML model for each promoter in the genome, to predict the promoter activity across cell types, as a function of the activity of nearby regulatory elements (Figure S6; see the *Methods*). The models identify small sets of predictive enhancers for each promoter, subject to the constraint that they contact the promoter in three-dimensional space. We infer that the regulatory elements selected by these models are enhancers for the corresponding promoter. In turn, genetic variants that fall within (or very near) an enhancer are linked to that enhancer’s target gene(s).

#### Machine Learning Models Identify Enhancers

To build the ML model for each promoter, we first identify a set of putative enhancers within 2Mbp. We reasoned that functional enhancers should exhibit coordinated activity with other nearby enhancers and promoters. If so, then it should be possible to accurately predict the activity (accessibility) of the *enhancers* themselves, on the basis of nearby regulatory elements. To test our hypothesis, we built models to predict the activity of each DHS (potential regulatory element; see the *Methods*). We measured the predictive power of the model for each DHS (given by its coefficient of determination, *R*^2^), and we tested whether DHSs that are better predicted by nearby open chromatin are more likely to overlap known or suspected enhancers. To independently assess whether a DHS overlaps a true enhancer, we first considered histone modifications thought to mark enhancers (The ENCODE Project Consortium et al., 2020). Indeed, as the DHS predictability (*R*^2^) increases, we find that DHSs are enriched in chromatin states associated with enhancer activity (Figure S7). Above an *R*^2^ threshold of ~0.7, DHSs are enriched for all the enhancer states identified by genomic segmentation algorithms on the basis of histone marks (Ernst and Kellis, 2012, 2017; Boix et al., 2021). We also considered overlap with active enhancers (enhancer RNAs, eRNAs; Lam et al., 2014; Core et al., 2014; Tippens et al., 2020) identified by the FANTOM5 consortium (Andersson et al., 2014), or with causal variants mediating gene expression and GWAS traits (see below). These features are more direct indicators of functional enhancer activity than statistically inferred chromatin states. Notably, DHSs preferentially overlap FANTOM5 enhancers and causal variants only after *R*^2^ exceeds ~0.8-ol, suggesting that strong evidence for enhancer activity does not emerge until a higher *R*^2^ threshold. Nevertheless, to ensure we capture all possible enhancers, we considered any DHS within 2Mbp of a promoter to be a putative enhancer if its *R*^2^ exceeds the less stringent 0.7 threshold.

#### Machine Learning Models Accurately Predict Promoter Activity

Given the putative enhancers for each promoter, we next built and evaluated the *promoter models*. Each promoter model is intended to identify a subset of its potential *cis*-regulatory elements that are true enhancers for the promoter. If the enhancers actually selected by each promoter model are correct, then the models should accurately predict promoter activity on out-of-sample data (which were not used to train the models), on the basis of these enhancers. To validate the promoter models, we therefore computed their coefficient of determination 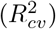, as measured on out-of-sample (three-fold cross-validation) data (Figure 2A). The distribution of 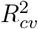 over all the promoter models is left-skewed, with a mode at 74%, indicating that 74% of the variation in promoter accessibility across cell types can typically be predicted from enhancers. (Note that this is likely an underestimate of the true performance on out-of-sample data, because the data folds are large, and some cell types are represented only in the validation fold. See the *Methods*.) A promoter for the *MYC* gene exemplifies the predictive performance of these models (Figure 2B). The promoter’s activity is accurately predicted over the full range of chromatin accessibility and across the gamut of human cell types assayed for open chromatin, including an outlier observation with extremely high promoter activity. The variant-to-gene map incorporates predicted enhancers exclusively from the top 80% of promoters, ordered by predictive performance on out-of-sample data, thereby including promoters with 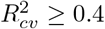.

**Figure 2.**
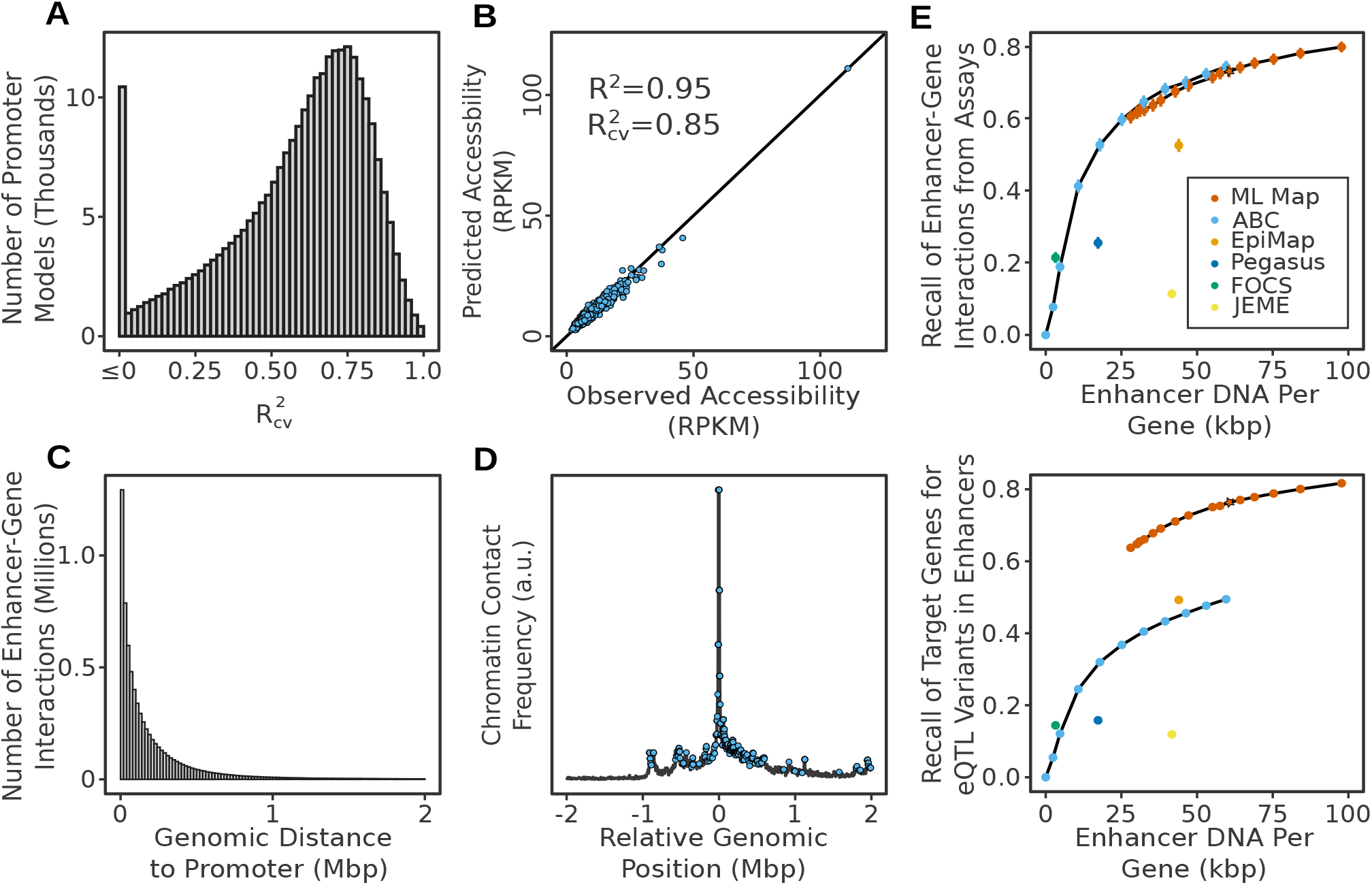
Enhancer-to-gene map links non-coding DNA to target genes. (A) Out-of-sample predictive power for the ML models of chromatin accessibility at the promoter. Histogram of 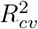 (percentage of variance in promoter accessibility explained by the model on out-of-sample cross-validation folds), over all promoter models. The models are based on chromatin accessibility at enhancers within 2Mbp of the promoter. (B) The predictive performance of a single model, for a promoter of the *MYC* gene. Predicted versus observed chromatin accessibility at the *MYC* promoter, across hundreds of ENCODE samples, measured in RPKM from open chromatin sequencing assays (see the *Methods*). (C) Distribution of the number of predicted enhancer-to-gene interactions, as a function of distance between the enhancer and promoter. (D) Normalized contact frequency between a *MYC* promoter and each candidate enhancer DHS within 2 Mbp. Blue points indicate *MYC* enhancers selected by the promoter model. (E) The validation rate of our enhancer-to-gene map (ML Map) and alternatives against: (top) a compendium of enhancer-gene interactions (primarily from high-throughput CRISPR screens) that have been experimentally validated; and (bottom) a set of causal variants mediating gene expression (eQTLs) that reside within 300 bp of likely enhancers (see Figure S9 for the validation rate on all eQTLs). Recall is plotted as a function of map density, i.e., the average length of DNA predicted to regulate expression of a given target gene. Error bars show standard error of the mean of 735 (top) or 4882 (bottom) validated enhancer-gene pairs. The star denotes the density at which we configured our variant-to-gene map.

#### Enhancer-Promoter Interactions Reflect Chromatin Looping

If the ML models capture the true enhancers for each promoter, then the distances between predicted enhancer-promoter pairs should be consistent with the three-dimensional organization of chromatin, which is the very scaffold on which these interactions occur. Distal *cis*-regulatory interactions are thought to be mediated by chromatin loops that bring genomic sites into close physical proximity. We therefore incorporated experimentally observed contact frequencies between promoters and candidate enhancers as prior information in the ML models. Consequently, we confirmed that the resulting distribution of enhancer-promoter distances matches the experimentally observed contact frequencies. As expected, we find that the number of enhancer-promoter interactions decays as a function of the enhancer distance from the promoter, at a rate which is similar to the decay of chromatin contact frequencies with distance (Figure 2C). Visualizing the density of predicted regulatory interactions on a logarithmic scale (Figure S8), we find a discontinuity in the distribution, which likely reflects the enrichment of regulatory interactions within the same topologically associated domain (TAD; Szabo et al., 2019). We observe two regimes, whereby more proximal interactions (<200 kbp) decay with a power law exponent of −0.64, whereas more distal interactions (>500kbp) decay with an exponent of −2.3. We note that the proximal decay rate nearly matches the prediction from an extrusion globule model of intra-TAD chromatin interactions (exponent = −0.7; Sanborna et al., 2015).

Because chromatin contacts decay rapidly with genomic distance, a common heuristic is to link enhancers to their nearest protein-coding gene. Indeed, we observe that 53% of enhancers are predicted to regulate the expression of their nearest gene. However, these interactions account for only 18% of the total predicted regulatory interactions in the genome (and only 27% of the interactions affecting protein-coding genes). If we were to link each enhancer only to its nearest gene, we would overlook the majority of *cis*-regulatory interactions and potential disease genes.

At the level of individual promoters, we find that the genomic distribution of predicted enhancers matches the local pattern of chromatin looping (Figure 2D). Considering our previous example of the *MYC* promoter, we find that chromatin loops bring specific enhancers, up to 2 Mbp away from the gene, into proximity with the promoter, and that these particular enhancers are identified as predictive by the ML model. (The distal *MYC* enhancers are among the longest-range interactions in the human genome that have been experimentally validated.)

#### Validation on Known Variant-to-Gene Interactions

Finally, the strongest test of the ML models is to determine how well they predict experimentally observed enhancer-gene interactions, as a function of their density (the number of enhancers linked to each promoter). We therefore compiled a database of functional genomics assays that measure enhancer-gene interactions using a range of techniques (File S1), resulting in 735 unique enhancer-gene pairs. Most of these derive from high-throughput CRISPR screens in the K562 cell line. We computed the proportion of known interactions recovered (recalled) by the ML models. Since the density of the models can be tuned with a single parameter, we tested the recall of the resulting map across a range of densities (Figure 2E), obtained by varying the tuning parameter. At the density we chose for our analyses in this work (the starred point in Figure 2E), the ML map recalls 73% of known enhancer-gene interactions. Next, we evaluated our enhancer-to-gene map on expression quantitative trait loci (eQTLs) discovered by the GTEx Consortium GTEx Consortium (2020). We considered only eQTLs that GTEx fine-mapped to single causal variants in/near likely enhancers, resulting in 4882 variant-gene associations (File S2). Here, we recover 76% of interactions. (See the *Methods* for details.) Note that we built the ML models from open chromatin profiles and chromatin contact frequencies (see the *Methods*). The validation data, on the other hand, are from different modalities (e.g., CRISPR screens and human genetic variation) and different distributions of human cell types/tissues.

For comparison, we evaluated the performance of several alternative methods from the literature, which also link enhancers to target genes: FOCS (Hait et al., 2018), JEME (Cao et al., 2017), PEGASUS (Clément et al., 2020), EpiMap (Boix et al., 2021), and ABC (Nasser et al., 2021). We find that the map introduced here significantly outperforms all of the alternative methods at predicting experimentally discovered enhancer-gene interactions, even at the same density as the other methods (Figure 2E). While the recently published ABC method matches the performance of the ML map on functional genomics assays, its performance drops on the GTEx eQTLs, which are more representative of the GWAS variants mediating disease risk.

We attribute the relative advantage of the ML map versus ABC, on eQTL data, to a difference in the way enhancers are modeled. Both methods incorporate similar chromatin conformation and accessibility information. However, the ML map discovers predictive relationships between enhancers and promoters, learned simultaneously from a large compendium of diverse cell types. ABC, on the other hand, bases its predictions largely on the strength of an enhancer’s accessibility in each cell type separately. We find that ABC performs exceptionally well at predicting enhancer-promoter interactions in the specific cell types used to build the map (e.g., the K562 cell line, in which most of the CRISPR studies were carried out). But its performance drops when considering arbitrary cell types (as in the bulk tissues from GTEx), because many enhancers that operate in these cells do not have high DHS activity in the cell types profiled by the ENCODE consortium (The ENCODE Project Consortium et al., 2020; Figure S10). We investigated the performance of the ML map and ABC (both tuned to similar enhancer density), stratified by the maximum accessibility of a validated GTEx enhancer across ENCODE cell types (Figure S10). We observe, first, that the functional genomics assays tend to discover enhancers with greater DHS activity (in ENCODE cell types) than the interactions observed in the eQTL studies. Second, we find that the ABC model excels on such high-activity enhancers, whereas the ML map maintains high performance across a wide range of enhancer activities.

### The Statistical Fine-mapping Model

Equipped with a partition of the genome into LD blocks, and a comprehensive map of variants to target genes, we extended the equations of statistical fine-mapping to compute the posterior probability of association (PPA) for all genes. (We explain the approach schematically in Figure S12 and formally in the *Methods*.) Our goal is to compute the probability that any particular gene is targeted by a causal variant. We therefore consider all possible combinations of causal variants that could plausibly explain the genetic architecture of a trait. We evaluate the probability of each combination, and sum the probabilities of all combinations that include a causal variant linked to the gene. As we show below, using this approach, it is possible to confidently identify genes targeted by a causal variant, without actually ascertaining the causal variants!

#### Incorporating Prior Knowledge of Variant Consequences

To compute the probability of any combination of causal variants requires two sources of knowledge: the GWAS data, and our prior beliefs. In particular, we must specify our prior belief (encoded as a prior probability) that any particular variant is causal for a trait, before the variant is actually tested for association. We reasoned that the prior probability that any variant is causal should depend upon its biochemical consequence: namely, whether, and how, the variant might alter the function or expression of a gene. These probabilities are difficult to deduce from first principles, but we can *learn* them directly from GWAS data. We begin by annotating each variant in the genome according to its most severe biochemical consequence (e.g., whether a variant falls into an active enhancer or a splice site), combined with an indicator denoting whether each variant resides in conserved DNA (Table S5). To categorize variants by their most severe consequence, we leverage the variant-to-gene map. Then, we apply the expectation maximization (EM) algorithm to a compendium of GWA studies, to learn the prior probability of association for variants in each biochemical category (see the *Methods*). The EM algorithm discovers the priors that maximize the probability of obtaining the GWAS data we actually observed. We refer to the probabilities obtained this way as the ‘biochemical prior’.

#### Validation of Statistical Fine-mapping

In order to validate the statistical fine-mapping calculations, we first compute the probability that each variant in the genome is causal—i.e., the posterior probability of association (PPA) for variants. For any variant, the PPA is obtained by summing the probability of all plausible combinations of causal polymorphisms that include the variant. Note that this is a summation over the *same* probability distribution we use to compute gene probabilities. Variant and gene PPAs are just sums (marginal distributions) over different combinations of causal variants, using exactly the same statistical model. Therefore, we can use variant probabilities to validate the core statistical computations underlying our gene probabilities.

Here, we may leverage our variant annotations once more. If our calculations are correct, then the probability that a variant is causal should depend on its biochemical consequence, even before we inject prior knowledge into the model. GWAS signals are induced by causal variants in exons, UTRs, and enhancers, for example. The statistical structure of the data alone should implicate these variants as most likely to be causal, because *their correlations* with neighboring variants are the ones that create the observed pattern of GWAS signals. To test whether the model correctly unravels the tangles of GWAS signals, without the aid of our prior knowledge, we built a simpler version of the model. This formulation uses a ‘uniform prior’ in which all variants are presumed equally likely to be causal, *a priori*. We applied this simpler model to a compendium of GWA studies, stratified all the variants by their causal probability in each GWAS, and tallied the variant consequences in each PPA bin (Figure 3A, left). As expected, we observe a linear relationship between the probability that a variant is causal and its likelihood to target plausibly functional DNA. The more probable variants are to be causal, the more likely they fall into exons, UTRs, enhancers, or other functional DNA elements that must be the source of GWAS associations. Then, when we encode variant annotations into the full model with the ‘biochemical prior’, nearly all potentially causal variants have a plausible mechanism of action (MOA; Figure 3A, right). Almost 90% of potentially causal variants computed by the full model can be linked to a gene by our variant-to-gene map. Moreover, the full biochemical model uncovers the relative frequencies of different types of variants among causal GWAS polymorphisms. Recapitulating previous results (Maurano et al., 2012), we find that most causal GWAS variants target promoters and enhancers (orange and blue, respectively, in Figure 3A).

**Figure 3.**
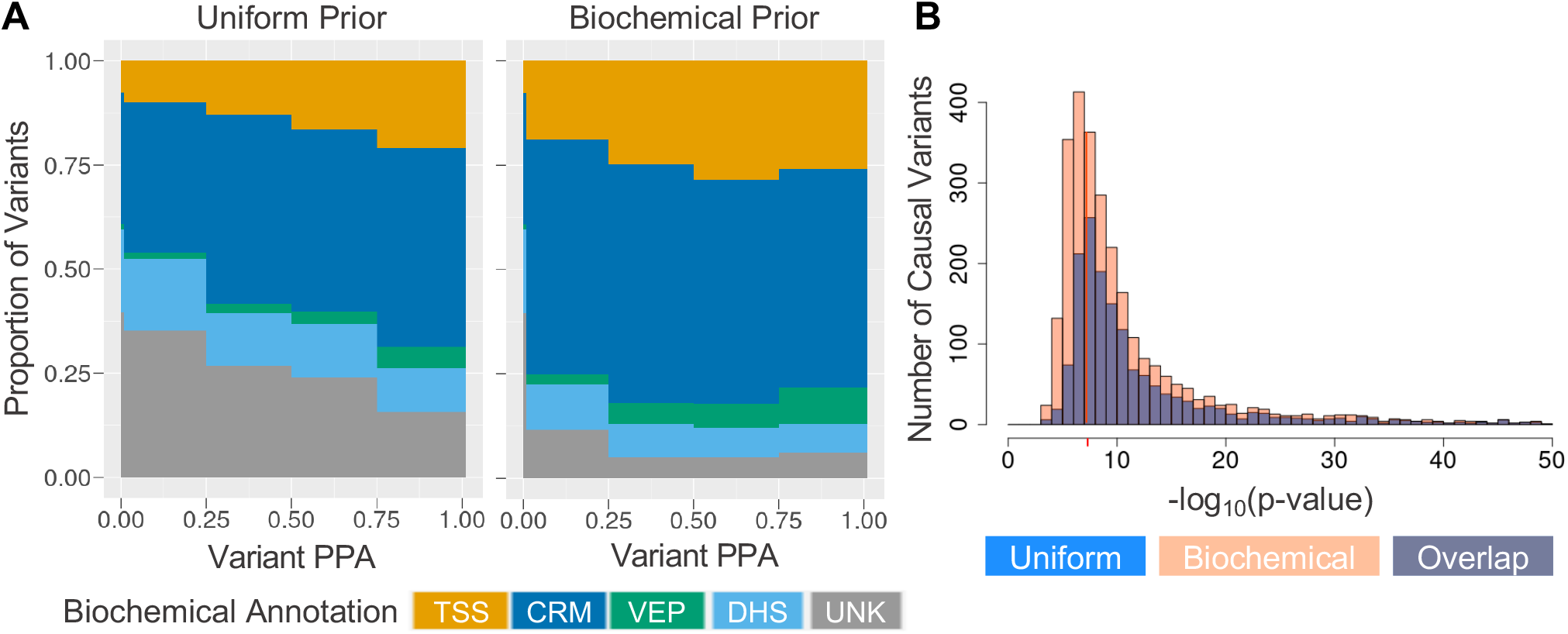
Statistical fine-mapping of causal variants. (A) Predicted biochemical consequences of variants, as a function of their posterior probability of association (PPA) to the traits in our GWAS compendium. Variant PPAs were computed either with the uniform (left) or biochemical prior (right). TSS denotes variants falling near transcriptional start sites; CRM indicates variants in/near an enhancer linked to at least one gene in the *cis*-regulatory map; VEP variants can be linked to a gene by a Variant Effect Predictor consequence; DHS represents variants in/near a DHS *not* linked to any gene by the enhancer-to-gene map; UNK denotes variants in otherwise unannotated intergenic or intronic DNA with unknown function. (B) Distribution of GWAS p-values for likely causal variants (i.e., variants with PPA ≥ 0.5), discovered from a collection of 40 GWA studies. The red line marks the threshold for genome-wide significance (5 × 10^−8^). The distribution obtained from the uniform prior is plotted in blue; the biochemical prior in transparent orange-red. The region where the two distributions overlap appears purple. Variants with a p-value less than 1 × 10^−50^ have been clipped from the right edge of the histogram, where the two distributions closely coincide.

#### Power to Detect Sub-GWS Causal Variants

We applied the fine-mapping algorithm genome-wide, with the expectation that it would uncover causal variants even if their GWAS p-values do not meet the stringent GWS threshold. To validate the power of the method to detect causal variants across the full range of GWAS signals, we collected all likely causal variants (PPA > 0.5) across our GWAS compendium, and plotted the distribution of their GWAS p-values (Figure 3B). The peak of the distribution occurs near the GWS threshold; a substantial proportion of likely causal variants can be identified below it. Moreover, incorporating variant annotations in the prior doubles the number of likely causal variants that can be detected. The advantage is most profound near the GWS threshold, as expected: prior knowledge has a larger impact when the data alone are indeterminate.

A small fraction of causal variants are apparently detected even from very weak association signals, with p-values greater than 10^−3^ (not shown). In theory, that result could be obtained, for instance, from positively correlated causal alleles that mediate opposing, counterbalancing effects. Nevertheless, seemingly causal variants emerging from the weakest trait associations are often spurious. They result from mismatches between the LD matrix in the GWAS cohort and our reference population from the UK Biobank, even after nugget regularization of the reference matrix (see the *Methods*). Conservatively, we generally reject such variants from consideration as potentially causal.

#### Distribution of Gene Probabilities

Given a sufficiently powered GWAS of a polygenic trait, we expect the distribution of gene probabilities to follow a bimodal distribution, with asymmetric peaks at its extremes. Genes are either associated with the trait, or they are not, and the latter proportion should be the dominant one. This is precisely what we observe. Consider, for instance, the probability distribution for protein-coding genes across four GWA studies of body weight traits (Figure 4A; the distribution is similar for each GWAS individually). The dominant peak at zero comprises genes with virtually no evidence of association to body mass index (BMI) or body fat percentage/distribution; the smaller peak at unity consists of genes that we predict, with near certainty, are targeted by a causal variant for these traits. Most of the probability mass falls on the left side of the distribution (PPA < 0.5), where genes are unlikely to be targeted by a causal variant. The probability mass between the peaks reflects uncertainty in our identification of trait-associated genes (i.e., genes targeted by a causal variant), which arises from uncertainty about which variants are causal. Theoretically, as the GWAS sample size increases, the probability mass in the middle of the distribution should progressively retreat into the peaks at the extremes. Encoding prior knowledge in the statistical model shifts the probability mass from the left to the right of the distribution, approximately doubling the number of high-confidence trait-associated genes. Across a collection of diverse traits, incorporating prior knowledge yields a greater probability for ~ 90% of plausible disease genes, with many undergoing substantial shifts in probability relative to an uninformed model (Figure 4B).

**Figure 4.**
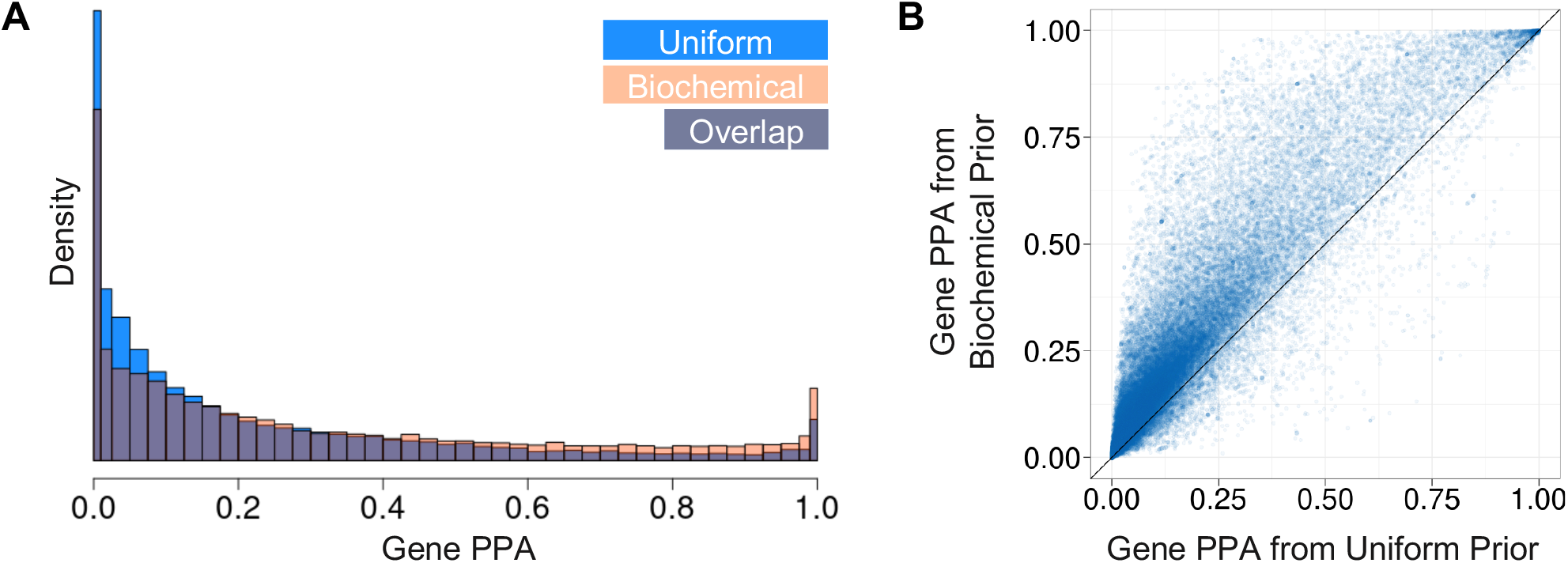
Statistical fine-mapping of trait-associated genes. (A) Distribution of the PPA for all protein-coding genes, taking the maximum PPA for each gene over two GWA studies of BMI (Canela-Xandri et al., 2018; Neale et al., 2021), a study of body fat percentage (Neale et al., 2021), and a study of waist-to-hip ratio after adjustment for BMI (Pulit et al., 2019). Gene PPAs computed under the uniform prior are plotted in blue. PPAs under the biochemical prior are overlaid in transparent orange-red. The region of overlap between the two distributions appears purple. (B) A density scatter plot showing the difference in gene PPAs computed by the biochemical vs. uniform prior, for protein-coding genes across a compendium of distinct traits.

#### Gene Clusters

We emphasize that a single causal variant may target multiple genes, as when a variant disrupts an enhancer that regulates the expression of several genes. Thus, not all genes targeted by a causal variant (i.e., genes with a high PPA) are necessarily causal themselves. In disease, we presume that the MOA of a causal variant involves the dysregulation of at least one gene, but the perturbation of other genes targeted by the same variant may be innocuous. It remains necessary to prioritize candidate causal genes from among those that are targeted by the same causal variant(s) (e.g., see Figure S13A). We therefore clustered all plausibly trait-associated genes (e.g., all genes with PPA ≥ 0.1) into groups of genes implicated by the same variants (see the *Methods* and Figure S13A). Considering all such gene clusters across the traits we analyzed, most clusters are small: the median cluster size is only 1 gene (Figure S13B). The distribution is right-skewed, with a mean of 2.7 genes and a standard deviation of 3.1 genes.

#### Power to Detect Trait-Associated Genes

In order to measure the sensitivity of our method to discover genes dysregulated by causal variants, we first computed the total number of causal variants whose existence can be detected in the GWAS. Formally, we calculated the expected value of the number of causal variants in the genome, considering only putative causal variants with a GWAS p-value less than a threshold (e.g., 10^−3^; see the *Methods*). This calculation counts all detectable causal variants—subject to the p-value constraint—regardless of whether they can be fine-mapped; it represents the maximum number of independent association signals that could be linked to genes. We then compared the number of trait-associated genes and gene clusters, as well as the number of fine-mapped variants, to this upper bound (Figure 5). We consider genes with PPA > 0.5 to be trait-associated, and variants with PPA > 0.5 to be likely causal (i.e., fine-mapped). Trait-associated gene clusters contain at least one trait-associated gene. Remarkably, across a representative sample of traits, the statistical framework generally identifies more trait-associated gene clusters than there are GWS causal variants in the GWAS data. This striking sensitivity results from the ability of the platform to extract gene-trait associations from across the genome, including loci with association signals that fall below the stringent genome-wide p-value threshold. Moreover, the number of trait-associated gene clusters exceeds the number of causal variants that can be fine-mapped, by severalfold. Many trait-associated genes integrate their PPA over multiple potentially causal variants. In order to implicate gene(s) in a trait-associated locus, it is often unnecessary to pinpoint the causal variant(s), because different candidate causal variants often implicate the same gene(s).

**Figure 5.**
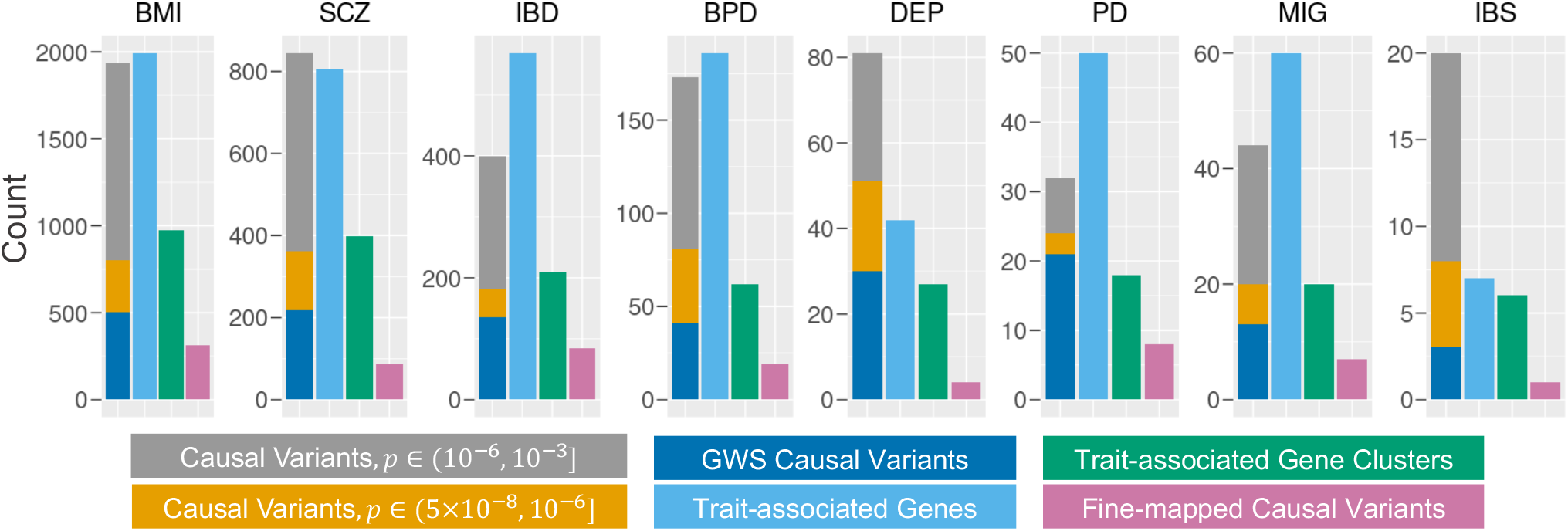
Identification of trait-associated genes and gene clusters. The number of trait-associated genes and gene clusters discovered from 8 representative traits, compared to the total number of causal variants and the number of fine-mapped variants in each trait. All probabilities were computed with the biochemical prior. Traits: BMI, body mass index (Neale et al., 2021); SCZ, schizophrenia (Psychiatric Genomics Consortium, 2021); IBD, inflammatory bowel disease (de Lange et al., 2017); BPD, bipolar disorder (Mullins et al., 2021); DEP, depression (Howard et al., 2019); PD, Parkinson’s disease (Nalls et al., 2019); MIG, migraine (Canela-Xandri et al., 2018); IBS, irritable bowel syndrome (Eijsbouts et al., 2021). For each trait, the leftmost bar indicates the expected value of the number of causal variants (i.e., the total number of independent detectable signals in the GWAS), considering only putative causal variants with a GWAS p-value smaller than 10^−3^. The total number of causal variants is partitioned into 3 bins according to the p-values of each variant. GWS (genome-wide significant) causal variants have p-values ≤ 5 × 10^−8^. Trait-associated genes are those with PPA ≥ 0.5; similarly, trait-associated gene clusters contain at least one gene with PPA ≥ 0.5. Likewise, fine-mapped causal variants have PPA ≥ 0.5.

The level of success achieved by the fine-mapping framework depends on the details of its implementation. For one, the biochemical prior doubles the number of causal variants and trait-associated genes that can be fine-mapped (Figure S14). We show that a small, fundamental set of annotations, when incorporated into the model via EM, substantially improves the identification of trait-associated genes. For another, consider if we had precomputed a set of fine-mapped variants, and then linked potentially causal variants to genes in a subsequent step. We would have limited our discoveries to genes targeted by a fine-mapped variant, diminishing the number of recovered disease genes.

#### Validation on Well-known Disease Genes

Next, we validated that the platform rediscovers known genedisease associations. We considered inflammatory bowel disease (IBD) as an exemplar of a well-studied polygenic disease, in which many risk-modifying genes have already been established. As in Nasser et al. (2021), we obtained a set of 83 known IBD genes from a recent review (Graham and Xavier, 2020). To measure the performance of the platform against known sets of disease genes, we must first choose a probability threshold beyond which we consider a gene to be implicated. Naturally, we chose a gene PPA threshold of 0.5 (i.e., 50%), beyond which genes are more likely than not to be targeted by a causal variant. We obtained GWAS data on IBD, which comprises the disease subtypes Crohn’s disease (CD) and ulcerative colitis (UC), from de Lange et al. (2017). We analyzed three GWA studies, on a CD cohort, an UC cohort, and a combined IBD cohort (consisting of both CD and UC cases). Pooling the genes implicated by the platform across those studies, we recall 70% of the 83 IBD genes. Our platform can only detect disease genes targeted by variants that induce GWAS signals. We therefore considered a subset of the 83 IBD genes, consisting of 26 genes within 1 Mbp of a GWS association signal in noncoding DNA, as in Nasser et al. (2021). These are known IBD genes that could be plausibly targeted by a causal variant in the best available GWAS data. On that subset, the platform recalls 81% of the genes.

We compared this performance to alternative methods for identifying genes targeted by causal variants; namely, Summary-based Mendelian Randomization (SMR; Zhu et al., 2016), Coloc (Giambartolomei et al., 2014), and the ABC method introduced in Nasser et al. (2021). The approach described here recalls about twice as many known IBD genes as the previous algorithms (Table 1). Strikingly, our recall nominally exceeds even Summary-MultiXcan (Barbeira et al., 2018), a transcriptome-wide association study (TWAS) method. S-MultiXcan integrates GWAS signals genome-wide with the full complement of GTEx eQTLs across tissues, without requiring fine-mapping or making assumptions on the number of causal variants. It therefore achieves remarkable sensitivity to identify proximal genes dysregulated by nearby GWAS variants (for which GTEx has the most power), thereby accounting for many already-known disease genes. However, unlike the other methods, S-MultiXcan does not specifically identify genes that are directly targeted by a causal variant (Wainberg et al., 2019); many of the genes it associates with disease are likely to be targeted by passenger variants in LD with causal polymorphisms. In IBD, S-MultiXcan implicates ~3.5 times as many genes as the number of distinct gene clusters we identify, without any means of identifying which genes should be prioritized against each other.

**Table 1.** Comparative recall of known IBD genes. Recall of known disease genes in IBD from this work and alternative methods. Each method was evaluated on the union of genes it discovers from GWA studies of IBD, CD, or UC; see the *Methods*. ‘IBD Gene Recall’ is the percentage of well-known IBD genes rediscovered by each method, using the set of 83 genes from Graham and Xavier (2020). ‘IBD Subset Recall’ is the same, but evaluated on a subset of these genes tested in Nasser et al. (2021), which reside within 1 Mbp of GWS associations in non-coding DNA. ‘Total IBD Genes’ indicates the total number of genes associated with IBD or its subtypes, by each method. In parentheses, we note the Jaccard index between the validation genes recalled by our approach versus the alternatives (middle two columns), or between the full set of IBD-associated genes in our method versus the others (rightmost column). In brackets, we estimate the number of distinct gene clusters we identified from studies of IBD, CD, and UC; see the *Methods*.

| Method | IBD Gene Recall | IBD Subset Recall | Total IBD Genes |
| --- | --- | --- | --- |
| This work | 70% (1.00) | 81% (1.00) | 876 [334] |
| SMR | 42% (0.48) | 27% (0.27) | 1169 (0.13) |
| S-MultiXcan | 67% (0.73) | 69% (0.77) | 1190 (0.21) |
| Coloc | 30% (0.38) | 19% (0.24) | N/A |
| ABC-All | 25% (0.32) | 54% (0.59) | 378 (0.15) |
| ABC-Max | 21% (0.27) | 54% (0.59) | 58 (0.04) |

Similarly, we validated that the platform captures disease genes previously implicated by exome studies. Here we considered obesity, another polygenic disease with well-studied genetics. We combined the results from two association studies of BMI—one, a large exome array study (Turcot et al., 2018); the other, an equally large exome sequencing study (Akbari et al., 2021)—into a unique list of 27 BMI-associated genes implicated by protein-altering variants. For each gene, we computed its maximum PPA across two GWA studies of BMI (Canela-Xandri et al., 2018; Neale et al., 2021), one GWAS of body fat percentage (BFP; Neale et al., 2021), and a fourth study of waist-to-hip ratio adjusted for BMI (WHRadjBMI; Pulit et al., 2019). The platform recalls 78% (21/27) of the BMI genes (Table S8), which substantially exceeds previous methods (Table 2); the closest counterparts, SMR and S-MultiXcan, recall 48%, despite implicating more genes overall.

**Table 2.** Comparative recall of known obesity genes. Recall of known disease genes in obesity from the method introduced here and alternative approaches. Each method was tested on the combined genes it uncovers from GWA studies of BMI, body fat percentage, and waist-to-hip ratio after adjustment for BMI; see the *Methods*. ‘Obesity Gene Recall’ is the recall of each method on the BMI genes identified from protein-altering variants in Turcot et al. (2018) and Akbari et al. (2021). In parentheses: the Jaccard index between the known obesity genes recalled by our approach versus the validation genes recovered by each alternative. ‘Total Obesity Genes’ is the total number of genes each method associates with any of the three body weight traits. In parentheses: the Jaccard index between the full set of obesity genes implicated by our method versus the others. In brackets: the number of distinct gene clusters we identified from the three body weight traits; see the *Methods*.

| Method | Obesity Gene Recall | Total Obesity Genes |
| --- | --- | --- |
| This work | 78% (1.00) | 4891 [2111] |
| SMR | 48% (0.42) | 7851 (0.20) |
| S-MultiXcan | 48% (0.55) | 4969 (0.30) |
| Coloc | 19% (0.24) | N/A |
| ABC-All | 15% (0.19) | 691 (0.06) |
| ABC-Max | 4% (0.10) | 157 (0.02) |

Interestingly, one of the supposed BMI-associated genes we fail to implicate (*HIP1R*, PPA: 0.404) was actually misidentified from the exome (Figure S15). *HIP1R* (Huntington Interacting Protein 1 Related) was originally implicated by the missense variant rs34149579 (Turcot et al., 2018), which also has a GWS association to BMI in GWAS data (Neale et al., 2021). However, our fine-mapping indicates this variant (PPA = 0.001) is almost certainly not causal. (The modest PPA we compute for *HIP1R* derives from a number of variants other than the missense variant, mostly variants in regulatory DNA.) Instead, the missense variant is a passenger of two distinct causal variants flanking *HIP1R* (see Figure S15); its association to BMI is driven by its LD with those variants, not by any effect on *HIP1R*. In particular, the missense variant is correlated (Pearson correlation R = 0.504) with the lead variant in the locus, which has a stronger association signal (*p* = 1.7 × 10^−17^ versus *p* = 9.9 × 10^−13^) and is far more likely to be causal (PPA = 0.772). Our platform predicts that the lead variant targets an enhancer, which in turn regulates the expression of the GPCR *HCAR2*. While *HIP1R* has no obvious role in metabolism (Turcot et al., 2018), *HCAR2* is a well-known metabolic sensor whose activation promotes adiponectin secretion and inhibits lipolysis (Tunaru et al., 2003; Wanders et al., 2013), which are important pathways in body weight regulation. Therefore, the platform not only recovers the majority of disease genes identified from exome studies, but it avoids errors in interpretation or disease gene ascertainment that arise (rarely) from an exclusive focus on the exome.

#### Discovery of Disease Genes with Novel Genetics Evidence

Even for the best-studied complex diseases, the genes with well-validated genetic associations account for a small fraction of heritability. There must be a bevy of disease genes that remain undiscovered from GWA studies. A platform that can discover disease genes from a substantial fraction of GWAS signals should uncover many well-known genes, but it should also yield a majority of novel disease genes. We validated that our platform implicates a multitude of genes with novel genetic associations but with compelling biological rationale and strong supporting evidence from preclinical and clinical experiments.

A prime example is provided by the GPCR *GLP1R*, which we implicate with a PPA of 85.5% in a GWAS on BMI (Canela-Xandri et al., 2018) from the UK Biobank (Bycroft et al., 2018). The satiety hormone Glucagon-like peptide-1 (GLP-1), via binding to its cognate receptor, is well known to inhibit food intake and decrease body weight, in both humans and animal models of obesity (Drucker, 2021). Moreover, *GLP1R* agonists are the most efficacious non-surgical therapeutics for obesity in the general population (Wilding et al., 2021), and yet *GLP1R* had never been implicated by the human genetics of obesity. Our platform implicates *GLP1R* primarily on the basis of a missense variant whose association signal does not reach genome-wide significance (Figure 6). (Under the uniform prior, the PPA is only 36.6%.) To our knowledge, the platform is also the first to implicate the well-known satiety hormone *CCK* (99.1% PPA), its receptor *CCKBR* (84.3% PPA), and several other neuropeptide hormones and receptors with known roles in food intake regulation. Like GLP-1, cholecystokinin (CCK) has very strong preclinical validation as a regulator of appetite and food intake, and is the target of ongoing drug development for obesity (Christoffersen et al., 2020; Miller et al., 2021). There are as many as three causal variants that alter the expression of *CCK* (Figure S16), each falling in a distinct enhancer with a different pattern of chromatin accessibility in the brain, intestinal mucosa, and nerve tissue (determined from our atlas of ENCODE open chromatin profiles). The best fine-mapped of these variants (rs28350, PPA = 88.4%) targets *CCK* through a distal enhancer (110 kbp away from *CCK*, which is not the nearest gene).

**Figure 6.**
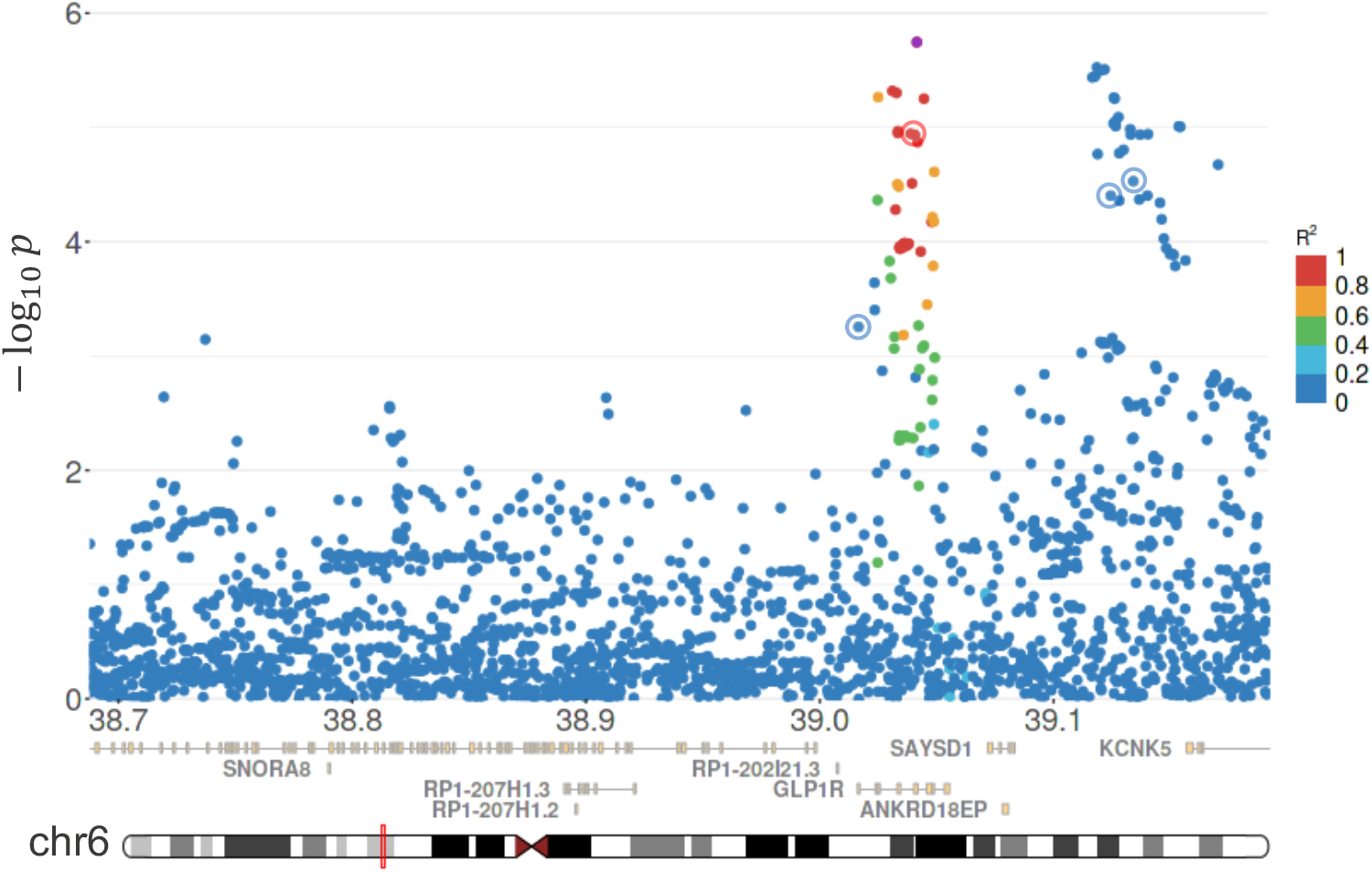
Discovery of *GLP1R* as an obesity gene. The association signal in an LD block (red box) on chromosome 6, from a GWAS on BMI (Canela-Xandri et al., 2018) in the UK Biobank (Bycroft et al., 2018). Each point is a genetic variant within the block: its nucleotide position (x-axis) is given in Mbp, relative to build GRCh37, and its association to BMI (y-axis) is expressed as the negative logarithm of its GWAS p-value, *p*. The lead variant in this locus (rs1042044, purple) is a missense variant in *GLP1R*, with a p-value of 1.79 × 10^−6^ and a PPA of 71.6%. The color scale indicates the correlation, *R*^2^, between the minor allele of rs1042044 and the minor alleles of the other variants in the block. *GLP1R*, which has a PPA of 85.5% in these data, is primarily implicated by the missense variant, but it also absorbs probability mass from other potential, but less likely, causal variants. Four of these, with PPAs ranging from 3.6% to 5.5% are circled. Three are variants in cis-regulatory elements (blue) linked to *GLP1R;* the fourth is a synonymous variant (red) in *GLP1R*.

As another example, consider the toll-like receptors (TLRs) in IBD. TLRs are fundamental determinants of pathogen recognition and immune activation (both innate and adaptive immunity; Fitzgerald and Kagan, 2020). They mediate homeostasis of the host-microbe interface in the gastrointestinal tract, which is the very root of IBD pathology (Graham and Xavier, 2020). Moreover, TLRs regulate the secretion of TNF-*α*, IL-10, TGF-*β*, IL-1, and IL-18 (Fitzgerald and Kagan, 2020), all of which are known drug targets and/or well-established IBD genes (Graham and Xavier, 2020). In particular, *TLR1* coordinates the innate and adaptive immune responses to infection in the intestinal epithelium, by inducing the production of *CCL20* (Sugiura et al., 2013), another well-established IBD gene (Graham and Xavier, 2020). *TLR1* deficiency exacerbates inflammation and intestinal injury following infection in an IBD model (Choteau et al., 2017), and it can result in chronic inflammation even after pathogen clearance (Kamdar et al., 2016). Nevertheless, the TLRs, and *TLR1* in particular, have not been strongly implicated in the human genetics of IBD. Applying our framework to a GWAS meta-analysis of IBD (de Lange et al., 2017), we compute that the PPA for *TLR1* in IBD is 89.0%, with most of the association stemming from an enhancer variant more than 667 kbp away from the gene (Figure S17). Cross-referencing the distal enhancer with our compendium of open chromatin profiles from ENCODE, we find its strongest DHS signal in natural killer (NK) cells, glomerular endothelial cells, and B cells. NK and B cells are both known to express *TLR1* (Noh et al., 2020; Browne, 2012) and have been implicated in the pathogenesis of IBD (Ginsburg et al., 1983; Hall et al., 2013; Poggi et al., 2019; Bittencourt et al., 2021; Graham and Xavier, 2020).

Recently, *PPIF*, a regulator of the mitochondrial permeability transition pore, was proposed as an IBD gene, based on a single variant with a 19% PPA and functional validation that the encompassing enhancer alters *PPIF* expression and mitochondrial membrane potential (Nasser et al., 2021). Our platform computes a 80.5% PPA for *PPIF* in Crohn’s Disease (52.8% in IBD), by integrating over the probability mass of multiple potential causal variants in the same association signal analyzed by Nasser et al. (2021). While none of the individual variants has a PPA greater than 25%, they all target *PPIF*-linked enhancers with strong DHS activity in immune cells and tissues, including T cells, B cells, macrophages, monocytes, natural killer cells, the spleen, and the Peyer’s patches in the gastrointestinal tract.

A final example demonstrates the power of the platform for novel drug discovery. From a GWAS of depression (Howard et al., 2019), the top gene implicated by the platform is *NRDC* (nardilysin convertase, PPA = 96.5%), which is the only gene linked to a genome-wide association signal near its 5’ untranslated region (Figure S18). *NRDC* (*NRD1*) is an endopeptidase that cleaves endogenous opioids from the prodynorphin gene (*PDYN*), including dynorphin A, dynorphin B (rimorphin), and *α*-neoendorphin (Chesneau et al., 1994). *NRDC* transforms all of these into dynorphin A-(1-6), also known as Leu-enkephalin-Arg, thereby shifting the balance of receptor agonism from the *κ*-opioid receptor *OPRK1* to the *δ*-opioid receptor *OPRD1* (Chesneau et al., 1994; Mansour et al., 1995; Merg et al., 2006; Morgan et al., 2017), with a concomitant shift from negative to positive emotional affect (Valentino and Volkow, 2018). *OPRK1* agonism is well known to elicit aversive, anhedonic, and depressive-like states in both humans and preclinical rodent models; conversely, antagonism evokes anti-depressive states (Peciña et al., 2019). *OPRD1* agonists, on the other hand, have anxiolytic and antidepressant activity (Valentino and Volkow, 2018). In order to develop a therapeutic to leverage our insights from the human genetic association, ideally one would develop an activator of *NRDC* enzymatic activity, but it is generally infeasible to activate enzymes pharmacologically. Instead, we could theoretically mimic the effect of *NRDC* activation with a dual *OPRK1* antagonist/*OPRD1* agonist combination therapy. In fact, similar strategies are already being pursued in the clinic for depression, including a selective *OPRK1* antagonist and buprenorphine, a *OPRK1* antagonist with partial agonist activity at the *μ*-opioid receptor, *OPRM1* (Jacobson et al., 2020). The early results from these clinical trials are encouraging. Selective *OPRK1* antagonism is not addictive, and buprenorphine in particular has demonstrated immediate efficacy, with no short-term increase in suicide risk (unlike other depression drugs), that is observed even in patients with treatment-resistant depression (Peciña et al., 2019; Jacobson et al., 2020). While these promising therapies were presumably identified on the basis of the preclinical data, we independently identified a similar therapeutic strategy directly from the human genetics of depression.

## DISCUSSION

Previous GWAS methods have explored different solutions for finding trait-associated genes, by cleverly circumventing the major challenges described at the outset. One class of methods proposes causal genes on the basis of prior knowledge (e.g., text mining, gene co-expression, or biological networks; Raychaudhuri et al., 2009; Rossin et al., 2011; Pers et al., 2015; Taşan et al., 2015; Greene et al., 2015). These algorithms implicate genes in trait-associated loci that may be functionally related to genes at other, independently associated loci. This approach generates hypotheses without requiring direct experimental evidence that the implicated genes are targeted by causal variants or dysregulated in disease. Other methods generate hypotheses of trait association by computing rigorous gene-level p-values. One such approach identifies genes that overlap a statistically significant association signal (de Leeuw et al., 2015). Another approach, TWAS, integrates GWAS data with eQTL (gene expression) studies, to identify genes whose expression is mediated by variants in GWAS loci (Gamazon et al., 2015; Gusev et al., 2016; Barbeira et al., 2018). A sophisticated extension to TWAS can compute probabilities of disease association for genes (Mancuso et al., 2019), when the disease is mediated by a single cell type with well-powered eQTL data. Related methods leverage GWAS loci where the association signal has the same statistical structure in an eQTL study (Giambartolomei et al., 2014; Zhu et al., 2016; Hormozdiari et al., 2016). Matching association signals in eQTLs and GWAS loci indicate that the same causal variant(s) mediate both disease risk and the expression of particular genes. Whenever such loci can be found, these methods identify the genes targeted by causal variants (Giambartolomei et al., 2014; Zhu et al., 2016; Hormozdiari et al., 2016).

Here, we took advantage of theoretical and experimental advances that were not available during the development of these previous methods. These advances enable us to compute the probability that each gene is targeted by a causal variant, without the requirement for eQTL data or assumptions about the disease tissue(s). We merely assume that the variant-to-gene map is correct. (Relaxing that assumption simply qualifies the interpretation of the gene PPAs, whose meaning becomes: the probability that a gene is targeted by a causal variant, *according to the variant-to-gene map*.) We demonstrate the sensitivity of the platform, which often finds more trait-associated gene clusters than there are GWS causal variants. That sensitivity translates into a remarkable recall of known disease genes, which may be ~80% of the genes implicated by GWAS signals, and is substantially greater than the current state of the art. We compare this paradigm to a recent method (Nasser et al., 2021) that links fine-mapped variants to target genes (Note S2). The new framework identifies many more disease genes than are currently well-known, enabling manifold opportunities for novel drug development and biological discovery.

We expect the power of this new paradigm, and its implications for biological discovery, to scale with the available GWAS data. New studies of large disease cohorts, with associated longitudinal clinical data, will expand the spectrum of diseases amenable to genetic analysis. Furthermore, as GWAS sample sizes increase, the uncertainty in the statistical calculations will diminish. Not only will the *number* of detectable association signals improve, but the *proportion* of these that can be fine-mapped to causal variants and trait-associated genes will increase. In addition, as the cost of sequencing continues to fall, large whole genome sequencing (WGS) studies are becoming feasible. WGS will uncover novel and ultra-rare variants, but it will also improve the genotyping of relatively more common variants (minor allele frequency, MAF ≥ ~0.1%) that are not well-imputed with existing reference panels. Combinations of complementary analysis methods may be required to fully leverage WGS data. The statistical framework we introduce is well-powered to detect associations induced by rare variants (MAF ≈ 0.1%). On the other hand, family-based studies (Ronemus et al., 2014) and statistical methods designed specifically for ultra-rare variants (Lee et al., 2014) will be necessary to uncover disease associations from the rarest variants. It may be fruitful to combine the methods specialized for these variants with the variant-to-gene map. An outstanding question is whether there are ultra-rare variants in enhancers, uncovered from WGS, that mediate disease risk. Integrating the methods for ultra-rare variants with the variant-to-gene map may address this question, and potentially discover novel disease genes.

## Acknowledgments

We thank Dr. Tom Maniatis, Dr. Simon Tavaré, and Dr. Charles Zuker for critical reading of the manuscript and insightful comments. This research has been conducted using the UK Biobank Resource under Application Number 41018.

## Author contributions

J.N.J.M. conceived the overall framework and its individual components, and supervised the work. J.N.J.M. and S.C. developed the theory for partitioning the genome into LD blocks; S.C. designed, implemented, and proved the correctness of the algorithms to find the optimal population-specific and universal LD partitions. J.N.J.M. and R.J.L. developed the theory and implementation of the variant-to-gene map, including the machine learning framework to predict enhancer-promoter interactions. R.J.L. validated the machine learning models. J.N.J.M. extended the statistical fine-mapping equations into the domain of genes and implemented the fine-mapping algorithm. J.N.J.M. and D.L. developed the theory for learning parameters of the statistical model with expectation maximization (EM). D.L. implemented the EM algorithm and a distributed computational architecture to run the framework. J.N.J.M. collected the GWA studies and analyzed them with the platform. J.N.J.M. and R.J.L. compared the performance of the framework with the state of the art. J.N.J.M wrote the manuscript, with contributions from R.J.L., D.L., and S.C. All authors reviewed and approved the final manuscript.

## METHODS

### PARTITIONING THE GENOME INTO LD BLOCKS

#### Filtering of 1KGP Variants

For each 1KGP super population, we filtered the variants included in the sample correlation matrices according to quality control criteria. We excluded multiallelic polymorphisms (which account for a small fraction of variants), as well as biallelic variants with a minor allele frequency (MAF) less than 0.05. Variants with a low MAF often induce spurious long-range correlations in small sample sizes. Excluding these uncommon variants precludes the need for shrinkage estimation of the correlation matrix, while preserving enough genetic variation to obtain a high-resolution LD map from the raw data. Otherwise, shrinkage estimation to correct sampling errors will change the statistical structure of the variant correlation matrix and may introduce bias, for instance by relying on recombination rates averaged over multiple populations (Berisa and Pickrell, 2016).

Due to the filtering criteria, some variants are excluded from the LD block analysis. Since the block boundaries are defined at the locations of variants included in the analysis, this means that some of the excluded variants may fall between blocks. To assess the coverage of our LD blocks, we computed the percentage of all 1KGP variants (including multiallelic and uncommon variants) that fall within an LD block. More than 99.5% of 1KGP variants are encompassed within the LD blocks defined for any super population, and the consensus blocks cover more than 99.9% of all such variants.

#### Recombination Analyses

We calculated the recombination rates for each super population by taking a weighted average of the rates reported for individual subpopulations (Auton, 2013), according to the proportion of each subpopulation in the super population. In order to compute the frequency of the consensus recombination hotspot motif at the borders between LD blocks, we used the algorithm described by Mackiewicz et al. (2013).

#### LD Block Discovery as a Graph Clustering Problem

We show that partitioning the genome into LD blocks is equivalent to clustering a weighted graph embedded on a line, and we provide the optimal solution to this clustering problem. Consider a graph whose vertices are the genetic variants on a chromosome. Further, let the edge weight between any two vertices be the squared sample correlation, *r*^2^, between the corresponding variants. Then the desired partition of the genome into LD blocks is nothing more than a clustering of this graph, subject to the geometric constraint that clusters must be composed of physically contiguous variants on the chromosome. That is, we seek a clustering which groups strongly connected vertices into the same cluster, while dividing weakly connected vertices into different clusters. If we require that the optimal clusters map to non-overlapping, abutting intervals along a line (the chromosome), then this is equivalent to finding LD blocks in which correlated variants are grouped together but uncorrelated variants are separated. This clustering is depicted schematically in Figure S1. See Berisa and Pickrell (2016) for a justification of *r*^2^ as the LD metric.

#### Modularity Clustering

The clustering of weighted graphs is a well-studied problem in the computer science literature (Fortunato, 2010; Fortunato and Hric, 2016; Buluç et al., 2016), from which modularity clustering, and its derivatives, have emerged as leading solutions (Blondel et al., 2008; Traag et al., 2019). Modularity clustering aims to find a partition that maximizes the modularity function (Blondel et al., 2008):

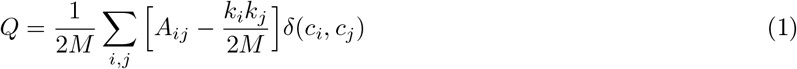

where *M* is the total edge weight in the graph; *A_ij_* is the edge weight between vertices *i* and *j*; *k_i_* ∑_*j*_ *A_ij_* is the total edge weight incident on the *i*-th vertex; *c_i_* is the cluster to which vertex *i* is assigned; and *δ*(*c_i_*,*c_j_*) is the indicator function whose value is 1 if vertices *i* and *j* fall into the same cluster and 0 otherwise. The term *k_i_k_j_*/2M in equation (1) is the expected edge weight between vertices *i* and *j* in a randomized network, in which the interaction partners of each vertex are randomly reassigned while preserving the degree *k_i_* of each vertex (Newman and Girvan, 2004; Newman, 2006). Therefore, the modularity function *Q* measures the total edge weight between all co-clustered vertices, *relative to a random network* and normalized to take values in the range [–1,1].

Despite its ubiquity in clustering applications, there are two major limitations of modularity clustering, which spur continued research in this domain. The first is the well-known ‘resolution limit’ of the modularity function (Fortunato and Barthelemy, 2007; Fortunato, 2010). The null model (i.e., randomized network) in the modularity function assumes that any vertex is equally likely to connect with any other vertex in the graph, even though real networks have local structure that circumscribes sets of plausibly connected vertices. Consequently, as the size of the input graph grows, the expected edge weight between any two vertices in the null model becomes progressively smaller, leading to a weaker penalty for co-clustering poorly connected vertices and a lower clustering resolution. In our context, this means that the clustering resolution (and therefore LD block size) would be artificially skewed by the density of genetic variants and the length of chromosome segments under consideration. The second challenge is that, for arbitrary graphs, finding a clustering that optimizes the modularity function is an NP-hard problem (Fortunato and Barthelemy, 2007; Fortunato, 2010). Exact solutions to find the global optimum are not available; instead, current research focuses on the development of fast algorithms to find progressively better local optima (Blondel et al., 2008; Traag et al., 2019).

#### LD Modularity

We develop a definitive algorithm for clustering LD blocks that overcomes the limitations of modularity clustering, by leveraging prior knowledge of the graph induced by the variant correlation matrix. Consider, first, the resolution limit. In general, the appropriate null model for any clustering problem is unknown, because it depends on the local statistical structure of the graph. In our setting, however, the null model is simply the expected sample correlation between two uncorrelated variants. Because the sample correlation is a biased estimator of the true population correlation, this null model is not zero. If we restrict ourselves to biallelic variants whose genotypes are encoded as 0 (indicating the major allele) and 1 (indicating the minor allele), then the sample Pearson correlation *r* between any two variants is mathematically equivalent to the sample Phi correlation *ϕ*. And from Tschuprow (1939), we know that the expected sample Phi correlation between uncorrelated variables is given by

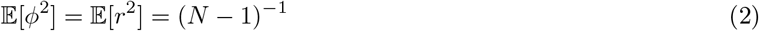

where *N* is the sample size. Given this statistically rigorous null model, we can define the desired resolution of the LD clustering explicitly. Let 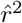 denote the minimum squared Pearson (Phi) correlation between two variants that are said to be in LD. Then we replace the generic null model from the modularity score with a resolution-controlled alternative, to obtain our (unnormalized) ‘LD modularity’ objective:

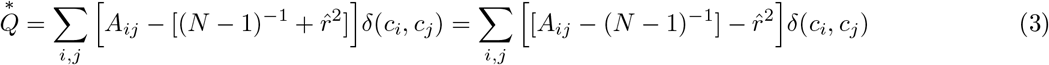

This objective corrects each sample correlation by the expectation under the null (*A_ij_* – [*N* – 1]^−1^) and compares the corrected correlation to the threshold 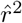. The corrected correlations satisfy

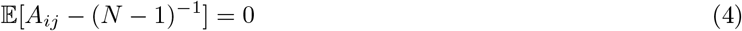

for uncorrelated variants *i* and *j*. Therefore, the LD modularity has the desired property: Bias-corrected correlations (*A_ij_* – [*N* – 1]^−1^) which exceed the threshold 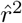 will incentivize grouping the corresponding loci into the same LD block (by increasing 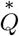). Conversely, corrected correlations below this threshold will promote the separation of loci into different LD blocks. In practice, we set 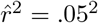. Below, we show that the penalty term 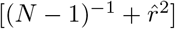 can also be interpreted as the critical value of the *χ*^2^ statistical test, whose significance level *α_N_* is a function of the sample size *N*. That is, the LD modularity tends to group locations into the same LD block only if their correlation is statistically significant, at a confidence level that becomes stricter with larger sample sizes. Smaller populations enjoy a relaxed statistical threshold, ensuring that real but marginally significant correlations are not ignored. The key terms of the LD modularity objective are evaluated for each 1KGP super population in Table S3.

#### The Clustering Algorithm Groups Loci with Significant Correlations

Beyond our derivation of the null model in the LD modularity, from Tschuprow’s equation, there exists an even deeper statistical justification for our optimization criterion. Namely, our model satisfies the property that suprathreshold edge weights, which drive the agglomeration of sites into LD blocks, occur only between loci with a statistically significant correlation. In fact, we identify our penalty term as the critical value of the *χ*^2^ statistical test, whose significance level is a function of the sample size *N*. (The *χ*^2^ test is the asymptotically valid procedure for testing the significance of variant correlations.) The connection between our model and the *χ*^2^ test follows from the fact that the *r*^2^ correlation (for biallelic genotypes) is nothing more than the *χ*^2^ test statistic divided by the sample size:

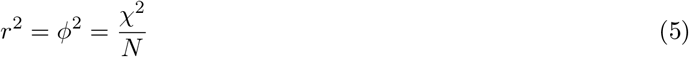

In any empirical statistical test, the critical value of the test statistic is the threshold beyond which the test statistic is considered significant. In the present case, that corresponds to the smallest correlation, 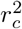, that yields a suprathreshold edge weight. Since 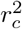 is merely the penalty term

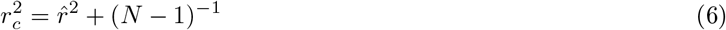

then the critical value of the test statistic is:

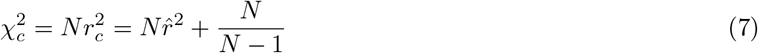

As with any one-sided statistical test, the confidence level *α* at which we reject the null hypothesis is the area under the cumulative distribution function (CDF) to the right (or left) of the critical value. In our case, the significance level *α_N_* at which we deem a correlation to be significant is therefore

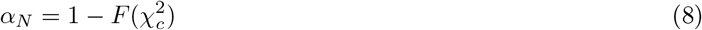

where 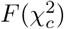 denotes the CDF of the *χ*^2^ distribution evaluated at the critical value. The null hypothesis of independence will be rejected at significance level *α_N_* for any pair of genomic sites with an observed correlation exceeding 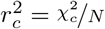. Therefore, two sites will have a suprathreshold edge weight iff their sample correlation is statistically significant at level *α_N_*. Our notation signifies that the confidence level depends on the sample size *N*. Smaller sample sizes yield smaller critical values, and therefore less stringent significance levels (Table S3).

#### Optimizing the LD Modularity

Now consider the optimization of the LD modularity, whereby we maximize suprathreshold correlations within LD blocks while minimizing correlations between them. We leverage the constraint that clusters must map to contiguous segments on the chromosome, to reduce the feasible solution space of the problem and to develop an algorithm that is guaranteed to find the globally optimum clustering of each chromosome.

The optimization algorithm comprises two dynamic programming (DP) stages. First, we score all possible valid clusters of the graph using DP. Let *n* denote the number of genetic variants on a chromosome and *l* the maximum number of variants in an LD block, where *l* ≤ *n*. Then there are at most *nl* possible clusters, since there are *n* possible starting coordinates and *l* available ending positions, given a choice of start position. This number is dramatically fewer than the number of possible clusters in a graph without restrictions. In the second step, we again use DP to choose a subset of these clusters that form a valid partition of the graph and that maximize the optimization criterion. Overall, the algorithm has a computational complexity of order *O*(*nl*). In practice, we choose *l* < *n* as a computational convenience (because LD blocks spanning an entire autosome are implausible), while ensuring that *l* is sufficiently large to allow for arbitrarily wide LD blocks. A more detailed discussion of the algorithm, the running time analysis, proofs of correctness, and discussion of the parallel implementation can be found in the next section.

#### Clustering Algorithm

##### Input

We are given as input a weighted graph *G* = (*V*, *E*) where |*V*| = *n* and |*E*| = *m*, along with some weight function 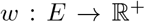. We are also given a linear ordering of the vertices via an injective coordinate function 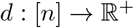, where *d*(*i*) denotes the coordinate of vertex *v_i_* and no two vertices have the same coordinate. WLOG, we assume that the indexing of the vertices, *v*_1_, *v*_2_,…, *v_n_*, respects the ordering imposed by these coordinates.

Finally, we are given a maximum distance *D* beyond which two vertices cannot be assigned to the same cluster (i.e., |*d*(*i*) – *d*(*j*)| ≤ *D* if *v_i_* and *v_j_* are in the same cluster). In practice, if *D* is set to a value greater than *v_n_* – *v*_1_ then this will not be a tight constraint. However, there are certain contexts in which prior knowledge may inform the selection of a maximum distance, and setting this parameter can reduce the running time. We use *l* to denote the maximum *number* of vertices within a distance of D.

##### Partitions

A *partition P* of the graph *G* is an exact cover of *V*. In other words, a partition of *G* is an assignment of its vertices to a set of clusters *P* = {*C*_1_, *C*_2_,…, *C_k_*} (for some 1 ≤ *k* ≤ *n*) such that:

- For all *v* ∈ *V*, there exists some cluster *C_i_* such that *v* ∈ *C_i_*.
- For all *i*,*j* ∈ [1, *k*] such that *i* ≠ *j*, we have 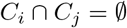.

A *linear partition* of the graph *G* with respect to coordinate function *d* is a partition with the additional constraint that:

- For all *i* ∈ [1, *k*], if *v_a_*, *v_c_* ∈ *C_i_*, then for any *v_b_* such that *d*(*a*) < *d*(*b*) < *d*(*c*), *v_b_* ∈ *C_i_*.

That is, if we ordered the vertices according to their coordinates along a line, a linear partition can be achieved by making cuts along this line.

##### Problem Statement

We propose an algorithm to find the clustering of a graph that achieves the global optimum of a modularity objective, subject to the following constraints:

1. We require that the optimal clustering be a linear partition.
2. The distance between coordinates of vertices assigned to the same cluster may not exceed the maximum distance *D*.

Our optimization algorithm finds the global maximum of any modularity objective that can be written as a sum of independent pairwise interactions between vertices. To demonstrate the generality of our method, we employ it to optimize the standard modularity function *Q*, in addition to the new LD modularity objective 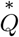 we introduce in this work. We show that *Q* and 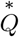 can be optimized with the same algorithm in Section *Preprocessing*. We then demonstrate how the clustering score for all possible valid components of the graph can be efficiently computed with DP in Section *DP to Calculate Cluster Scores*. Finally, we show how DP can again be used to find a valid partition of the graph that maximizes the desired clustering score, in Section *DP to Find Optimal Linear Partition*.

##### Preprocessing

For both *Q* and 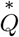, we need only consider the sum over *unique* (ordered) pairs of vertices. The contributions from self-loops (*A_ii_* ∀*i*) can be ignored, because they are constant (i.e., independent of the partition *P*). Then, since *A_ij_* = *A_ji_*, restricting the sum to ordered pairs is equivalent to multiplying the objective by a constant factor of ½. Likewise, when maximizing the modularity *Q*, we can ignore the normalization term (2*M*)^−1^ because it, too, is constant. Then 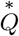 differs from *Q* only in its penalty term (i.e., null model), which is simpler. After some preprocessing, we show that these two problems can be solved with the same algorithm. Note that in both cases, any *A_ij_* that is not explicitly given as an input edge weight is considered to be 0.

##### Standard modularity

First, we calculate *M* directly by summing all the weights in the input. Next, we initialize a 1 × *n* vector *K* to store the value of *k_i_* for each vertex *i*. We iterate over the input weights and store the sums. We then compute the *normalized weight* of an edge for all pairs of vertices *v_i_* and *v_j_* such that *d*(*j*) – *d*(*i*) ≤ *D*:

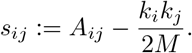

##### LD modularity

The preprocessing for LD modularity is even simpler. Here we merely subtract a constant to compute the normalized weight of an edge, again for all pairs of vertices *v_i_* and *v_j_* such that *d*(*j*) – *d*(*i*) ≤ *D*:

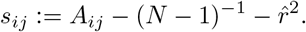

Now, the problem of optimizing either objective reduces to maximizing the sum of the normalized weights *s_ij_* within partition components (i.e., clusters).

##### Step 1 DP to Calculate Cluster Scores

Inspecting the structure of equations (1) and (3), we observe that each cluster contributes independently to the objective function. We therefore separately compute the score of every possible cluster. We rearrange equation (1) to emphasize this point:

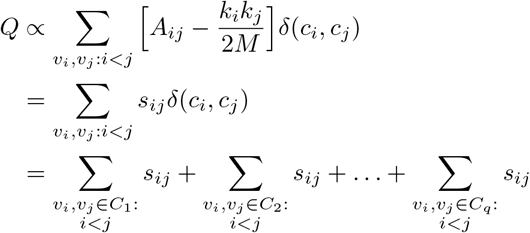

where *q* is the number of clusters in the partition *P*. We define the cluster score, for any cluster *C*, as the sum over distinct pairs of vertices in *C*, 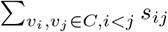. Note that there are at most *O*(*nl*) possible clusters given the constraints on our optimization problem (# of starting points × # of ending points). We now define the following function:

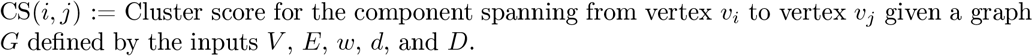

For a given cluster *C* starting at vertex *v_i_* and ending at vertex *v_j_* (w.r.t the ordering *d*), we can calculate CS(*i*, *j*) directly from the definition:

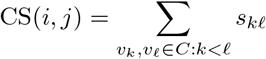

Since evaluating this equation for one component requires *O*(|*C*|^2^) = *O*(*l*^2^) time, computing this quantity directly for each of *O*(*nl*) possible components would take *O*(*nl*^3^) time.

We can speed this up by using dynamic programming. Consider the following recursion:

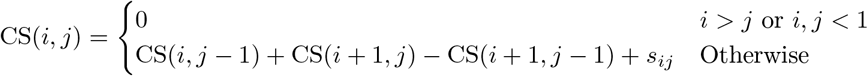

We show that this recursive equation holds with a short proof by induction.

*Proof*. We perform induction on the size of the component (i.e., |*j* – *i*|).

###### Base Case

First, we consider the well-defined case where the length of the component is 0 (i.e. *i* = *j*). We get the following as desired:

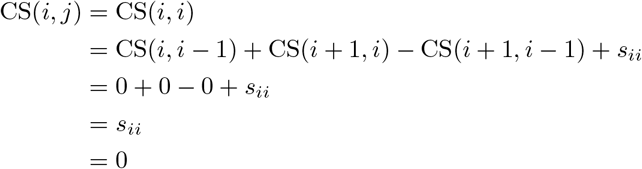

###### Inductive Step

Now assume that the recursion holds for components of size 0 to *n*. Consider a component ranging from *v_i_* to *v_j_* where *j* – *i* = *n* + 1. We have that:

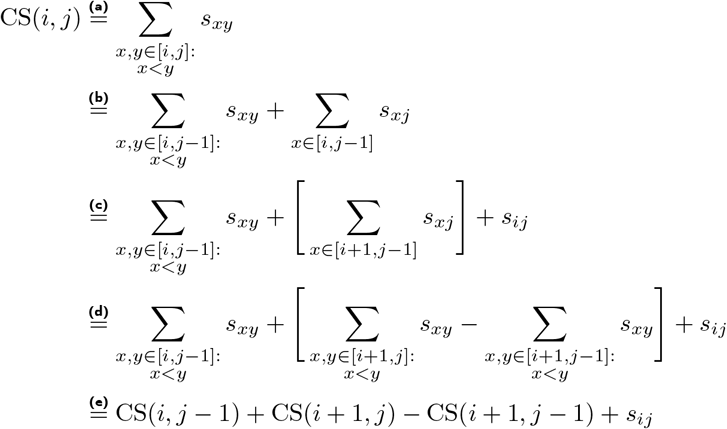

where (a) is by definition; (b) pulls out terms involving *v_j_*; (c) pulls out *s_ij_* from the second term; (d) recognizes the sum over pairs with *v_j_* as the sum over *all* pairs, minus the pairs that do not include *v_j_*; and finally (e) is by definition. Thus, by the inductive hypothesis, our recursion holds.

##### Step 2 DP to Find Optimal Linear Partition

After computing each cluster’s contribution to the modularity, we need to select a subset of these clusters that form a valid linear partition and that maximize the clustering score. We achieve this efficiently by using dynamic programming once again. We define another recursive function:

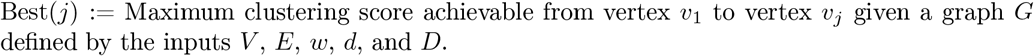

Note that this equation differs from CS(1, *j*) in that Best(*j*) is made up of (potentially) many different, non-overlapping components.

We calculate Best(*j*), defined for *j* ≥ 0, using the following recursion:

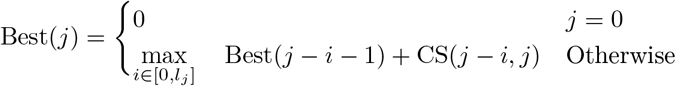

The variable *l_j_* ≤ *l* denotes the number of variants *v_i_* such that |*d*(*i*) – *d*(*j*)| ≤ *D* and *i* < *j*. Intuitively, we consider all possible clusters to which the final variant *j* may be assigned, and then recurse on the possibilities for the remainder of the chromosome.

*Proof*. We now give another short proof by induction on the vertex coordinate j, signifying the end of the window [1, *j*].

###### Base Case

If *j* = 1, then *l_j_* = 0 since there are no variants with an index less than 1. We then get the correct partition corresponding to the score of just the single vertex:

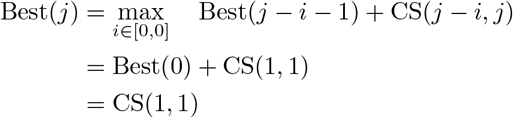

Note that Best(0) isn’t meaningful, but has been defined out of convenience.

###### Inductive Step

Now assume that the recursion holds for all endpoints 1 up to some index *n* – 1. Consider index *n*. Let *Opt* be the score of some optimal partition *P_opt_* that clusters all variants from 1 to *n*.

By the definition of a partition, the rightmost vertex *v_n_* must be in exactly one cluster of *P_opt_*. Any valid cluster containing *v_n_* as its final variant must be contiguous and may hold no more than *l_n_* additional variants (by the distance constraint). Therefore, there are exactly *l_n_* + 1 possible clusters to which *v_n_* can belong, corresponding to *l_n_* + 1 possible positions at which the left boundary of the cluster may be placed. (This includes *l_n_* possibilities in which the leftmost variant is strictly less than *n*, plus the single possibility whereby the leftmost and rightmost variant are the same—i.e, where *v_n_* is the only vertex in its cluster). Therefore, CS(*n* – *i*, *n*) ∀*i* ∈ [0, *l_n_*] are the score contributions from all valid clusters in which vn can be included.

Taking the maximum over the scores achieved from all possible clusters containing *n*, plus the optimal score contributed by all other clusters, exhaustively covers all possibilities:

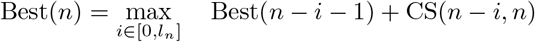

Best(*n* – *i* – 1) is well-defined since *l_n_* < *n*. By the inductive hypothesis, Best(*n* – *i* – 1) contains the best score achievable on the remaining portion of the chromosome, given a choice for the final cluster encompassing *v_n_*. Since we examine all possible final clusters, and choose the option which maximizes the aggregate score of *all* clusters, Best(*n*) = *O_pt_*.

To find the optimal clustering score over the full graph, we will simply evaluate Best(*n*), where *n* is the index of the last variant on the chromosome. We then perform standard backtracking to recover a partition that achieved this optimal score.

#### Running Time Analysis

Recall that |*V*| = *n*, |*E*| = *m*, and *l* is the maximum number of vertices within a distance of *D*. Since we disregard any edge whereby the distance between the two endpoints is greater than *D*, this implies that *m* ≤ *n* · *l* ≤ *n*^2^.

##### Preprocessing

For the modularity score, it is straightforward to see that *M* can be calculated directly in *O*(*m*) time. Similarly, we can compute *k_i_* for all *i* by simply taking one pass over the edge weights in *O*(*m*) time. We can then directly compute the normalized weight of an edge for all pairs of vertices *i* and *j* such that *d*(*j*) – *d*(*i*) ≤ *D* in *O*(*nl*) time. It is easy to see that the preprocessing for the LD modularity has the same time complexity (i.e., it is linear in the number of edges). Thus, all the preprocessing for either score can be done in *O*(*nl*) time and *O*(*nl*) space.

##### Step 1 Calculating all cluster scores

In practice, we set the order of enumeration for this calculation based on the cluster size, starting with clusters of size 1 and ending with clusters of size *l* + 1. Visually, this looks like traversing diagonals on a matrix from the upper left to the bottom right. There are *O*(*nl*) subproblems, each of which takes constant time to compute (performing 4 lookups and then adding these 4 numbers). Thus, the overall running time for this step is *O*(*nl*) and will likewise require *O*(*nl*) space to store all of the cluster scores.

##### Step 2 Finding optimal linear partition

Note that in this step there are *n* subproblems, each of which take *O*(*l*) time to evaluate. This gives an overall running time of *O*(*nl*) and requires *O*(*nl*) space. In addition to storing the maximum score for each subproblem, it is also necessary to store the value *i* ∈ [0, *l*] which achieved this maximum score. These *i* values correspond to the cuts on the chromosome that give us an optimal partition. Backtracking can likewise be done in *O*(*nl*) time.

Finally, across all steps this algorithm takes *O*(*nl*) time and *O*(*nl*) space.

#### Implementation Details

The algorithm has been implemented in C++. We use OpenMP to parallelize the inner loops of both steps 1 and 2. For larger chromosomes, *O*(*nl*) space may be too large to fit in working memory. Our implementation allows the user to specify an additional parameter that chunks the chromosome into smaller pieces, solves these pieces with memory proportionate to the chunk size, and stitches the results back together. This feature does not impact the accuracy of the calculation, but increases the running time when reading and writing to disk, in exchange for less memory usage.

#### Consensus-finding algorithm

Given sets of LD blocks from different populations, the consensus algorithm seeks to find common endpoints shared by every population. In general, however, the boundaries of population-specific LD blocks will not align on precisely the same variant. Different populations are partitioned on slightly different subsets of variants (due to the MAF filter); recombination hotspots are not individual nucleotides but rather 1–2 kb genomic ranges (Wall and Pritchard, 2003; Paigen and Petkov, 2010); and sampling noise will introduce variability in the precise breakpoints between LD blocks in each population. Therefore, rather than identify precise nucleotides at universal LD borders, we instead search for narrow *regions* along the chromosome which contain an LD boundary from each population. We refer to these regions as cuts, which partition the genome into universal LD blocks without bisecting any of the input blocks. By requiring that the cuts be narrow (with a maximum width of *β* = 2 kb), we ensure that the boundaries between our universal LD blocks are as highly resolved as recombination hotspots themselves.

##### Consensus cut problem

We are given as input a set of blocks *B*, where each block *b* ∈ *B* is defined by a start point *s_b_* and an endpoint *e_b_*. Let *s_min_* be the smallest coordinate in *B* and *e_max_* be the largest coordinate in *B*. Likewise, a cut *c* is a closed interval defined by a start and endpoint: [*s_c_, e_c_*]. A cut is *eligible* if the following holds:

1. *s_c_* and *e_c_* are endpoints in *B* (not necessarily from the same population).
2. There does not exist an input block *b* such that *s_b_* < *s_c_* and *e_b_* > *e_c_*.
3. *e_c_* – *s_c_* ≤ *β*.

Furthermore, we say that two cuts are disjoint if the genomic intervals which define them do not touch; the width of any cut is the width of its interval. The *consensus cut problem* seeks to find the largest set of disjoint eligible cuts (i.e., the most highly resolved universal LD blocks). If more than one maximal set exists, we choose the one with the smallest total cut width (i.e., the best-resolved LD boundaries).

##### Consensus cut algorithm

To solve the consensus problem, we first identify a set of narrow eligible cuts. We then perform dynamic programming over this set of eligible cuts to find a maximal, disjoint subset.

To identify eligible cuts, we iterate over all endpoints in *B*. For each endpoint *p*, we identify the set *C_p_* of blocks in *B* with *s_b_* < *p* ≤ *e_b_* (i.e., blocks spanning *p*). Let *e_c_* be the largest endpoint in *C_p_*. If *e_c_* – *p* ≤ *β*, then we add [*p*, *e_c_*] to our list of eligible cuts. Given our definition of an eligible cut, [*p*, *e_c_*] is the most narrow eligible cut containing the coordinate *p*. All eligible cuts are required to start at endpoints in B, and this algorithm is guaranteed to return the narrowest eligible cut for each endpoint (if one exists). Given our optimization criteria, any cut which is wider than necessary cannot be optimal. Therefore, when applied to all endpoints *p* ∈ *B*, this procedure generates the complete set of eligible cuts that may comprise an optimal solution.

Next, we use dynamic programming to find a maximal, disjoint subset of these eligible cuts. We sort the eligible cuts in ascending order of their start point (i.e., [*s_i_*, *e_i_*] is the cut with the *i*-th smallest start index). Assume there are *k* eligible cuts in total. And let *MC*(*j*) denote the maximal number of disjoint cuts contained within the range *s_j_* to *e_max_*. We can calculate *MC*(*j*) recursively:

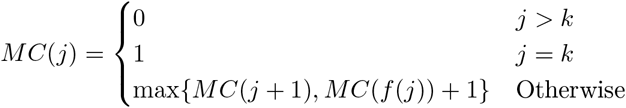

where *f*(*j*) maps index *j* to the next largest index *i* such that *e_j_* < *s_i_* (if none exists, *j* maps to *k* +1). If both terms in the argument to max{} are equal, we store the one with the smallest total cut width. For each value of *j*, the recursion chooses between the best of two options: ignoring the *j*-th cut, or taking the *j*-th cut. If we don’t take cut *j*, then we move to the index of the very next (possibly overlapping) cut; otherwise, we take the cut *j*, move to the nearest *disjoint* cut, and increment our count of disjoint cuts. We call *MC*(1) to get the maximal number of disjoint cuts over the entire range of input blocks, store solutions to all subproblems, and perform backtracking to return an optimal solution.

##### Obtaining consensus blocks from cuts

The definition of universal LD blocks follows immediately from the enumeration of the optimal cuts. The universal LD blocks are the genomic regions intervening the optimal cuts, starting and ending with the positions nearest to each cut.

### MAPPING ENHANCERS AND VARIANTS TO GENES

#### Data Acquisition

The enhancer-to-gene modeling requires: (1) a set of candidate regulatory elements; (2) assays of chromatin accessibility at these elements across the gamut of available cell types and tissues; (3) assays of transcriptional activity, to validate the identification of regulatory elements; and (4) assays of chromatin contact frequency, to measure physical interactions between putative enhancer-promoter pairs. We obtained the data for each of these modalities from publicly available sources.

##### Atlas of open chromatin elements

*cis*-Regulatory elements reside within open-chromatin (i.e., DNase-I hypersensitive sites, DHSs), allowing for access by transcription factors or other transcriptional machinery (Andersson et al., 2014). These DHSs can be identified experimentally through chromatin accessibility assays, including DNase-seq and ATAC-seq, and are often highly cell-type specific (Elkon and Agami, 2017). To construct our enhancer-to-gene map, we require a cell-type agnostic atlas of open chromatin; that is, the union of DHSs across many human cell types and tissues. We therefore downloaded the set of DHSs, 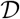, identified by ENCODE (Meuleman et al., 2020), composed of ~3.5 million sites which we consider candidate regulatory elements (promoters or enhancers). This atlas of DHSs has a total coverage of 664.05 Mbp, or approximately 21% of the human genome.

##### Open chromatin sequencing

We treat chromatin accessibility as a continuous variable, by computing normalized read counts from sequencing assays that target open chromatin. Using the ENCODE portal (ENCODEproject.org; Davis et al., 2018), we downloaded aligned read (.bam) files from chromatin accessibility assays (DNase-seq and ATAC-seq) contributed by the ENCODE, Roadmap Epigenomics, and Genomics of Gene Regulation consortia. These data contain 2005 samples (set 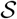) and 493 distinct pairs of biosample types and donor life stages as annotated by ENCODE (set 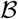). The samples include human bulk tissue, primary cells, *in vitro* differentiated cells, and cell lines, comprising all chromatin accessibility data in *H. sapiens* available from ENCODE as of March 2021 (see File S3 for a list of ENCODE accession numbers and associated metadata for each sample used in this work). We downloaded all aligned reads in GRCh38 coordinates; converted the reads to BED files with the `bamtobed` function in `BEDtools`; and finally mapped the reads to the DHS atlas in the same coordinates. For subsequent analyses, because many GWAS are available only in GRCh37 coordinates, we used liftOver to map the DHS atlas to GRCh37 coordinates, retaining only those DHSs that map one-to-one between genome builds. Lastly, we normalized the read counts in each DHS to RPKM (Reads Per Kilobase of transcript, per Million mapped reads) using `edgeR` (McCarthy et al., 2012) and treating the DHS width as analogous to transcript length. The result is a normalized measure of how many DNase- or ATAC-seq reads overlap each DHS, in each sample. Hereafter, DHS ‘activity’ refers to these normalized read counts in RPKM.

ATAC-seq and DNase-seq both measure chromatin accessibility, but have assay-specific effects that may confound an analysis that includes data from both technologies (Buenrostro et al., 2015). Therefore, we developed a simple method for assimilating ATAC-seq and DNase-seq data into the same analysis, under the assumption that each assay measures the same biological phenomenon but with a different, potentially nonlinear measurement function. Under this assumption, sequence biases of the enzymes used by the assays (a modified TN5 transposase and DNase-I, respectively) are considered negligible. This assumption is plausible for DHS-scale measurements, where strong correlations are seen between DNase-I seq and ATAC-seq data (Buenrostro et al., 2015; Martins et al., 2018). Specifically, we assumed that there exists a monotonic function from the expected DHS activity measured by ATAC-seq to the expected DHS activity measured by DNase-seq. To learn this function from data, we identified all the ATAC-seq samples available from ENCODE that were also assayed using DNase-seq (where a ‘sample’ must be from the same tissue/cell type and originate from the same donor). Using the `scam` R package (Pya, 2019), we fit monotonic smoothing splines to relate the log-transformed ATAC-seq activity to the log-transformed DNase-seq activity for each of these pairs. (To compute log-transformed RPKM, we added a pseudo-count to all measured values, equal to one half the minimum observed nonzero activity for each sample). We observed a common signature across nearly all samples, such that the ATAC-seq activity was consistently lower than DNase-seq at low values, and higher at high values. (The ATAC-seq signal has a larger dynamic range.) Ultimately, we chose to model the transformation from ATAC-seq to DNase-seq with the spline fit to data from adrenal gland tissue, since this sample had the highest geometric mean sequencing depth between the two assays. We then used the same model to transform normalized ATAC-seq read counts to DNase-seq data for all of the ATAC-seq samples (Figure S11).

Due to considerations of computational complexity in our regression models (see below), we merged data from multiple assays of the same or similar cell types. We manually curated the ENCODE biosample types and donor life-stage information to generate a surjective map from ENCODE metadata to a set of qualitatively defined ‘cell types’ 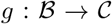, which is available in File S4. Using our qualitative cell type definitions, there are 201 unique cell types, the majority of which (165) have been assayed multiple times; 86 were assayed greater than three times. For samples assayed more than three times, we computed the arithmetic mean of the activity at each DHS, to merge samples together such that no cell type is included more than three times as an observation in the regression models. (We allow up to three observations of the same cell type, so that when we perform three-fold cross-validation, we can balance replicates of the same cell type between folds to the greatest extent possible.) This procedure resulted in 464 observations in the regression models, denoted by the set 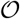. Using these observations, we define the DHS matrix 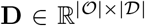, which includes the DHS activity in all observations at all DHSs, with each column mean-centered and standardized.

##### Hi-C data

In order to measure contact frequencies between genomic loci in three-dimensional space, we downloaded processed Hi-C data files (.hic files) from the Hi-C data archive at http://www.aidenlab.org/data.html for GM12878, HeLa, HMEC, HUVEC, K562, and NHEK cells (Rao et al., 2014). We computed normalized Hi-C contact matrices at 5 kbp resolution using the Knight-Ruiz matrix balancing algorithm (Knight and Ruiz, 2012) as implemented in the `Juicer` Hi-C analysis software (Durand et al., 2016). Following the example of Fulco et al. (2019), we replaced the diagonal of the output matrix with the maximum value of each element’s nearest neighbors. (The diagonal of the raw matrix is highly correlated with the number of restriction sites in each bin, rather than true contact frequency between sites within the bin.) We then averaged the contact frequencies (weighted by sequencing depth) across the aforementioned cell lines.

##### Enhancer validation

Active regulatory elements are often identified by additional sequencing assays that detect features such as bi-directional transcription (Lam et al., 2014; Tippens et al., 2020; Andersson et al., 2014; Core et al., 2014) or histone modifications (Ernst and Kellis, 2012, 2017; Boix et al., 2021). Therefore, we downloaded a set of 32,693 enhancers identified by CAGE-seq from the FANTOM Consortium (Andersson et al., 2014). In addition, we downloaded the `ChromHMM` segmentation of the genome into chromatin states within K562 cells (Boix et al., 2021). While neither dataset provides a comprehensive set of *cis*-regulatory elements, we used them to validate our detection of functional enhancers.

#### The elastic net regression model

Our enhancer-to-promoter map is derived from first principles in functional genomics. Among these is that the transcriptional activity at CREs (both promoters and enhancers) can be measured with DNase-seq or ATAC-seq, because open chromatin is permissive for the binding of transcription factors (TFs). Moreover, chromatin accessibility is not a binary property; it varies along a continuum, with progressively greater levels of accessibility thought to promote correspondingly more frequent TF binding. We therefore assume that the *degree* of chromatin accessibility at a CRE, determined by the number of sequencing reads from the element, is proportional to its transcriptional activity. In particular, the level of chromatin accessibility at a promoter is assumed proportional to gene expression. Another fundamental notion is that the chromatin accessibility of an enhancer should covary with the accessibility of a promoter it regulates: the more active an enhancer, the greater the expected transcriptional activity (accessibility) at an interacting promoter. Some of the earliest attempts to build enhancer-promoter maps were based on these same principles (Ernst et al., 2011). In addition, we leverage another key concept: that, mechanistically, enhancers boost transcription through chromatin loops that bring them into physical contact with promoters in three-dimensional space. We model these ideas with a simple equation that expresses promoter activity as a linear function of enhancer activation. The solution to this equation, which we estimate via elastic net regression, provides a set of predicted enhancers for a particular promoter. In this way, enhancers are identified by virtue of their ability to predict promoter activity, subject to two constraints. First, a sparsity constraint selects the smallest set of enhancers that accurately predict the promoter activity. Second, the elastic net models preferentially select enhancers in physical contact with the promoter, from the Hi-C data.

We model the regulation of gene expression with the following equation:

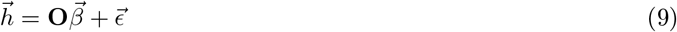

where 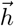 is a 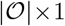 vector giving the chromatin accessibility of a particular CRE across 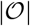 sequencing observations. Likewise, **O** is a 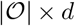 submatrix of **D**, containing only the columns corresponding to putative enhancers within a maximum distance of the modeled CRE. The inputs 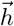 and **O** encode the number of normalized sequencing reads from CREs (measured in RPKM), primarily in DNase-seq samples. We mean-center and standardize the dependent variable, 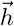, as we do the columns of **O**. We denote by 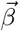 the *d* × 1 vector of enhancer coefficients, which model the effect of each enhancer on the dependent variable. Consider the *i*-th enhancer. A one-standard-deviation change in the enhancer RPKM would account for a shift of 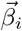 standard deviations in the RPKM of the dependent variable. As usual, we assume 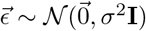, where **I** is the 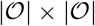 identity matrix.

Our goal is to solve equation (9) for 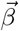, subject to several constraints. Recall the mean-centered and standardized RPKM matrix **D**. Furthermore, let 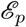 denote the set of all putative enhancers within 2 Mbp of the promoter *p*. (See below for the method used to obtain putative enhancers.) Then we solve the constrained optimization problem defined by the elastic net objective function:

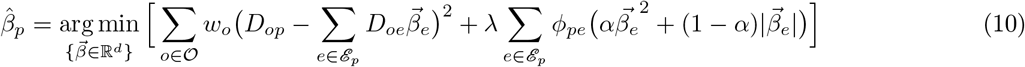

Here, 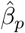 denotes the optimal enhancer coefficients for the promoter *p*. *D_or_* is the RPKM count in observation *o*, from cis-regulatory element *r*. The constant *w_o_* is a weight that normalizes the contribution of each cell type to the optimization; it is inversely proportional to the frequency of the corresponding cell type in 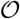. The first term in the optimization, then, is clearly the (weighted) prediction error obtained with the coefficient vector 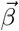. The second term, on the right, is a summation over penalties that constrain the optimization. Progressively larger values of the regularization parameter λ yield sparser solutions, with fewer nonzero coefficients in 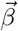. Conservatively, we set λ to the largest value that achieves an out-of-sample predictive performance (on three-fold cross-validation data) within one standard error of the best achievable with this model. The parameter α controls the relative balance between the lasso (*α* = 0) and ridge (*α* = 1) components of the elastic net regression; we vary this parameter to tune the density of the map. We set *α* = 0.04 to generate the enhancer-to-gene map in this work (corresponding to the starred points in Figure 2E). Finally, *ϕ_pe_* is the penalty imposed for including enhancer e in the regression model for promoter *p*. As the inverse of the chromatin contact frequency between *p* and *e*, *ϕ_pe_* encodes the prior belief that functional enhancers require physical interaction with the promoter.

#### The Hi-C prior

The Hi-C prior, encoded in the elastic net penalties, incentivizes the model to reject most distal interactions, except those with unique predictive value. Unlike DHS activity, which is highly variable across cell-types, the 3D structure of chromatin is less variable (Rao et al., 2014), allowing us to employ the same penalty across cell types. To compute the penalty, *ϕ_pe_* in Equation (10), we refer to the normalized Hi-C contact matrix obtained as described above. Given that matrix, we first define unnormalized penalties as:

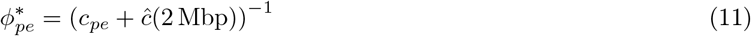

where *c_pe_* is the normalized contact frequency between promoter *p* and enhancer *e* from the corresponding bin in the contact matrix. In addition, *ĉ*(*d*) is the estimated contact frequency between two loci separated by a genomic distance *d*, determined from a power law model fit to the contact matrix via ordinary least squares:

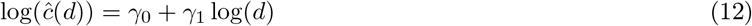

We added the term *ĉ*(2 Mbp) to regularize the penalties. Otherwise, some penalties would be undefined, corresponding to bins in the (sparse) contact matrix with zero reads (Fulco et al., 2019). The penalties are then normalized such that they sum to the number of predictor variables:

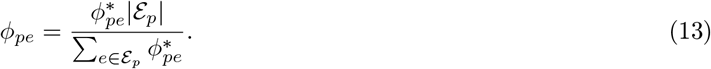

#### Identification of promoters and putative enhancers

Solving the optimization problem in Equation (10) is predicated on identifying candidate CREs (promoters and enhancers) for each gene of interest. Often, promoters and enhancers are identified by the presence of histone modifications; for example, the H3K4me3 mark is often seen on active promoters and H3K27ac on active enhancers. Others have used an array of such marks to decode chromatin to nucleosome resolution, thereby annotating regions as one of several states (Ernst and Kellis, 2012, 2017). For our purposes, however, we require a cell-type agnostic definition, and preferably one for which only chromatin accessibility data are needed (without additional chromatin mark data). Therefore, we use the following definitions:

1. **Promoter**: Any DHS in the atlas of open chromatin sites within 500 bp upstream or downstream from any transcription start site is considered a promoter for the corresponding gene. We use the database of transcription start sites for all annotated genes from Ensembl (Cunningham et al., 2019) in GRCh37 coordinates.
2. **Candidate Enhancer**: Any non-promoter DHS in the atlas of open chromatin sites within 2 Mbp of any promoter.
3. **Putative Enhancer**: Any candidate enhancer whose activity can be accurately predicted from nearby, non-overlapping DHSs.

Given an atlas of open chromatin sites, promoters and *candidate* enhancers are defined purely in terms of genomic neighborhood. *Putative* enhancers, however, which we consider as sites that may regulate gene expression, are only a subset of those DHSs in the relevant genomic neighborhood. We hypothesized that if a DHS is a true enhancer, then its activity should be coordinated with that of other enhancers, and it should influence promoter activity. That is, the activity of the enhancer itself should be predictable from nearby DHSs. Therefore, we define a model analogous to Equation (9):

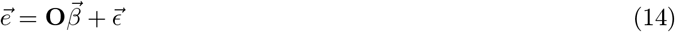

Here, 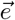 is the activity of a candidate *enhancer* and **O** is the RPKM matrix at DHSs within a particular distance of the enhancer being modeled (we use 250 kbp to minimize computational costs). As with Equation (9), the problem is under-constrained and we estimate 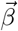 using an elastic net objective function:

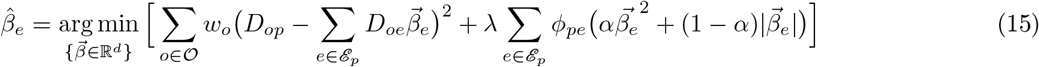

For the enhancer models, we set the elastic net parameter *α* to its common default (*α* = 0.5). Using this formulation, we built an elastic net model for each candidate enhancer, and computed its coefficient of determination (*R*^2^), to measure how well its DHS activity could be predicted from nearby *cis*-regulatory elements. We considered only DHSs with *R*^2^ > 0.7 (approximately 75% of candidate enhancers) as putative enhancers in our promoter models (see the main text for the justification of this threshold).

#### Linking enhancers to promoters

Enhancers are linked to promoters via the solution to the elastic net regression problem posed in Equation (10). We assign enhancers to promoters if the coefficient *β_pe_* > 0; putative enhancers with coefficient *β* ≤ 0 are rejected. A negative coefficient would indicate the DHS is a silencer. While silencers may be an important component of gene regulation (Jayavelu et al., 2020), computational predictions of silencer-gene interactions cannot be validated with existing data. Absent the validation data, we discard these interactions to avoid linking false-positive DHSs to genes in our map.

#### Identifying validated interactions

We used two approaches to validate that our enhancer-to-gene map accurately identifies enhancer-gene interactions. We compared the output of our map (and other enhancer-to-gene maps), first against a set of known enhancers from functional genomics experiments, and second against variant-gene interactions discovered from eQTL association studies.

We compiled a set of functional genomics experiments that use CRISPR to identify enhancer-gene interactions (Laurell et al., 2012; Claussnitzer et al., 2014; Prager et al., 2014; Hilton et al., 2015; Claussnitzer et al., 2015; Canver et al., 2015; Thakore et al., 2015; Diao et al., 2016; Korkmaz et al., 2016; Fulco et al., 2016; Xie et al., 2017; Klann et al., 2017; Simeonov et al., 2017; Gupta et al., 2017; Gasperini et al., 2019; Schraivogel et al., 2020; Reilly et al., 2021), including four high-throughput assays that constitute the majority of these data (Fulco et al., 2016; Gasperini et al., 2019; Schraivogel et al., 2020; Reilly et al., 2021). Taken together, these experiments include a total of 866 unique, well-validated enhancer-gene interactions. We removed from this list any interactions in which the target gene does not have a promoter in our atlas of open chromatin sites (1 out of 866), as these could not be identified in our map. We further removed any interactions in which the enhancer is within 300 bp of a promoter for the gene (34 out of 866, as these interactions are automatically included in our map). Finally, to assure a fair comparison between our method and others in the literature, we only include interactions with genes targeted by at least one enhancer from each alternative method in Figure 2 (FOCS, JEME, EpiMap, PEGASUS, and ABC). After filtering, 735 enhancer-gene interactions targeting 473 unique genes are included in the validation data. Three of these experiments (Gasperini et al., 2019; Schraivogel et al., 2020; Reilly et al., 2021) target candidate regulatory elements specifically, with known genomic coordinates; for these cases, we require an enhancer targeting the same gene to overlap with those coordinates, in order to consider the enhancer-gene interactions validated. For the other methods, we considered the specific assay (e.g., CRISPR inhibition or CRISPR activation) and the genomic distance over which the disruption (centered at the sgRNA) is likely to function, resulting in an allowable ‘buffer distance’ between the assay target and the enhancer in our atlas (see Table S6). If an enhancer targeting the same gene is within this buffer distance, we consider the interaction to be validated. All of the interactions identified from literature—and whether they are observed in the enhancer to gene map—are reported in File S1.

We also used eQTL studies from GTEx, restricting our analysis to cases in which the eQTL can be fine-mapped to single-variant resolution (GTEx Consortium, 2020). We downloaded eQTL studies fine-mapped with `CAVIAR` from the GTEx portal (GTEx Consortium, 2021), and filtered the list to only include variants with a PPA > 0.9. We used `liftOver` to convert the coordinates from GRCh38 to GRCh37; we removed any variants that did not map one-to-one between genome builds. Additionally, we removed variants greater than 1 Mbp from the associated gene (because this is the maximum distance between variants and transcription start sites tested by GTEx; any distances nominally above this are likely due to mismatches between genome builds). We further removed any variant-gene pair predicted by VEP, as these interactions are less likely to be enhancer-mediated, and interactions within 300 bp of any promoter for the target gene. As with the functional genomics assays, we only include interactions that target genes with at least one enhancer in each method we assess (FOCS, JEME, EpiMap, PEGASUS, ABC), ultimately resulting in 9541 variant-gene interactions that include 9208 unique variants and 5268 unique genes. The variant-gene interactions are considered to be in our map if the causal variant resides within 300 bp of an enhancer that we link to the target gene. All of the variant-gene interactions and their validation results are described in File S2. In Figure 2, we plot the validation rate measured against likely enhancers, whose regression models must satisfy *R*^2^ > 0.84 (an even more strict criterion than we used to define putative enhancers; see *Identification of promoters and putative enhancers*). Figure S9 plots the validation rate against all the variant-gene pairs described here.

In order to compare the performance of our enhancer-to-gene map to alternatives in the literature, we computed both the validation rate (i.e., the recall, or fraction of observed interactions that are recovered by each map) and the enhancer density (i.e., the average length of DNA, in kbp, predicted to regulate a target gene). Note that the enhancer density depends strongly on the set of genes used for measurement. For example, we noted that in our enhancer-to-gene map, protein-coding genes tend to have significantly greater enhancer density than non-coding genes. Therefore, to compute the enhancer density, we compute the mean length of regulatory DNA for genes that are common to all the methods under comparison (i.e., are linked to at least one enhancer in each method) and have one validated enhancer-gene pair (either in the CRISPR studies or in the fine-mapped GTEx interactions). Frequently, in classification problems, the precision of the classifier, (i.e., the fraction of predicted enhancers that actually regulate the target gene), is also computed. In the case of enhancer-to-gene maps, however, computing precision is not possible. There is no unbiased sample of true negatives, which do not regulate the expression of a gene in any cell type or any context. We therefore use enhancer density as a proxy and assume that a sparser map has higher precision.

#### Linking variants to genes

The primary component of the variant-to-gene map is the enhancer-to-gene map described above. We link variants to genes by way of this map if they lie within a DHS (enhancer or promoter) predicted to regulate the target gene, or if they are within 150 bp of such a DHS. If there are no DHSs within 150 bp of a given variant, then we assign that variant to any DHS within 300 bp of the variant.

In addition to the enhancer-to-target-gene map, we link variants to genes using: (1) close proximity to the transcription start site; (2) predictions from VEP; (3) fine-mapped GTEx eQTLs (PPA > 0.9); or (4) validated enhancer-gene interactions from a functional genomics assay, using the same assays as we used to validate our enhancer-to-gene-map (see *Identifying validated interactions*). For the proximal variants, we link them to a gene if they fall within 2000 bp upstream or 500 bp downstream of a known transcription start site. Using VEP, we link variants to genes if they are likely to affect the protein product, mature transcript, or transcript splicing. Specifically, we link variants to genes if the variants have any of the following predicted consequences: ‘transcript ablation’, ‘splice acceptor variant’, ‘splice donor variant’, ‘stop gained’, ‘frameshift variant’, ‘stop lost’, ‘start lost’, ‘transcript amplification’, ‘inframe insertion’, ‘inframe deletion’, ‘missense variant’, ‘protein altering variant’, ‘incomplete terminal codon variant’, ‘start retained variant’, ‘stop retained variant’, ‘synonymous variant’, ‘splice region variant’, ‘coding sequence variant’, ‘mature miRNA variant’, ‘5 prime UTR variant’, or ‘3 prime UTR variant’.

### THE STATISTICAL FINE-MAPPING MODEL

#### Model Definition

The core of our framework is a Bayesian regression model, an extension of previous statistical fine-mapping methods (Servin and Stephens, 2007; Hormozdiari et al., 2014; Benner et al., 2016) that set the foundation for this work. For simplicity, we consider the common case of a GWAS on a single trait carried out in a population of homogeneous ancestry. The generalization of the model to multi-ancestry data is straightforward and is provided in a subsection below. Denote by *m* the number of variants in the GWAS. *A priori*, we know that most of these *m* variants are not causal modifiers of the phenotype under study, even if they are correlated with the phenotype via LD with causal polymorphisms. Let 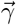 be a latent indicator vector that encodes whether each variant *i* is causal 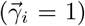 or not 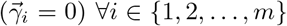. Each realization of 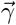 is a ‘causal model’: it encodes the hypothesis that a particular genome-wide set of variants is responsible for the observed GWAS data. Denote by 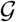 the set of all genes in the human genome which are directly affected by a causal variant. Our goal is to compute two crucial quantities, given the data observed in the GWAS: 1.) the probability that each variant is causal, 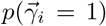; and 2.) the probability that any gene *g* is targeted by a causal polymorphism, 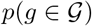. Call these quantities the variant and gene posterior probabilities of association (PPAs), respectively. Following the standard in fine-mapping methods, we model the GWAS with the Bayesian linear regression

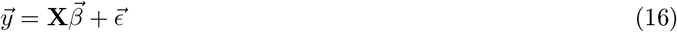

where 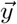 is a *n* × 1 column vector of the phenotype observed across all *n* individuals in the GWAS cohort; **X** is the *n* × *m* genotype matrix, for *m* variants; 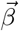 is the *m* × 1 column vector of (unknown) effect sizes for each variant; and 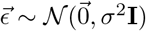, with **I** the *n* × *n* identity matrix. The matrix **X** is encoded, and its columns standardized, following convention; for instance, see (Chen et al., 2015; Yang et al., 2011; Price et al., 2006; Yang et al., 2015, 2010). Without loss of generality, we assume that **X** and 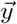 have already been corrected for non-genetic covariates and population stratification (Price et al., 2006).

The Bayesian character of the regression—and the inference of causality for each variant—enter through a prior on the effect sizes 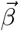. As in other models (Chen et al., 2015; Benner et al., 2016), we assume 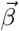 follows a multivariate normal distribution, given the causal indicator 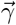:

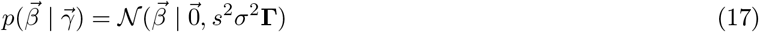

where **Γ** is the *m* × *m* diagonal matrix with 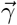 along the diagonal. We denote by s the standard deviation of the causal effect sizes, as a fraction of the standard deviation of the phenotype, *σ*. Note that s is an input parameter of the model; it should be set according to the polygenicity of the phenotype (see Note S5, for which Notes S3–S4 provide relevant background).

We will need to evaluate the probability of observing the GWAS phenotypes 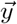, given the genotypes and a hypothesized set of causal variants: 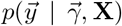. However, the raw, individual-level data (**X**, 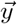) are generally unavailable. Instead, as shown previously (Yang et al., 2012; Chen et al., 2015; Benner et al., 2016), we can express the desired probability in terms of the GWAS z-scores 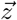 (i.e., Wald statistics) and the variant correlation matrix **R** = *n*^−1^**X**^┬^**X** (which *are* available):

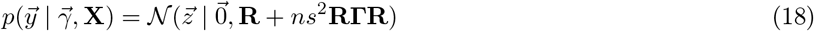

We will leverage this result (below) to compute the probability of each causal model, given the data, which is the central calculation in statistical fine-mapping. This formulation also unifies the treatment of quantitative and binary phenotypes: given the z-scores, equation (18) holds for both continuous phenotypes and case-control study designs (Yang et al., 2012; Kichaev et al., 2014; Chen et al., 2015; Benner et al., 2016). For completeness, we derive this equation, and show how it can be efficiently computed, in Notes S3–4 and S6.

The matrix **R** can be estimated from a large reference sample; it is the Pearson correlation matrix between the diploid genotypes at each pair of variants. We computed **R** directly from the individual-level genotype data available from the UK Biobank (Bycroft et al., 2018), and we restricted our GWAS compendium to studies in European cohorts. It is straightforward to show that each element of **R** is equal, in expectation, to the *ϕ* correlation between variants, assuming the allele frequencies are in Hardy-Weinberg equilibrium. (Recall that the *ϕ* correlation is the Pearson correlation computed on *haplotypes*, so that genotypes are encoded as {0, 1}.) Therefore, **R** is precisely the same matrix underlying the LD partition algorithm.

In order to compute the PPAs for variants or genes, we require an expression for 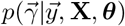, where ***θ*** denotes the parameter vector of the prior distribution 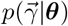 (see below). Let Γ denote the set of all 2^*m*^ possible realizations of the indicator vector (i.e., the set of all possible models encoding the genetic architecture of a trait). Then, leveraging the chain rule of conditional probability and the law of total probability, we have

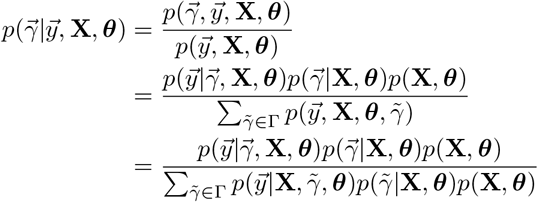

(using the dependence structure of the model)

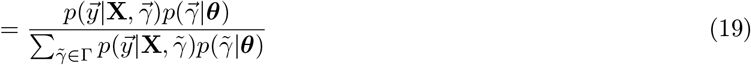

We tested two alternative formulations of 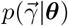, the prior distribution on the causal indicator 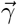. The first is a uniform prior, 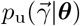, encoding the assumption that all variants are equally likely, *a priori*, to be causal:

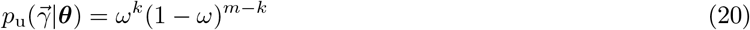

where ***θ*** = [*ω*], *ω* is the prior probability that any single variant is causal, and 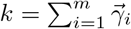 (i.e., the number of causal variants). The parameter *ω* may be directly supplied, as a model input, or it can be learned from the data, via expectation maximization (EM; see below). While this prior is oversimplified, it is often chosen for convenience.

The second of our two priors is novel: it encodes different prior probabilities for variants as a function of their biochemical consequence. We annotate each variant in a GWAS according to its predicted biochemical effect, using a waterfall logic to assign the most severe consequence to variants with multiple effects (see Table S5). Let 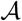 denote the set of variant annotations, with 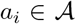 the annotation of the *i*-th variant. Denote by 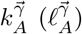 the number of causal (noncausal) variants in 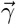 with annotation *A*, and let *P*(*C*|*A*) represent the probability that a variant with annotation *A* is causal. Then the ‘biochemical’ prior on the causal indicator is given by:

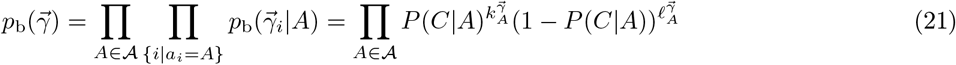

The probabilities 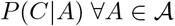 are learned directly from a compendium of GWAS data, using the EM algorithm (see below).

Given the foregoing definitions and results, we may finally define the variant and gene PPAs. First, the variant PPAs, for all variants *i* ∈ {1,…, *m*}, are given by:

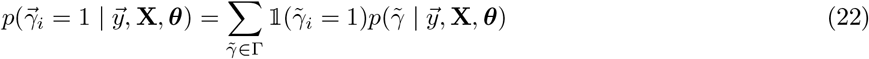

where 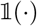 is the indicator function. Similarly, the gene PPAs are:

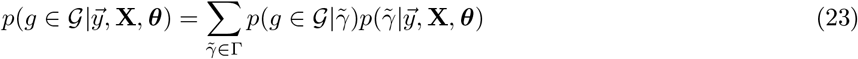

In equations (22) and (23), the term 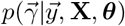 may be computed with either the uniform or biochemical prior, depending on the desired calculation.

#### Derivation of Variant PPAs

Consider equation (19). The posterior probability of a causal model entails a summation over all 2^*m*^ possible models, which is intractable if attempted directly. (For a typical GWAS, *m* ≈ 10^7^.) However, given a partition of the human genome into uncorrelated LD blocks, the summation can be divided into smaller, independent subproblems, which can be solved with a stochastic search algorithm (Hans et al., 2007; Speed and Tavaré, 2011; Benner et al., 2016).

Recall that 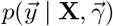 follows a multivariate normal distribution with covariance matrix **C** = (**R** + *ns*^2^**RΓR**). The (*i*, *j*)-th element of **C** is

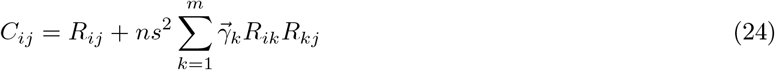

By definition, variants have negligible correlation outside LD blocks, so *R_ij_* ≈ 0 ∀(*i*, *j*) such that variants *i* and *j* fall in different LD blocks. Therefore, the only nonzero terms in the summation over *k* in equation (24) occur when variant *k* resides in the same LD block as both *i* and *j*, which implies that *i* and *j* are in the same LD block. Therefore, 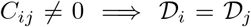, where 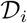 denotes the LD block of the *i*-th variant. Consequently, we can factorize 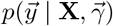 into the product of independent normal distributions segmented on the LD blocks in the human genome:

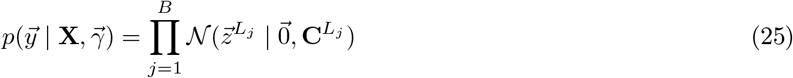

where B is the number of LD blocks in the genome; *L_j_* denotes the *j*-th LD block; and 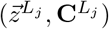 are the partitions of 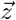 and **C** corresponding to the variants in 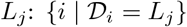. Since the probability 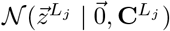 can be solved efficiently with shotgun stochastic search (SSS), as in Benner et al. (2016), we have decomposed the intractable genome-wide calculation into the product of tractable computations. We use this decomposition to rewrite the probability of a causal model, 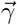.

First, we leverage equations (18) and (19) to write the probability of 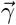 in terms of the GWAS summary statistics:

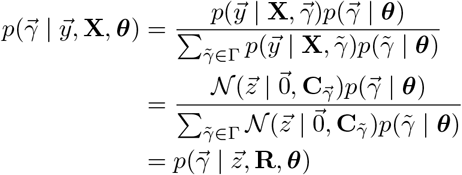

where 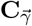 denotes the covariance matrix for the causal model 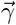. Then, we apply our decomposition to get a tractable expression for 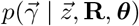. We start with:

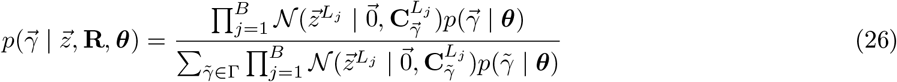

We rewrite the denominator of equation (26):

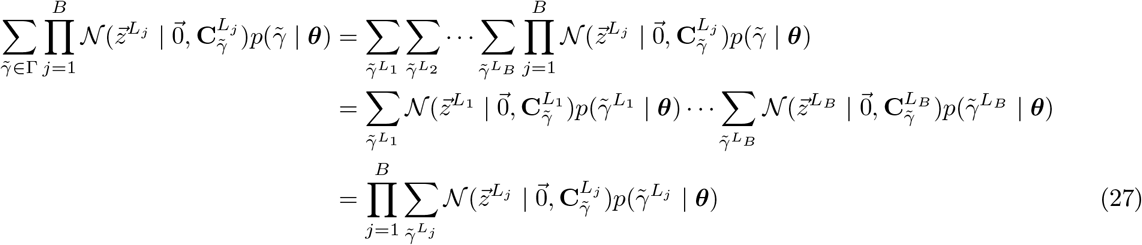

Substituting equation (27) into (26), we obtain

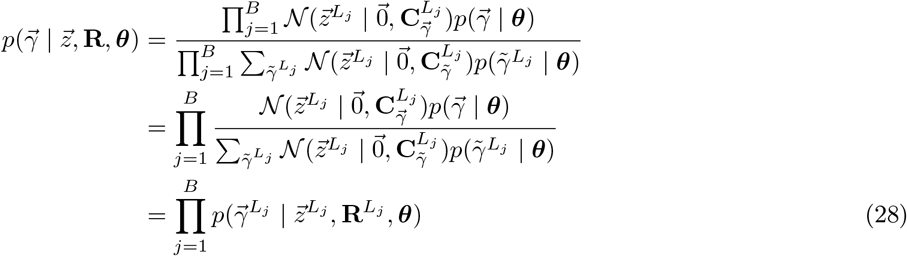

Equation (28) is tractable with SSS. To simplify notation, let 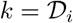 denote the index of the LD block in which the *i*-th variant resides. We can derive a computationally tractable formula for the PPA of the *i*-th variant:

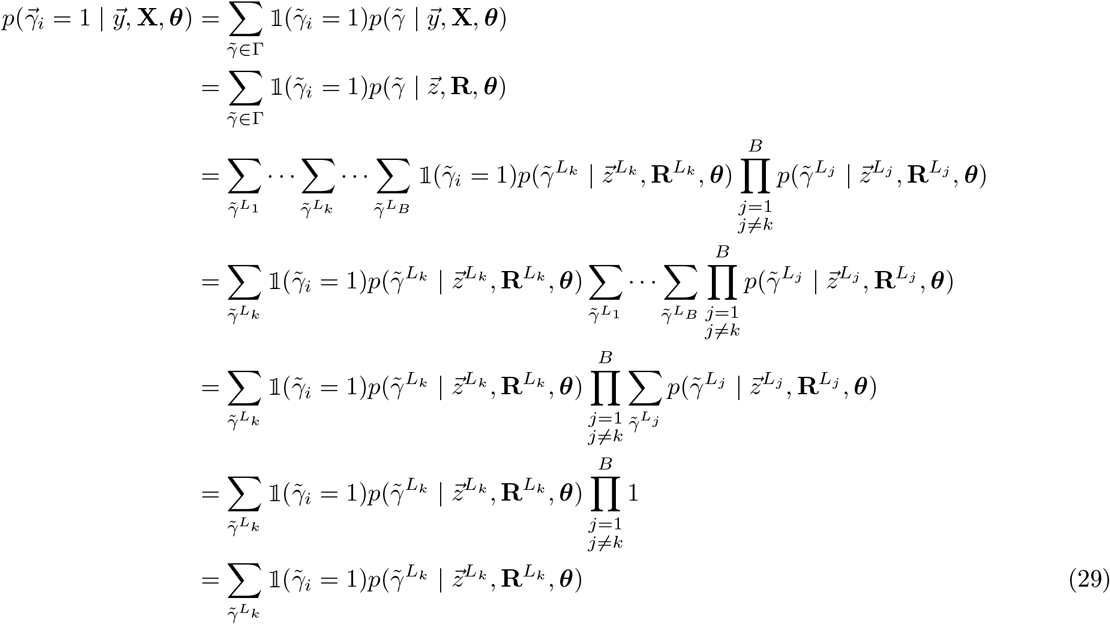

#### Derivation of Gene PPAs

Using the results from the previous section, and our variant-to-gene map, we can write a computationally tractable expression for the PPA of any gene *g*:

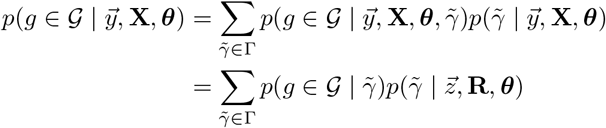

Let 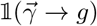 denote the indicator function whose value is unity if the model 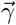 contains at least one causal variant linked to gene *g* by our variant-to-gene map, and zero otherwise. In addition, let the sequence [*V*_1_,…, *V_b_*] denote the LD blocks containing any variant linked to *g*. Further, let 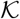 be a set containing the indices of these linked blocks. Lastly, let 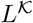 be the concatenation of the blocks [*V*_1_,…, *V_b_*], which need not be contiguous. Then we have 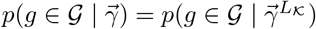 and:

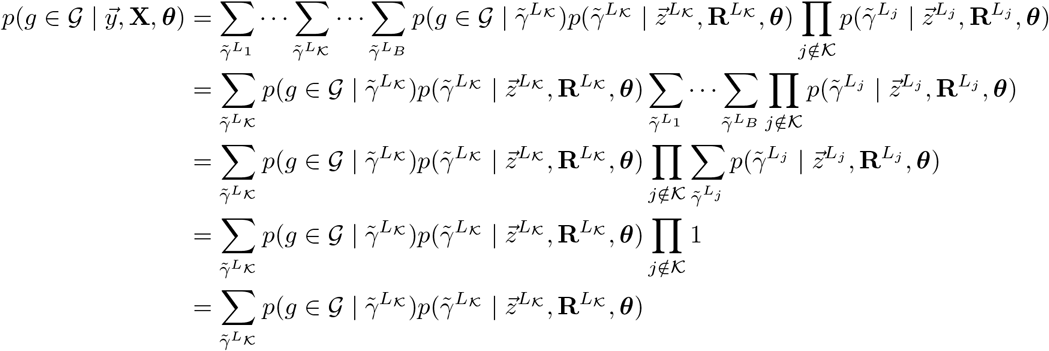

(assuming correctness of our variant-to-gene map)

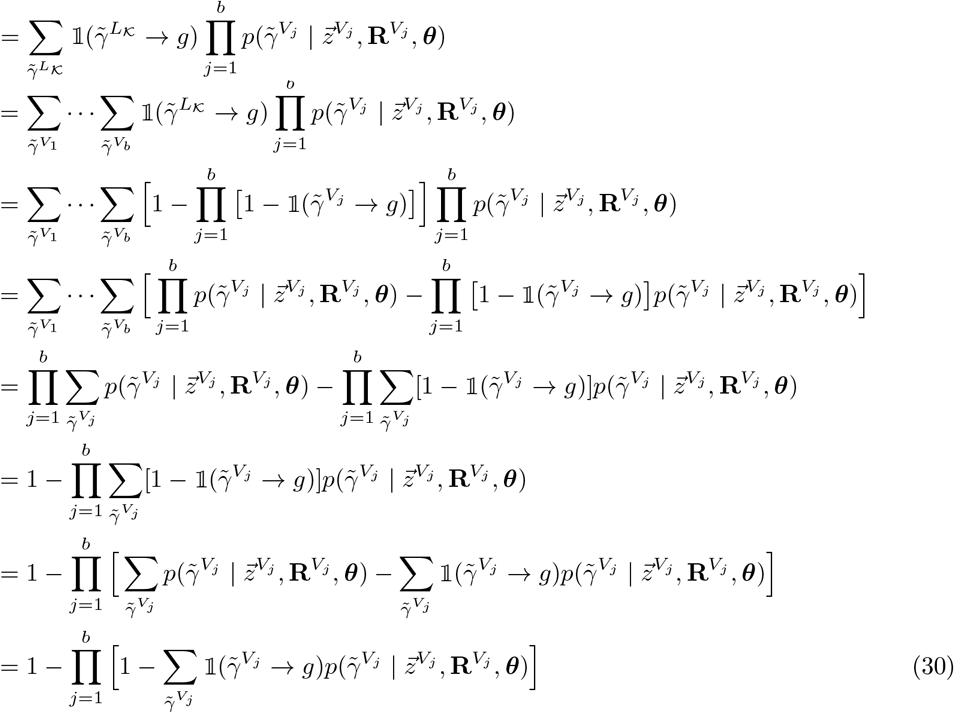

#### Computing PPAs with Expectation-Maximization

##### The Expectation-Maximization Algorithm

The two preceding sections establish tractable formulae to evaluate the PPA for any gene or variant in the genome. The equations, however, depend on a vector of unknown parameters, ***θ***. In a GWAS, we observe the phenotypes of the study population, with the aim to infer from the genotypes the latent variables 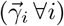 that mediate the phenotypes. In particular, we aim to compute the probability distribution over 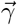, but that distribution depends on the unknown parameters ***θ***. The expectation-maximization (EM) algorithm (Demster et al., 1977; Moon, 1996) is the classical method in this setting, whereby one desires to estimate the parameters of a distribution (***θ***) in the context of latent variables 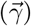. With the EM algorithm, we can learn ***θ*** directly from the data, by finding the parameters that maximize the joint probability of the observed data and the latent genetic architecture 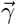, over all possible realizations of 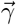. We then enter the learned value of ***θ*** into equations (29) and (30), to compute the variant and gene PPAs. To implement the EM algorithm in our setting, we must iteratively:

1. Compute the expected value of ln 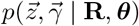 over the probability distribution 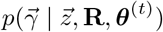, using the current estimates of the parameters, ***θ***^(*t*)^:

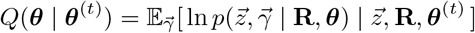
2. Replace the current estimates of the parameters with the values that maximize the expectation computed in the previous step (i.e., the so-called *Q* function):

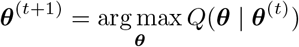

The EM algorithm is deemed to reach convergence when ||***θ***^(*t*+1)^ – ***θ***^(*t*)^|| < *ϵ*, for some prescribed value of *ϵ*. In the following subsections, we show how to apply this framework to learn the parameter(s) of the uniform and biochemical priors. In every case we consider, the *Q* function we optimize is strictly concave, so the EM algorithm is guaranteed to converge to the global optimum.

##### Learning the Parameter of the Uniform Prior

In the case of the uniform prior, ***θ*** = *ω* and the *Q* function is:

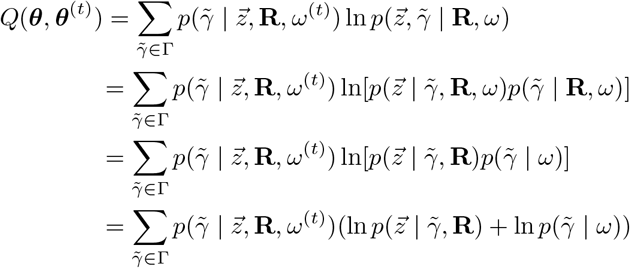

At each iteration, we must set *ω*^(*t*+1)^ to the value that maximizes *Q*(*ω* | *ω*^(*t*)^). To this end, we first note that:

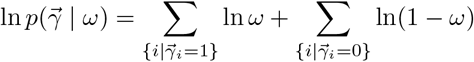

In addition, the term ln 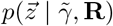 does not depend on *ω*, so we can omit it from the optimization. Therefore, it suffices to optimize:

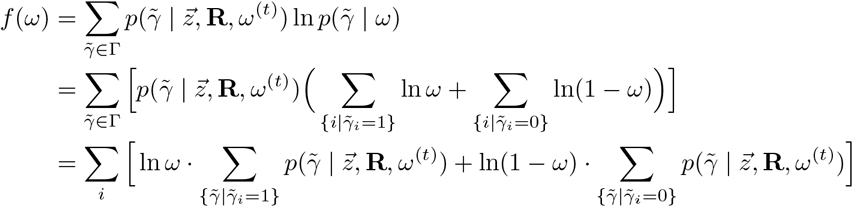

We recognize 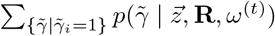 as the PPA for the *i*-th variant, computed at iteration *t*. We denote this as 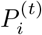. Similarly, the term 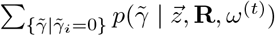 is nothing more than 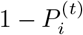. Therefore, we have:

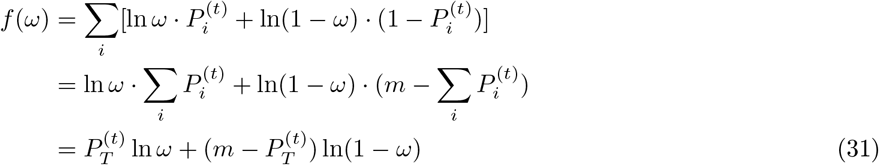

where we have defined the sum of all variant PPAs:

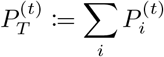

Recall that *m* denotes the number of variants in the GWAS. In order to maximize *f*(*ω*), we set its derivative to zero:

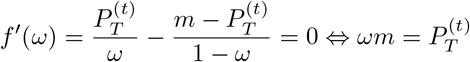

and we confirm that *f*(*ω*) is strictly concave:

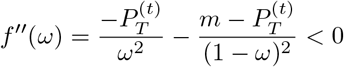

Therefore, *f* is maximized by setting the parameter at each iteration to the mean variant PPA:

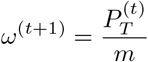

At each EM iteration, we compute all the variant PPAs, using the current estimate of *ω*, then update *ω* to the mean of those PPAs. We repeat this procedure until the estimate of *ω* converges. Lastly, we apply the ultimate estimate of *ω* to compute the variant and gene PPAs from equations (29) and (30), respectively. Unlike the case of the biochemical prior (see below), there is no information to share across traits in the uniform prior. That is, there is no relationship between the value of *ω* in one trait compared to another. Therefore, the parameter of the uniform prior is computed separately for each trait.

#### Learning the Parameters of the Biochemical Prior

##### Learning *θ* from a single trait

For clarity of exposition, we begin with the task of estimating ***θ*** in the biochemical prior from a single trait, although in practice the parameter vector is always learned from a compendium of traits. Suppose we have multiple annotations *A*_1_,…, *A_K_*, where each annotation *A* is a disjoint subset of variants with a particular biochemical consequence. (We use *A* to denote an arbitrary annotation, and *A_j_* to denote a specific annotation, corresponding to the *j*-th type of biochemical consequence, for *j* ∈ {1,…, *K*}.) For each annotation *A*, we would like to find a prior *ω_A_*. That is, ***θ*** is now the parameter vector ***θ*** = [*ω*_*A*1_,…, *ω_A_K__*]. We define *m_A_*:= |*A*| and

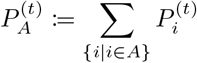

For each annotation *A*, *m_A_* is the number of variants with that annotation, and 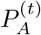 is the sum of their PPAs. We follow the calculation of *f* in equation (31), but now we must keep track of the prior for each variant. If we let *ω_i_* denote the prior for the *i*-th variant, then we have

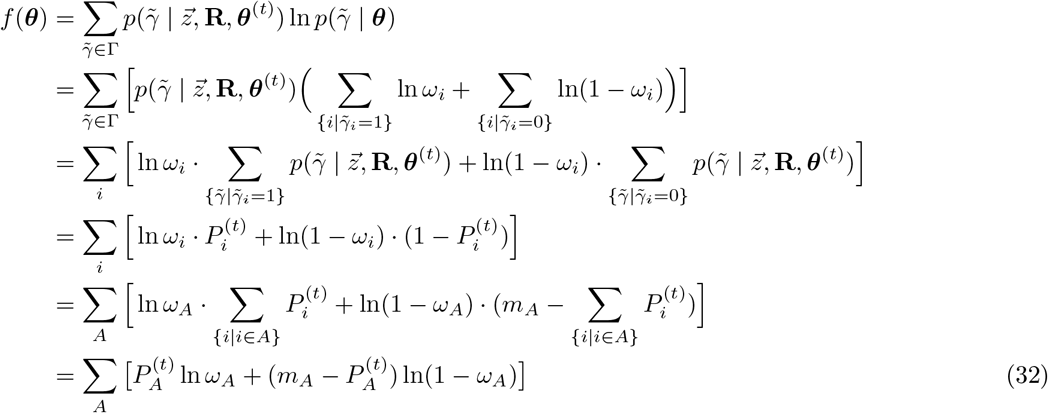

Here, *f*(***θ***) is a sum of terms, each of which depends on a single annotation. The gradient is simply

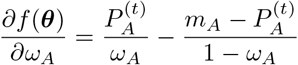

In addition, the Hessian is diagonal and negative definite, implying *f*(***θ***) is strictly concave, with each element given by:

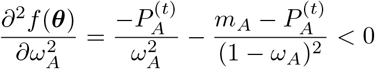

Therefore, we can maximize *f*(***θ***) by setting the gradient to zero, and updating each prior *ω_A_* to the mean PPA of all variants with the corresponding annotation:

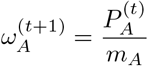

##### Learning *θ* from multiple traits

In practice, a GWAS on a single trait may not suffice to estimate the prior for variants of multiple biochemical types. We therefore extend our framework to optimize the parameters over multiple GWA studies, by sharing the parameter values across traits. However, different phenotypes have distinct genetic architectures and are mediated by varying numbers of causal variants. We cannot simply assume that the prior probability for each variant is the same across traits. Instead, we assume only that *relative* prior probabilities are shared across traits. For instance, suppose that a missense variant is 100 times more likely to be causal, *a priori*, than a variant with unknown biochemical consequence. We assume this constant of proportionality holds across all traits, while allowing the priors themselves to scale with the polygenicity of each trait. We use Bayes factors (see below) to share the relative prior probabilities across traits.

Suppose we have biochemical annotations 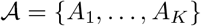 and a compendium of phenotypes 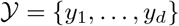. Variants with no predicted biochemical effect are unannotated. Now consider some annotation *A*. Let *P*(*C* | *A*) denote the probability that a variant is causal for a specific trait, given that it has annotation *A*. Then the odds version of Bayes theorem states:

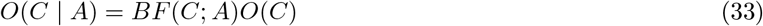

where the Bayes factor *BF*(*C*; *A*) is

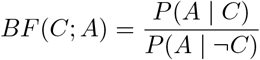

To see this, note that

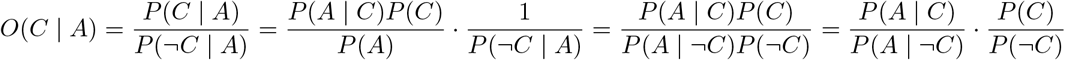

Let *σ*(*x*) denote the standard logistic function (i.e., the inverse of the logit function), and notice that:

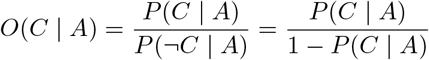

Therefore,

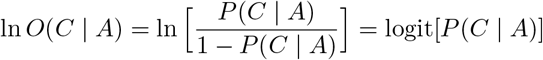

so that

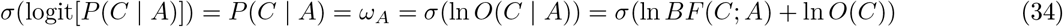

We introduce a change in variables, in order to express the forthcoming terms relative to unannotated variants. Define

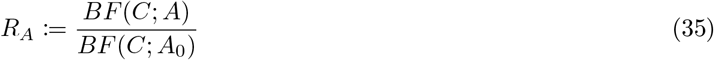

and

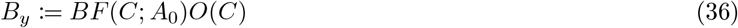

where *A*_0_ denotes that a variant is unannotated. *R_A_* is assumed constant across traits, but *B_y_* will differ for each phenotype *y*. Observe that

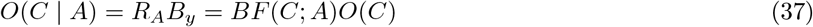

Since *R*_*A*_0__ = 1, this implies that *B_y_* = *O*(*C* | *A*_0_). Writing in terms of the new variables (*R_A_*, *B_y_*), the prior probability that a variant with annotation *A* is causal, in trait *y*, is:

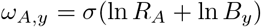

To simplify notation, we define

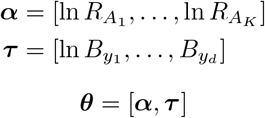

Therefore

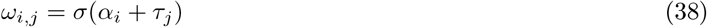

denotes the prior probability of a variant with the *i*-th annotation, in the *j*-th trait. We use this result to write the optimization function *f* for the EM algorithm in the case of multiple traits.

To extend the *Q* function for the multi-trait scenario, we simply concatenate the data from a compendium of GWA traits. We treat the concatenated traits as if they were an elongated sequence of uncorrelated LD blocks from a single study, but allowing the *τ* parameters to differ between traits. Therefore, the *Q* function is

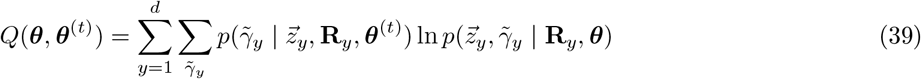

where 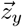 and **R**_*y*_ are the z-scores and correlation matrix for trait *y*; 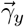 is the genetic architecture (indicator vector of causal variants) of the trait. The inner summation is identical to the *Q* function we optimized for a single trait in the preceding section. Therefore, we substitute equation (38) into formula (32) to obtain the function to optimize. Let 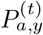 be the sum of variant PPAs in trait *y* from annotation *a*, with *m_a,y_* the number of corresponding variants. We also define 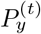 to be the sum over the PPAs of unannotated variants in trait *y*, with *m_y_* the number of these variants. Then we may write the function to optimize as:

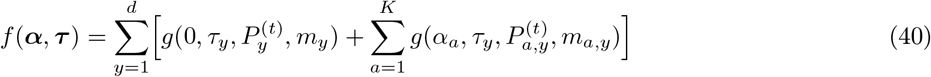

where

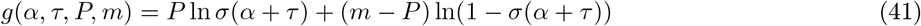

In Note S7, we show that maximizing *f*(***α***, ***τ***) minimizes the cross-entropy (discrepancy) between the prior and posterior probabilities of association for variants. That is, at each EM iteration, the prior for each type of variant is set, as closely as possible, to the average PPA for variants of that type. Moreover, we reiterate that our optimization function in equation (39) is a sum of the *Q* functions for the single-trait scenario, which we previously proved are strictly concave. Any nonnegative, nonzero sum of strictly concave functions is itself strictly concave (Boyd and Vandenberghe, 2004); therefore *Q*(***θ***, ***θ***^(*t*)^), and thus *f*(***α***, ***τ***), are strictly concave (i.e., they have a single global maximum). Therefore, we are guaranteed to find the unique maximum of *f*(***α***, ***τ***) with a Newton gradient method. We use this approach to find the maximum of f at each iteration of the EM algorithm, until it converges.

The result of the EM algorithm for the multi-trait case is the optimal vector 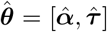. From this result, we compute the parameters of the biochemical prior, with equation (38). Given the values of the prior, we can finally evaluate the variant and gene PPAs with equations (29) and (30).

#### Extension to Multi-Ancestry GWAS

We show that the genome-wide statistical fine-mapping framework can be readily extended to multi-ancestry GWAS. Suppose that we have a collection of GWA studies on the same trait, across *p* populations from potentially distinct ancestries. Let 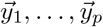 and **X**_1_,…, **X**_*p*_ denote the phenotypes and genotype matrices in these populations. Likewise, let 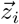 and **R**_*i*_ be the z-scores and correlation matrix from the *i*-th population. In this setting, our requirement is to compute 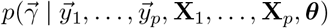. We leverage our ‘universal’ partition of the human genome into LD blocks that are mutually uncorrelated in all populations 1,…, *p*. Denote by *U* the number of universal LD blocks across the genome, and let *L_j_* represent the *j*-th universal block ∀_*j*_ ∈ {1,…, *U*}. Then we have the universal factorization

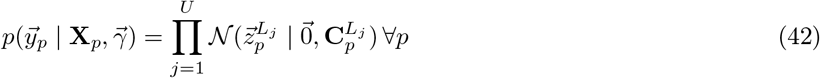

where **C**_*i*_ is the covariance matrix in population i, and our notation follows that in equation (25). Equipped with this universal partition, and following the procedures introduced in the single-population scenario, we have:

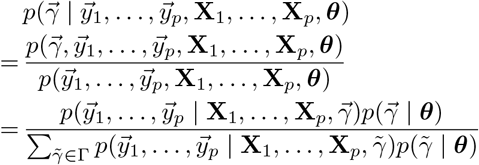

(using the independence of GWA studies)

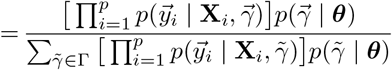

(using the universal LD partition)

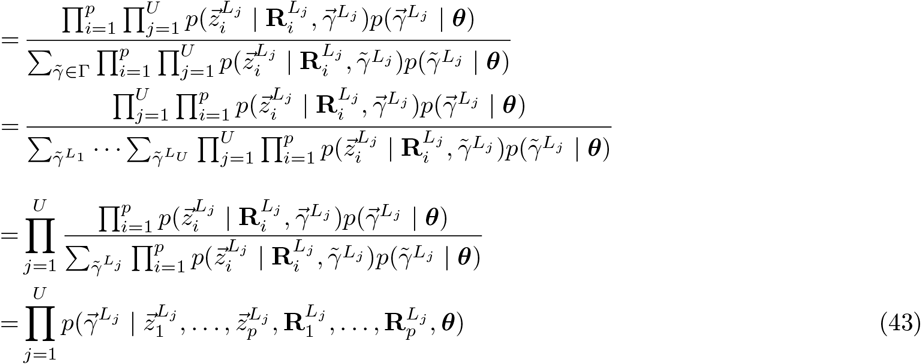

Equation (43) is tractable; it merely involves a product of the same multivariate normal probabilities we compute for the single-population case.

Given equation (43), it is easy to see that the PPA for the *i*-th variant, in a multi-ancestry GWAS, is:

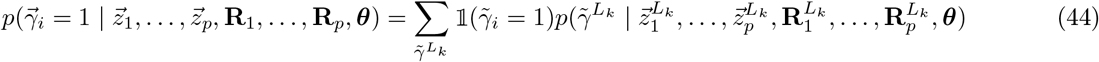

where *k* is the index of the LD block containing the *i*-th variant, as in equation (29). Similarly, using the same notation as in equation (30), the gene PPAs are given by:

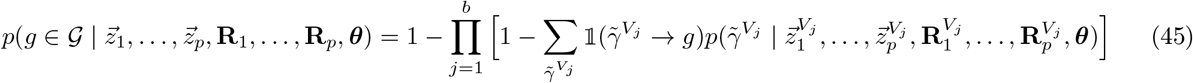

Finally, note that the parameter vector ***θ*** need not be learned directly from the multi-ancestry GWAS. Indeed, even for a GWAS on a homogeneous population, we may apply the parameters of the prior learned previously from an independent compendium of GWA studies.

#### Gene Clusters

In order to prioritize putative causal genes from among those that are targeted by a common set of causal variants, we computed ‘gene clusters’. After calculating the gene PPAs, we considered all genes with a PPA in excess of a threshold, which we set somewhere in the range [0.01, 0.1], depending on the distribution of gene PPAs for the trait. (For traits with fewer confidently implicated genes, we set the threshold lower, to ensure that all relevant genes were included in the clustering.) The genes with suprathreshold PPA were included as the nodes in a weighted, undirected graph, whose edge weights represent the extent to which connected genes are implicated by the same variant(s). More formally, consider two arbitrary genes, represented by *g*_1_ and *g*_2_. Let the indicator function 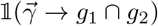 denote whether the configuration 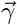 contains at least one causal variant that targets *both* genes *g*_1_ *and g*_2_. Similarly, let 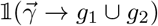 indicate whether the configuration contains a causal variant that targets *either* gene. Then, the edge weight between the two genes is given by:

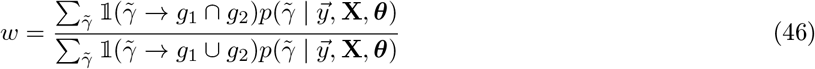

This edge weight is an adaptation of the Jaccard index; it is the probability that both genes are targeted by the *same* variant, divided by the probability that either gene is targeted by *any* variant. Given the nodes and edge weights as just described, we clustered the resulting graph using the Leiden algorithm (Traag et al., 2019) to optimize the Constant Potts Model (CPM) objective. We set the parameter of the CPM to 0.5 – *ϵ*, where *ϵ* is any small number (e.g., 10^−3^). This choice promotes the co-clustering of two genes only if their edge weight satisfies *w* ≥ 0.5. That is, genes tend to co-cluster if at least 50% of the probability mass implicating either gene is shared between them.

#### Merging Gene Clusters across Related Traits

For some of our validation analyses, we combined the genes we identified from multiple GWA studies (e.g., on IBD, CD, and UC). We therefore sought to estimate the number of *distinct* gene clusters implicated by the union of multiple subphenotypes or traits. To this end, we define another weighted, undirected graph, whose nodes are the genes that were assigned to a gene cluster in at least one trait under consideration. Furthermore, let the edge weight between two nodes be the fraction of traits in which the two genes are co-clustered. Then, in order to estimate the number of distinct gene clusters across related traits, we simply cluster this graph using the Leiden algorithm to optimize the CPM, as above. Here, we set the resolution parameter of the CPM to *t*^−1^ – *ϵ*, where *t* is the number of traits under comparison. This condition requires that the average edge density within a cluster exceeds *t*^−1^: each gene must share a gene cluster with every other gene in at least one trait, on average. (Alternative methods of counting the distinct gene clusters yield very similar results.)

#### Expected Number of Causal Variants

Let 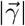 denote the number of causal variants in the model 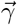, and let 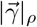 denote the number of causal variants with a GWAS p-value ≤ *ρ*, so that 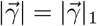. Then, by definition, the expected number of causal variants in the genome, for a given GWAS, is

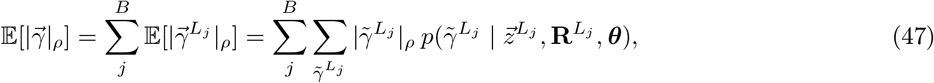

subject to the p-value threshold *ρ*. It is simple to compute this expectation as a matter of course, while calculating the genome-wide variant and gene PPAs, as described above.

### RUNNING THE GENETICS PLATFORM

#### Overview

We developed a proprietary software platform to efficiently implement our computations. The core calculations are primarily written in C++ and distributed with Python code over a dynamic computational architecture on Amazon Web Services (AWS). We designed the platform to process a collection of GWA studies at once, with support for efficiently adding new association studies as they become available. We ran the platform in a number of sequential phases. First, we precomputed the LD partition for each relevant 1KGP super population (or combination of super populations, for multi-ancestry GWAS). We carried out this computation just once, as described above. Second, we standardized the representation of genetic variants, so that the same variant is represented the same way in all GWA studies. At this stage, we imposed quality control (QC) criteria on the variants that we analyzed in subsequent stages. Third, we adjusted the LD partition for each GWAS, to account for the ‘leakage’ of extreme association signals across block boundaries. Fourth, we efficiently computed and stored the Pearson correlation matrix **R** within each adjusted LD block, across all studies. Fifth, we leveraged the results from the previous steps to find and store the most likely causal configurations 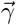 encountered during SSS (typically 150 million per LD block); these approximate the full probability space of all possible causal configurations in each LD block. During this stage, we also computed preliminary variant PPAs, given the uniform prior and an initial estimate of *θ*, in order to find the appropriate nugget regularization of **R**; see below. Sixth, we ran our EM algorithm to find the optimal parameters of the uniform or biochemical prior. Seventh, we evaluated the variant and gene PPAs for each GWAS, using the precomputed sets of causal configurations and prior distributions. Lastly (in the case of the biochemical prior), we grouped the genes with suprathreshold PPA into clusters of genes implicated by shared causal variants, to facilitate the prioritization of candidate causal genes for each trait.

#### Standardization and Quality Control of Genetic Variants

We standardized the representation of genetic variants, which are often reported differently across GWA studies. First, we set the effect allele to the alternative allele in the GRCh37 build of the human genome (switching the sign of the reported z-score, if necessary). We computed the correlation matrix **R** (see below) with respect to this convention for the effect allele. Next, we normalized the position and alleles of each variant with vt `normalize` (Tan et al., 2015). Following the variant normalization, we matched the variants reported in each GWAS to the same variants as reported by the UKBK. We therefore established a common representation of all variants, anchored on the variants in the UKBK. The UKBK polymorphisms define the superset of admissible variants we included in our GWAS analyses. (In part, this is because we computed **R** on UKBK variants.)

Following standardization, we filtered the variants by QC criteria. We considered only biallelic variants that fall within an LD block for a given LD partition. Moreover, we filtered out variants that are very rare (MAF < 0.1%), poorly imputed (INFO score < 0.8), or in Hardy-Weinberg disequilibrium (p-value < 10^−6^), according to the metrics computed by the UKBK. There are more than 12.5 million UKBK variants that satisfy these criteria. Whenever the relevant metrics were computed on a GWAS cohort and made available with the summary statistics, we additionally required variants to pass these QC criteria in the study cohort. For meta-analyses, if the heterogeneity statistic *I*^2^ was provided, we required that variants show minimal evidence of heterogeneity across cohorts (*I*^2^ ≤ 40). For case-control studies, following Neale et al. (2021), we also filtered out variants if the expected number of minor alleles in cases was less than 25.

#### Merging LD Blocks Bridged by Extreme Association Signals

Let *A* denote the association signal of a causal variant whose GWAS p-value is *p*, where *A*:= – log_10_ *p*. Further, denote by *r*^2^ the squared Pearson correlation between a passenger variant and this causal variant. Then, the expected association signal at the passenger variant is approximately *Ar*^2^ (Pritchard and Przeworski, 2001). Now consider the case of a causal variant with an extraordinarily small p-value—say, the smallest number greater than zero that can be represented with a double precision floating-point variable. Such a variant will have an association signal of about 307.65. Even an inert (passenger) variant with a weak correlation (say, *r*^2^ = 0.03) to the causal variant will have a genome-wide significant association signal. When extreme association signals like this occur at the edge of an LD block, a weakly correlated variant in an adjacent block may inherit a strong association. In these circumstances, it appears (artificially) as though an independent association exists in the neighboring block.

While this situation arises very rarely, we nevertheless took care to avoid it. We developed a simple C++ program to customize the LD partition for each GWAS, by merging two LD blocks whenever an association signal from one block would have a material impact on another. Consider a pair of variants in two separate LD blocks on the same chromosome. Let A be the maximum of the association signals at the two variants. We merged the corresponding blocks whenever the correlation between the variants satisfied *r*^2^ > ^3^/_*A*_. In practice, this procedure modifies the LD partition only slightly, and often not at all.

#### Computing the Pearson Correlation Matrix

Due to considerations of data availability, we restricted the application of our platform to GWA studies on cohorts of European ancestry. We therefore computed the Pearson correlation matrix **R** using the individual-level genotype data from the UK Biobank (Bycroft et al., 2018), considering 450,642 unrelated participants who self-identified as ‘white’ and of British ancestry. We wrote an efficient C++ program to compute **R** in each unique LD block across all the GWAS-adjusted partitions, using a single pass through the UKBK VCF files. When new GWA studies become available, an additional pass through the VCF files is required, but only to compute **R** for any unique blocks in the corresponding LD partition.

The white British subset of the UKBK is an excellent reference population on which to compute **R** for European cohorts, owing to its size and the quality of the genotype imputation. Nevertheless, the ancestry of the UKBK is not a perfect match for the European cohorts in GWA studies. Even for association studies performed on the UKBK itself, our matrix **R** is an approximation, since the UKBK subset we used to compute the correlations will differ somewhat from the samples included in the association study. It is therefore necessary to regularize the correlation matrix **R**, by adding a small, positive constant *η* to the diagonal of the matrix. This nugget regularization is well known and has been proposed previously for statistical fine-mapping (Chen et al., 2015). For association studies in the UKBK, we set *η* = 0.02. For studies in other European cohorts, we set *η* to the minimum value (typically in the range 0.1–0.2) such that the fine-mapping passed our quality control tests (see below).

#### Shotgun Stochastic Search

The evaluation of gene and variant PPAs by equations (29) and (30) requires a summation over all possible configurations of causal variants in each LD block. Attempted directly, that summation is intractable. Instead, we approximate the sum with Shotgun Stochastic Search (SSS), using our own implementation of the method described in Hans et al. (2007). Briefly, we perform a stochastic search to identify and store the most likely causal configurations in each LD block, until the variant PPAs computed from those configurations converge. The stochastic search is computationally expensive, so we store the approximating collection of causal configurations in a binary file for each LD block. The saved configurations are reused whenever we evaluate an equation that requires the summation over the causal models in a block.

#### Nugget Regularization and Quality Control of Fine-mapping Results

Accurate statistical fine-mapping of GWAS data relies critically on: 1.) the quality of the genotype imputation in the study; and 2.) a close correspondence between the correlation matrix **R** in the study and reference (UKBK) populations. These criteria may not hold for any particular GWAS. We therefore took QC measures to exclude studies that could not be fine-mapped. We evaluated two criteria on each GWAS, choosing a small value for the nugget regularization, *η* ≤ 0.2, at which the study passed both conditions. First, we investigated the GWAS p-values of all variants with a suprathreshold PPA (e.g., greater than 50%). A symptom of spurious fine-mapping is the apparent discovery of many likely causal variants without a strong trait association (e.g., GWAS p-values greater than 10^−2^). We required that variants with high PPAs should have very low GWAS p-values. Second, we assessed the distribution of likely causal variants in biochemically active DNA. We required that variants with progressively higher PPAs should be proportionately more likely to target DNA with a predicted biochemical function. If a study did not pass either of these QC criteria with a nugget less than 0.2, we excluded it from our GWAS compendium. We found that older GWA studies, published before 2017, tended to fail both criteria.

#### The EM Algorithm and Final Computation of PPAs

An assumption of our EM algorithm for learning the biochemical prior is that the GWA studies are analyses of distinct traits. We therefore selected a collection of studies on which to learn the prior, balancing the size of the catalog with its diversity. Running our EM algorithm on the studies enumerated in Table S7 yields a ratio of Bayes factors for each variant annotation, which we assume is constant across traits. Recall equation (35). Given the ratio of Bayes factors for each variant annotation, and the odds that an unannotated variant is causal (which we learn separately for each trait), we evaluate equation (38) to obtain the prior probability for each variant in any study. In practice, this procedure entails running the EM algorithm once on the GWAS compendium, and then storing the resulting ratio of Bayes factors for each variant annotation. At this stage, we have the biochemical prior for all traits in the compendium. In order to obtain the prior for any GWAS not in the collection, we run the same EM algorithm on the GWAS, except that we hold the precomputed Bayes ratios constant and learn only the prior odds for unannotated variants. We then apply equation (38) to obtain the prior for the new trait. Running the EM algorithm for the uniform prior is even simpler: the single parameter is optimized independently for each trait, without the need for a GWAS compendium. Given the prior parameters for any trait, it is then a matter of software engineering to efficiently compute the variant and gene PPAs, with equations (29) and (30).

### COMPARISON OF DISEASE GENE DISCOVERY ACROSS METHODS

We evaluated the recall of well-known disease genes in IBD and obesity, by our platform and by preexisting methods. We compared our method to the Activity-by-Contact map (ABC; Nasser et al., 2021), Summary-based Mendelian Randomization (SMR; Zhu et al., 2016), Summary MultiXcan (Barbeira et al., 2018), and Coloc (Giambartolomei et al., 2014). When evaluating each method on IBD, we took the union of genes discovered from three GWA studies (de Lange et al., 2017): on Crohn’s disease (CD), Ulcerative colitis (UC), and IBD (in which the CD and UC cases were combined). In the case of ABC, the data were obtained from an even larger study of IBD (Nasser et al., 2021). Likewise, for obesity, we combined the genes discovered from GWA studies of BMI (Canela-Xandri et al., 2018; Neale et al., 2021), WHRadjBMI (Pulit et al., 2019), and BFP (Neale et al., 2021). We computed recall of known IBD genes from two validation sets. First, a set of 83 genes from a recent review (Graham and Xavier, 2020); and second, a subset of these defined by Nasser et al. (2021), including only genes within 1 Mbp of a GWS association signal in non-coding DNA. For obesity, we obtained a set of 27 genes discovered from protein-altering variants associated with BMI (Turcot et al., 2018; Akbari et al., 2021). Since the ABC predictions were generated recently (Nasser et al., 2021), with up-to-date genetics resources, we obtained those results directly from Nasser et al. (2021), without re-running any analysis software. On the other hand, for the older methods SMR, MultiXcan, and Coloc, it was necessary to re-run the software on the latest genetics data.

#### SMR

We downloaded and ran SMR (https://yanglab.westlake.edu.cn/software/smr/#Overview) using the default settings recommended by the authors. We used summary statistics from (de Lange et al., 2017; Canela-Xandri et al., 2018; Pulit et al., 2019; Neale et al., 2021) and the white British subset of the UK Biobank (Bycroft et al., 2018) as a reference population (i.e., the same summary statistics and reference population as in our work). We used eQTL data from GTEx Version 8 (GTEx Consortium, 2020), using all available tissues as processed and made publicly available by the SMR authors. We used the Storey-Tibshirani (Storey and Tibshirani, 2003) method to adjust for multiple comparisons, and considered genes significant with *q* < 0.05 and *p*_HEIDI_ > 0.05; the latter is the mechanism to reject gene-trait associations not directly induced by causal variants.

#### Summary MultiXcan

We downloaded the MetaXcan software (https://github.com/hakyimlab/MetaXcan) and, starting with the summary statistics in (de Lange et al., 2017; Canela-Xandri et al., 2018; Pulit et al., 2019; Neale et al., 2021), followed the GWAS harmonization and imputation tutorial recommended by the authors. We downloaded the MASHR model weight and covariance files for GTEx Version 8 eQTLs from https://predictdb.org/, for every available tissue type. Using the MetaXcan repository, we ran S-PrediXcan on the GWAS and the MASHR models of each tissue, and subsequently ran S-MultiXcan on the collection of tissues. For each GWAS, we used the Storey-Tibshirani method to compute q-values, and considered genes significant if *q* < 0.05 and *p_b_* < 10^−5^, where *p_b_* is the most significant p-value for the gene in any tissue. The authors recommend the latter criterion to remove spurious associations.

#### Coloc

We downloaded and installed the `coloc` R package, together with the summary statistics from GTEx Version 7 eQTL studies in all available tissues. (GTEx Version 7 is the latest release for which the summary statistics are freely available.) For each gene in our validation data sets (from IBD or BMI), we checked if there was a GWS association within 1Mbp of the gene and a SNP in the eQTL study with *p* < 10^−5^. If so, we ran `coloc` (approximate Bayes factor method) on the lead SNP and up to 1000 surrounding SNPs. We included the gene as significant if the posterior probability of colocalization exceeded 0.5. Unlike the other methods, Coloc is not intended to be run genome-wide, and was only run on the genes from the validation set.

## DATA AND CODE AVAILABILITY

All of the data used in this research are publicly available and are accessible as described in this manuscript. We present the full mathematical and statistical theory constituting the methods of this research. Computer code to implement the mathematics may be written in a variety of ways and run on different computational architectures. Additional information required to apply the platform introduced here is available from the corresponding author upon request.

### SUPPLEMENTAL INFORMATION

#### Supplemental figures

**Figure S1.**
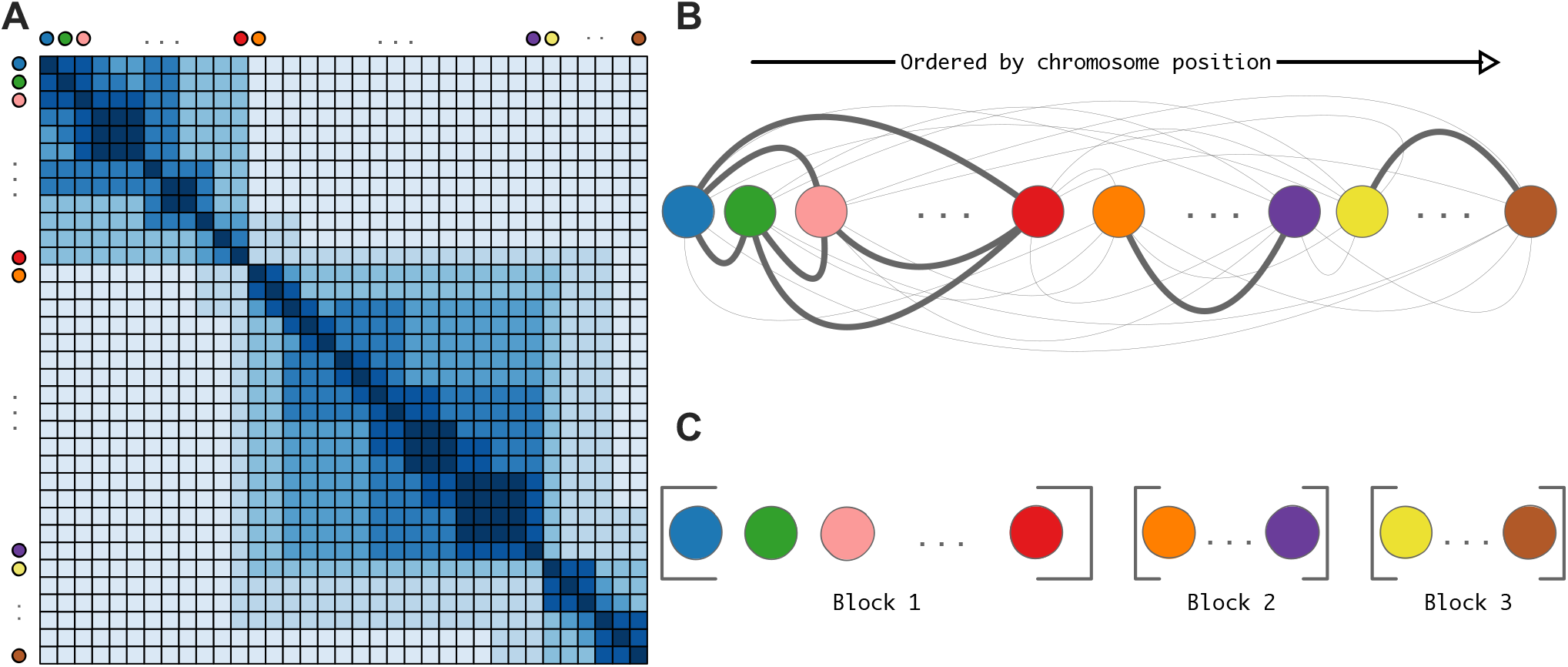
LD block discovery is a graph clustering problem, related to Figure 1. Genetic variants, illustrated by colored circles, are partitioned into LD blocks that achieve the global optimum of a graph clustering problem. (A) The input to the LD partition algorithm is a VCF file containing individual-level genotype data, from which we compute correlations between variants. We depict the resulting correlation matrix as a heatmap, with darker blue indicating stronger correlation (*r*^2^). Each element of the correlation matrix describes the sample correlation between the alleles of a pair of variants. The ordering of variants along the rows and columns of the matrix follows their linear arrangement on the chromosome. (B) The correlation matrix is interpreted as the weighted adjacency matrix of a graph that we embed along a line, preserving the chromosomal ordering of variants. Thicker edges denote stronger correlations between endpoints. (C) The algorithm outputs a partition of the graph that maximizes a clustering score exactly, subject to the ordering constraint. We interpret each partition component as an LD block.

**Figure S2.**
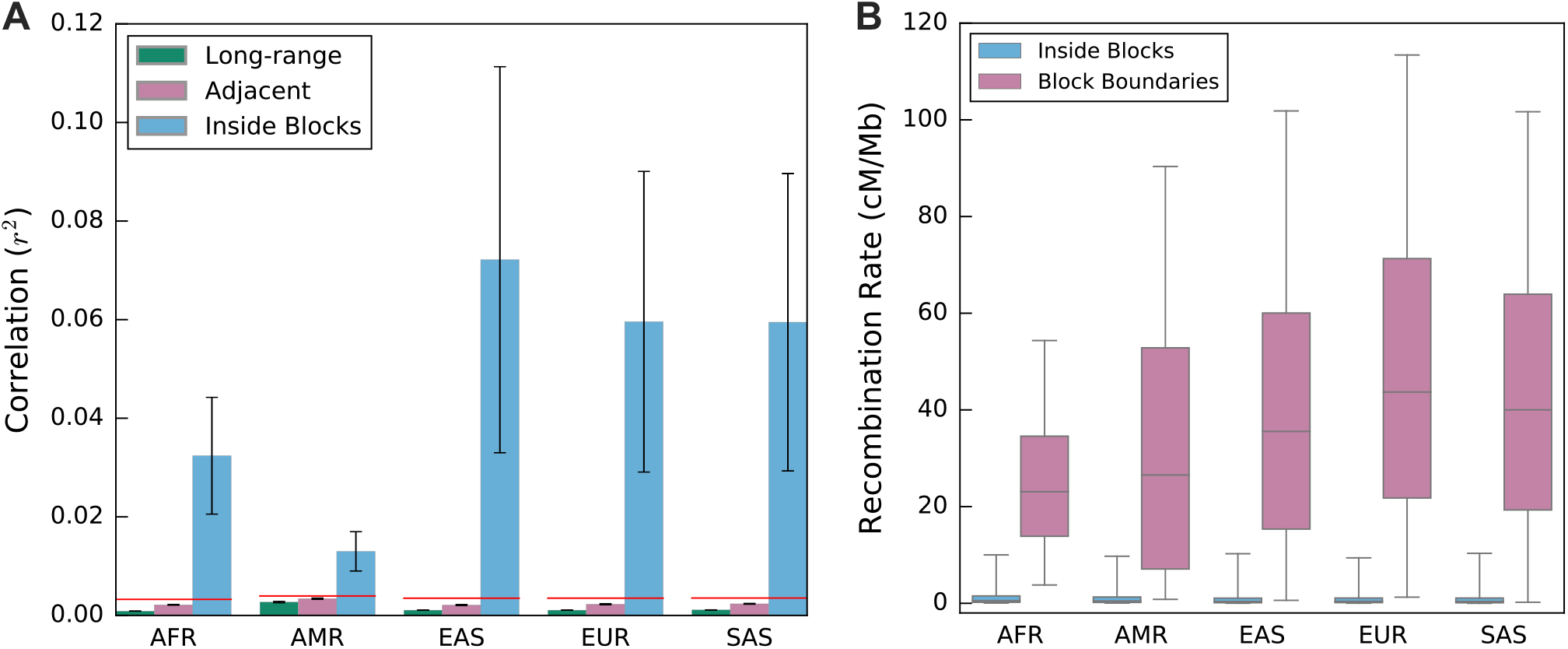
LD blocks exhibit weak inter-block correlations and high recombination rates at boundaries, related to Figure 1. (A) For each super population, every input correlation is assigned to one of three categories: inside an LD block, between two neighboring blocks, or separated by at least one other block (long-range). We plot the mean (± standard deviation) of the correlations in each resulting category. The red lines denote the minimal sample correlation we consider to be non-negligible, for each population. Note the low variability in correlations between adjacent and distant LD blocks (the standard deviations are barely visible). (B) Box plot of the super-population recombination rates, both within blocks (sky blue) and at the junctions between blocks (reddish-purple). For the within-block distribution, we randomly sample 10% of observations for display (for computational convenience). Each box delineates the interquartile range (middle 50% of the recombination rate distribution), with the median denoted by the horizontal line. The whiskers extend to the 5th and 95th percentiles. Observe that the recombination rates are considerably higher at LD block boundaries than within blocks (boxes are barely visible).

**Figure S3.**
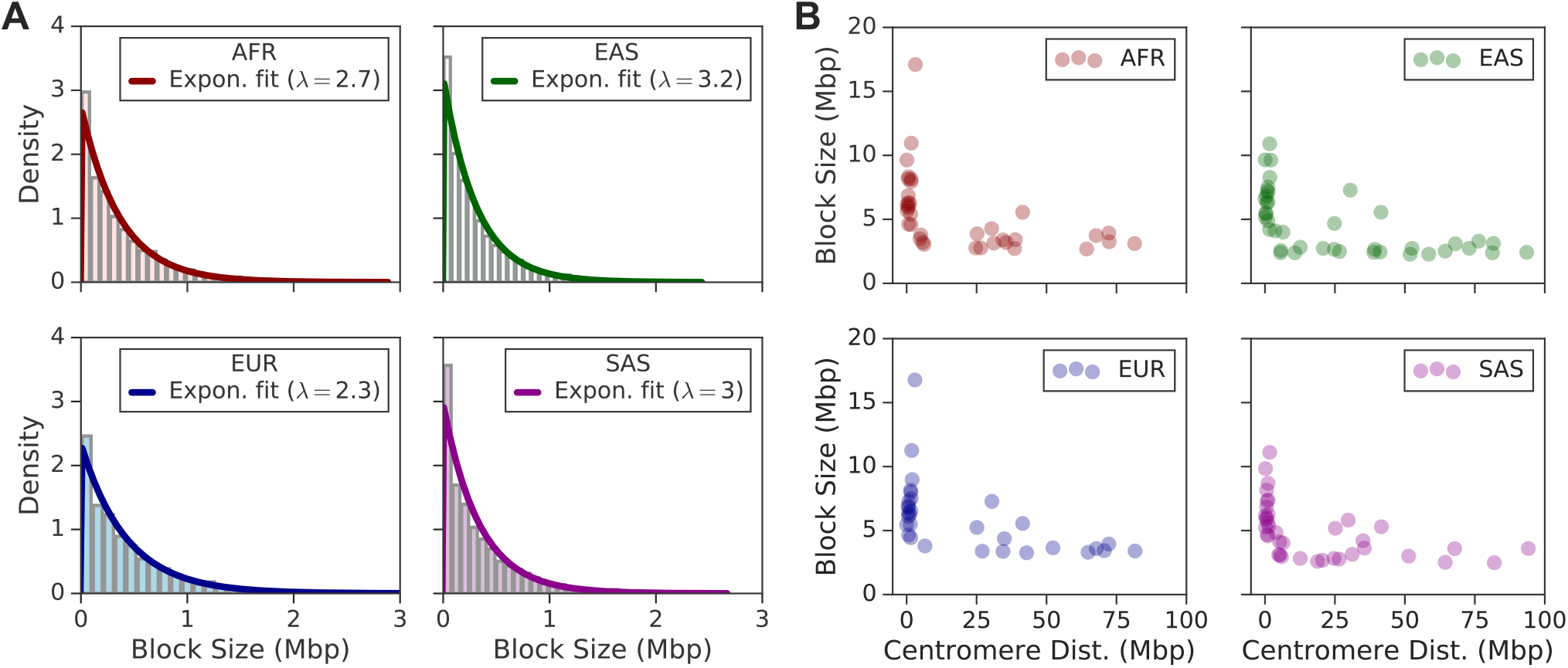
LD block sizes follow the exponential distribution, with the longest blocks near centromeres, related to Figure 1. (A) The histogram of block sizes for each population, where sizes in the top 0.5% have been removed. The thick line in each subplot shows the fit of an exponential distribution, with the given rate parameter λ, to the underlying histogram of block sizes. The scale of the y-axis reflects the fitted exponential probability density function, not the counts of the underlying histogram. (B) Scatter plot of the block sizes trimmed from the left subfigure as a function of distance to the centromere (measured from the block midpoint). Block sizes in the top 0.5% tend to occur near the centromere, where recombination is repressed.

**Figure S4.**
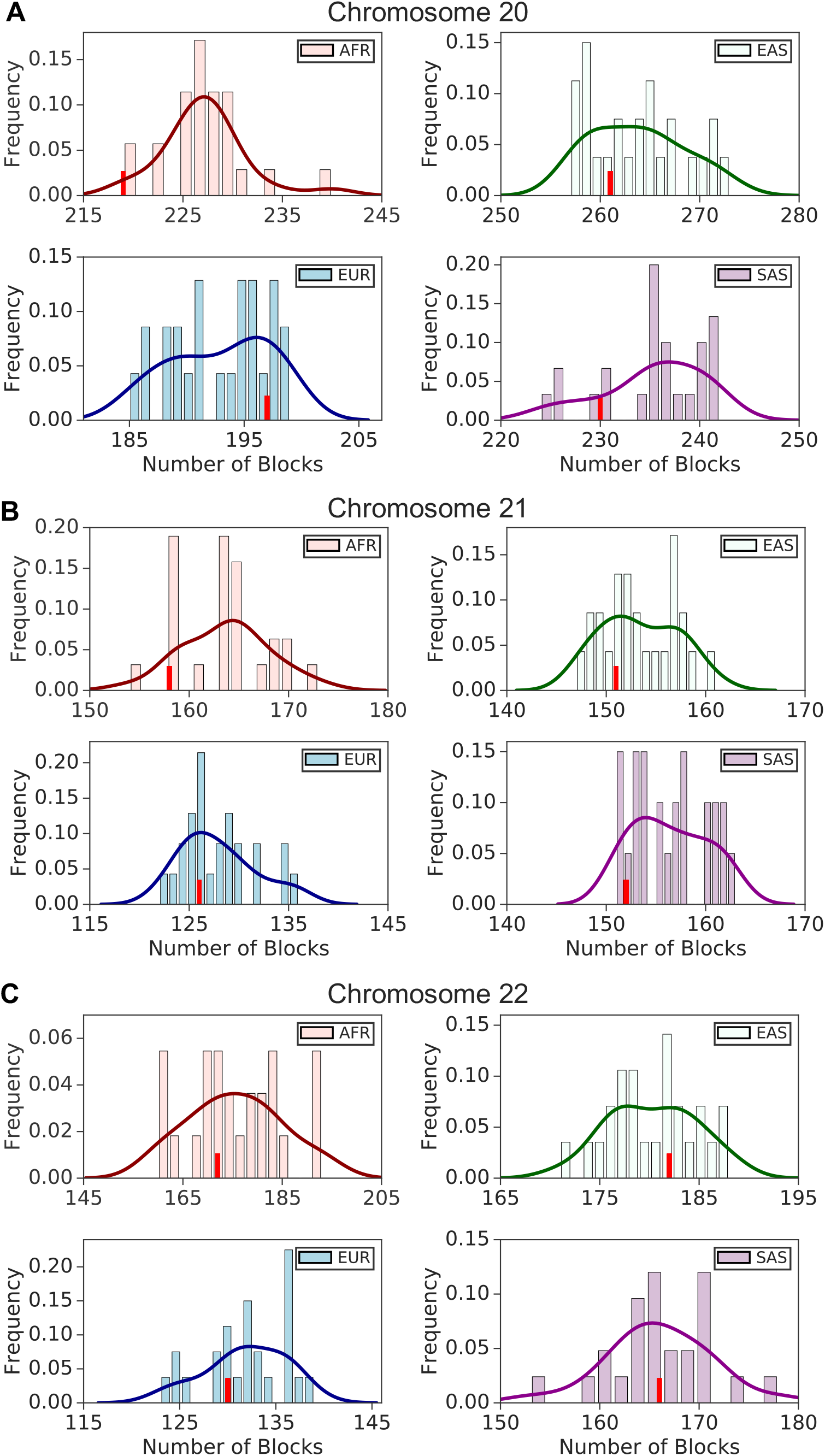
The LD partitions are independent of sample size, related to Figure 1. The LD partitions are largely invariant to the size of the reference sample from which they are inferred. The sample size of the AMR super population is substantially smaller than the unadmixed super populations. To validate that the larger blocks in AMR, relative to the other populations, are not an artifact of the small AMR sample, we randomly downsampled the other super population cohorts. The LD partitions from the randomly downsampled cohorts contain a similar number of blocks compared to the full data. (A-C) Distribution of block counts across 25 random subsamples of each unadmixed super population, on chromosomes 20–22. The full sample of each population (*N_AFR_* = 661, *N_EAS_* = 504, = 503, *N_SAS_* = 489) was randomly subsampled, without replacement, to the size of the AMR super population (*N_AMR_* = 347), and the LD partitions were computed on the random subsamples. The red tick mark in each histogram denotes the number of blocks from the full population. Despite reducing the sample size, the number of LD blocks in the AFR, EAS, EUR, and SAS super populations remains stable. In every population, the number of LD blocks computed with the full data sample falls within the narrow distribution obtained from the random subsamples. Furthermore, the number of blocks remains significantly greater than the count observed in AMR at this sample size (45, 33, and 19 for chromosomes 20–22, respectively).

**Figure S5.**
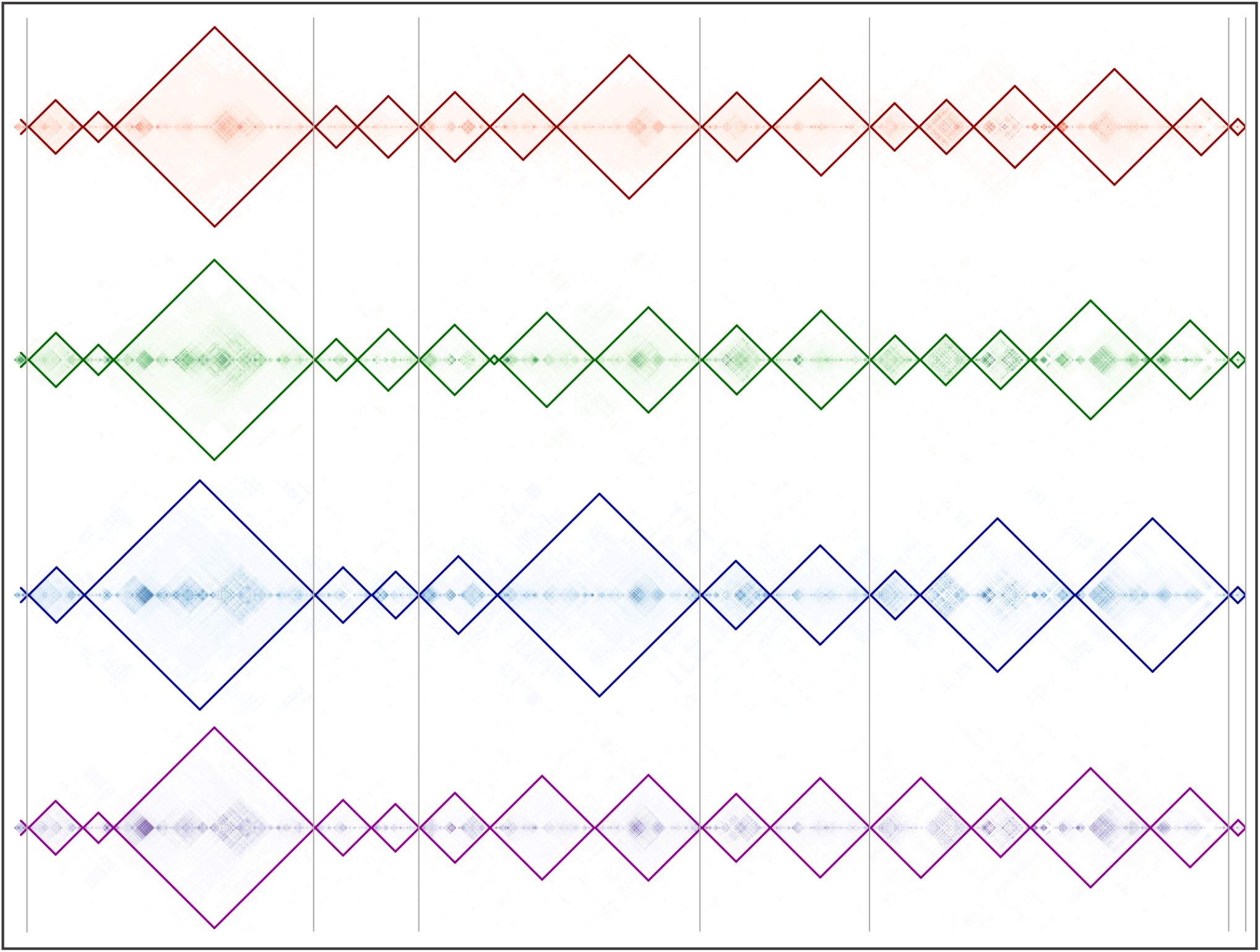
Shared LD boundaries across super populations, related to Figure 1. An example of ‘universal’ LD boundaries in the *APOE* locus, spanning the region on chromosome 19 from 43.9–48.6 Mbp (GRCh37 coordinates). The correlation matrices in each unadmixed 1KGP super population are overlaid with their corresponding LD block outlines. The blocks in the AFR, EAS, EUR, and SAS super populations are shown, from top to bottom, in red, green, blue, and purple, respectively. The thin vertical gray lines mark the positions of LD boundaries that are shared across all of these populations, at a resolution of 2 kbp; they denote ‘universal’ LD block boundaries in the consensus partition.

**Figure S6.**
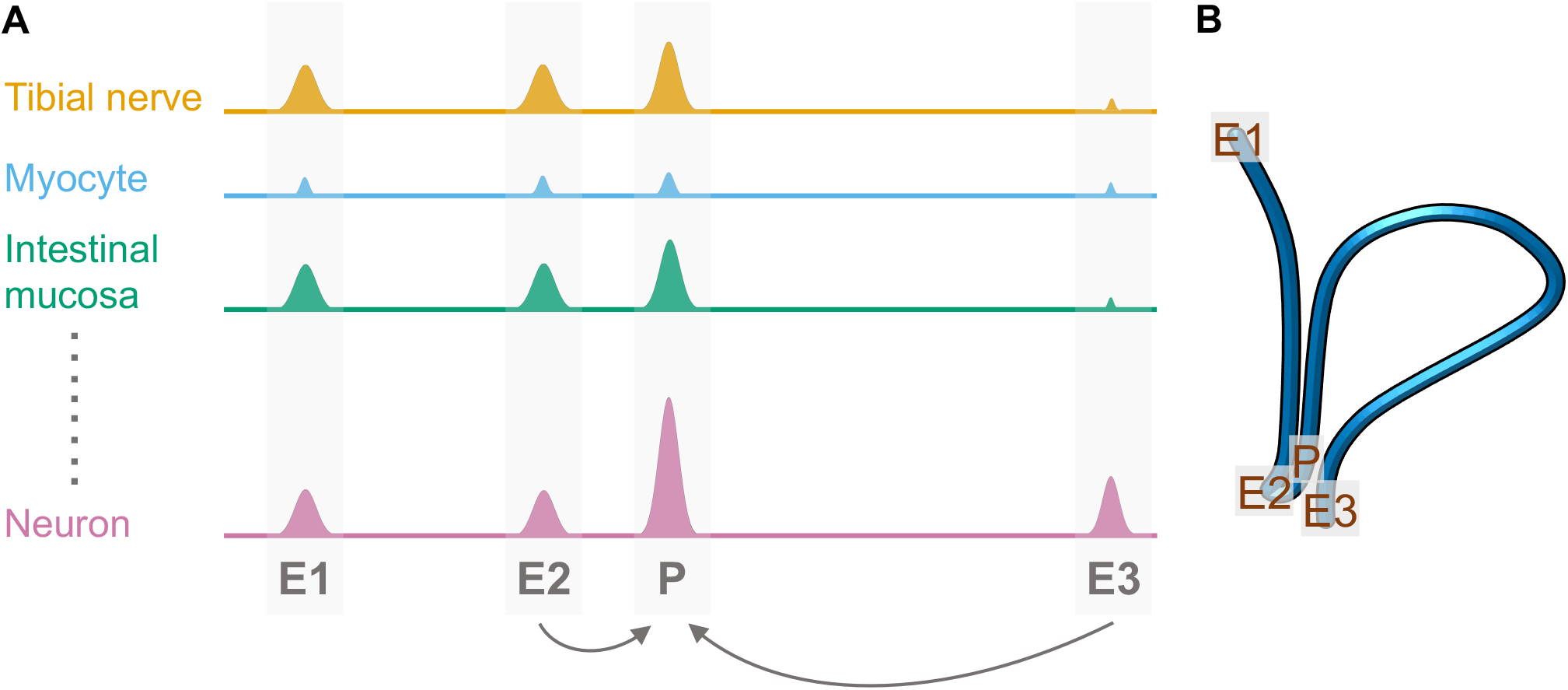
Discovery of enhancer-promoter interactions, related to Figure 2. Schematic of our enhancer-promoter modeling. For each promoter P in the genome, we built an elastic net regression model to predict the chromatin accessibility at P as a function of the accessibility at *cis*-regulatory elements (CREs), E1-E3, within 2Mbp. CREs that predict the open chromatin signal at P across cell types, subject to constraints on sparsity and physical proximity, are inferred to be enhancers of P. (A) Idealized open chromatin profiles across a collection of cell types from ENCODE, showing DNase-seq (DHS) signals at regulatory elements (shaded regions). CREs E2 and E3 are predicted enhancers for P. Increases in the DHS signal at E2 and E3 are independently predictive of increases in promoter accessibility, and Hi-C data indicates they physically interact with P (see panel B). On the other hand, E1 is rejected as an enhancer for P, because the collinear enhancer E2 accounts for its covariation with P (sparsity constraint) and E1 does not physically interact with P (proximity constraint). (B) Hypothetical chromatin loop that brings P into close physical proximity with E2 and E3, but not E1. Actual contact frequencies between regulatory elements are obtained from Hi-C data.

**Figure S7.**
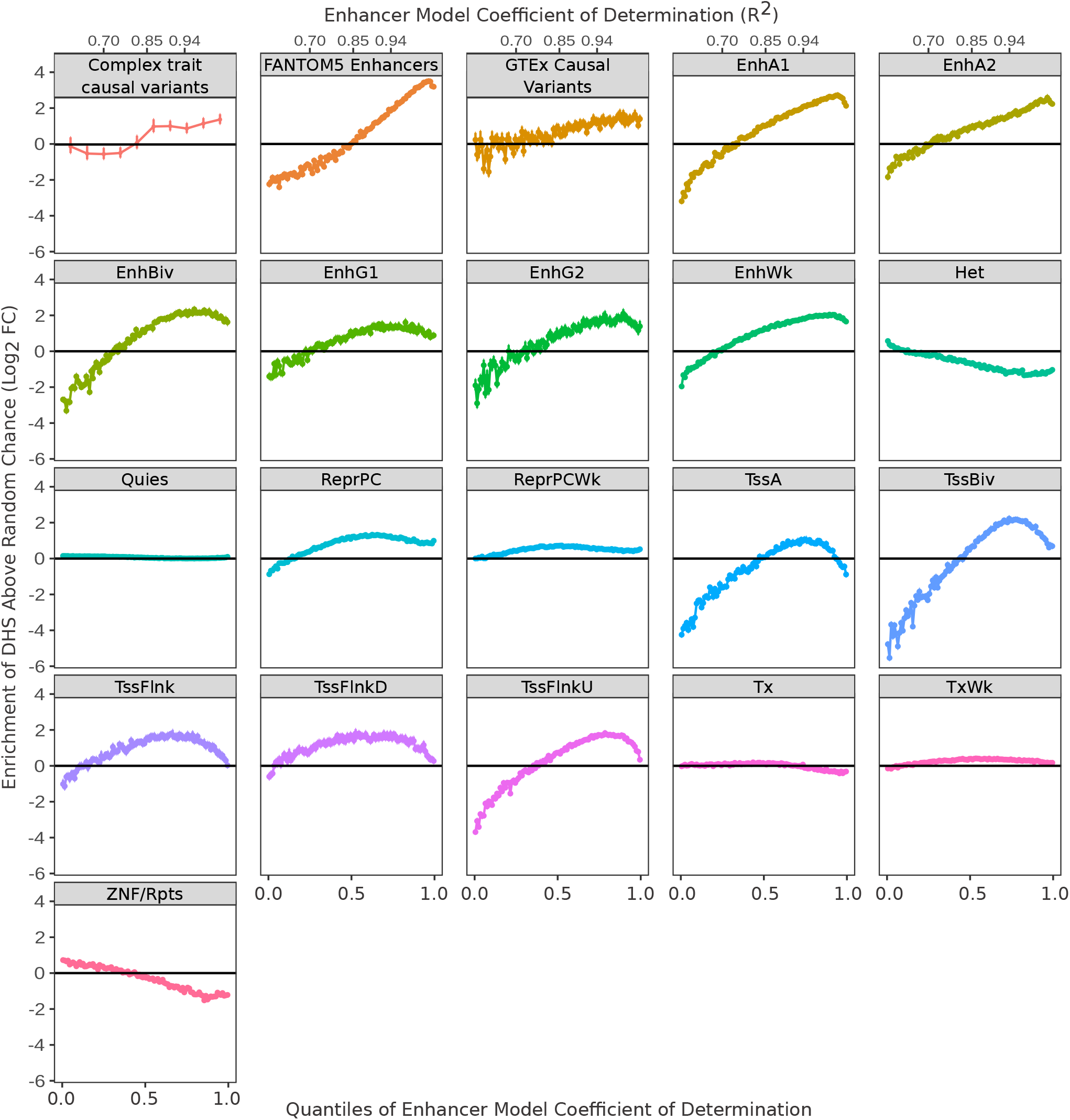
Identification of putative enhancers, related to Figure 2. Enrichment of putative enhancers, defined by various functional genomic features, in DHSs. We calculated the coefficient of determination (*R*^2^) for each DHS model, and divided the DHSs into equal sized bins ordered by *R*^2^. To compute enrichment, we evaluated the fraction of DHSs in each *R*^2^ bin that overlap with a given genomic feature, compared to the overlap after randomly reshuffling the positions of the features (null hypothesis). We plot the enrichment of DHSs in each genomic feature, relative to chance, as a function of *R*^2^. We considered the following genomic features, which are indicative of enhancers: causal variants from our fine-mapping of GWAS data (PPA > 0.5 using the uniform prior, restricted to non-exonic and non-splice site variants); eRNAs from FANTOM5 (Andersson et al., 2014); causal variants from GTEx eQTLs (PPA > 0.9, restricted to non-exonic and non-splice site variants); and distinct chromatin states defined by a genomic segmentation algorithm (Ernst and Kellis, 2012, 2017; Boix et al., 2021). Points denote the mean enrichment and error bars indicate the standard deviation over *N* samples from the null, where *N* is 20 for chromatin states and 50 for all other features.

**Figure S8.**
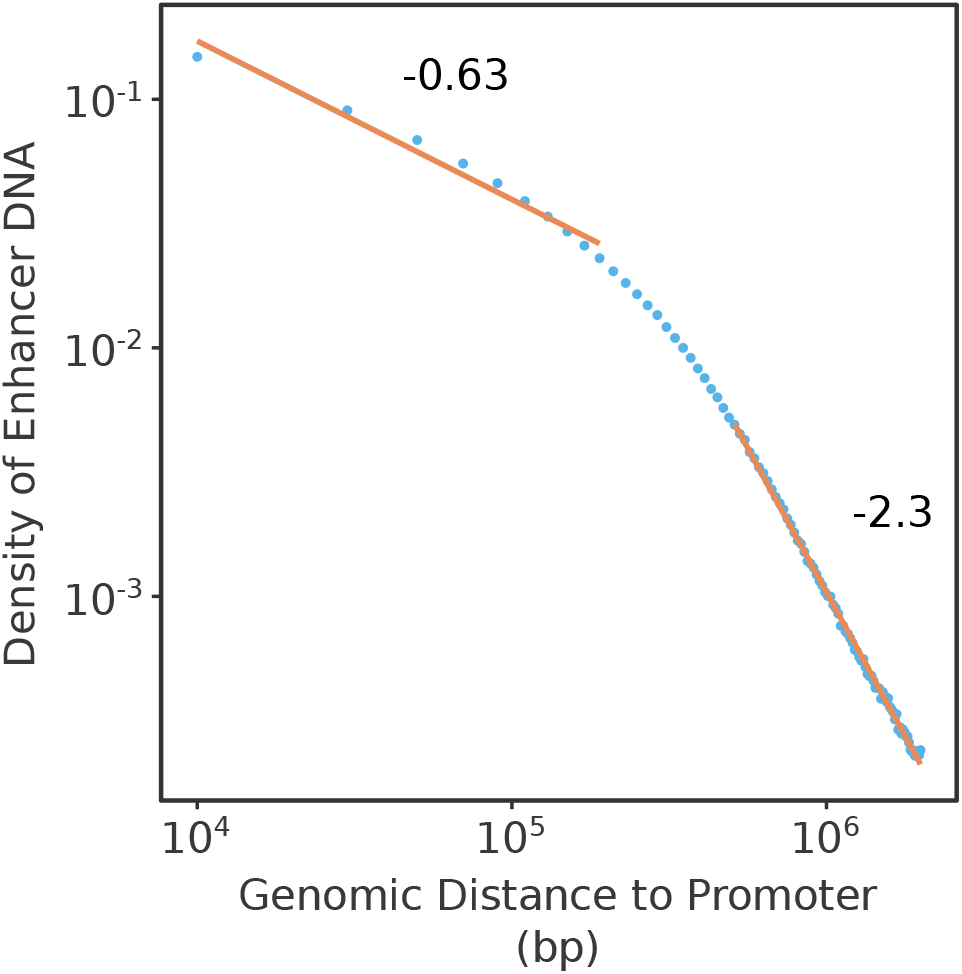
Density of enhancers decays with increasing distance, related to Figure 2. Fraction of DNA predicted to regulate a gene as a function of distance from the promoter. We considered all the enhancer-promoter interactions in our map, stratified into spatial bins based on the distance between each enhancer and promoter. Each bin spans 40 kbp of DNA (corresponding to two 20 kbp windows equidistant from the promoter, upstream and downstream). Denote by *N_b_* the number of enhancers in some bin *b* (i.e., the number of enhancers within the corresponding distance from their linked gene). Further, let *w* be the average enhancer width (in bp), *f* the average fraction of enhancer DNA that is unique (i.e., not double-counted by overlaps with other enhancers), and *G* the number of genes represented in the map. Now, consider an average gene. The density, *D_b_*, we compute for each bin is the fraction of DNA, at the distance *b* from the gene, that encodes an enhancer for the gene:

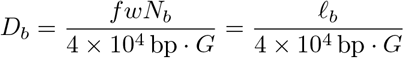

where *ℓ_b_* is the total length of enhancer DNA operating at the distance *b*.

**Figure S9.**
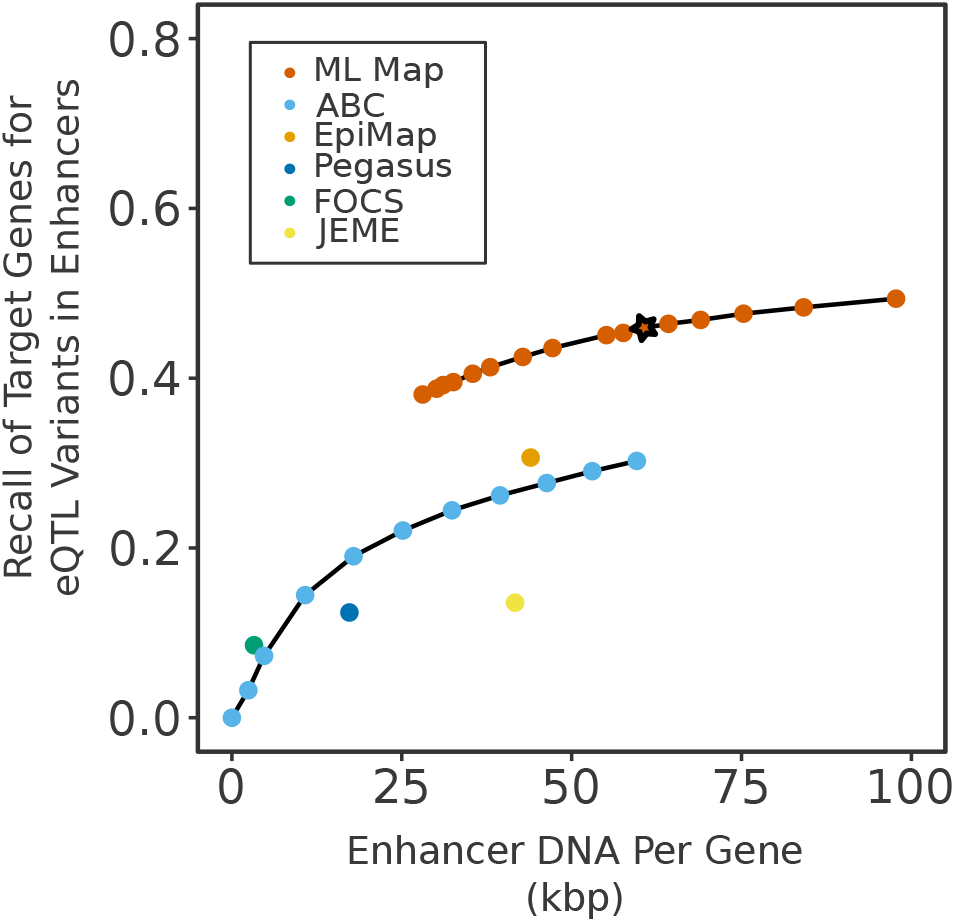
Validation rate of eQTL interactions, related to Figure 2. The validation rate of our enhancer-to-gene map and alternatives against a set of fine-mapped GTEx eQTLs, including all variants *except* those that are in: (1) a DHS within 500 bp of the transcription start site of the target gene; (2) the mature transcript (including coding region and UTRs) of the target gene; or (3) a known splice site for the gene. Therefore, it includes all fine-mapped variants that may function via distal regulatory interactions. The star denotes the density selected to generate the variant-to-gene map.

**Figure S10.**
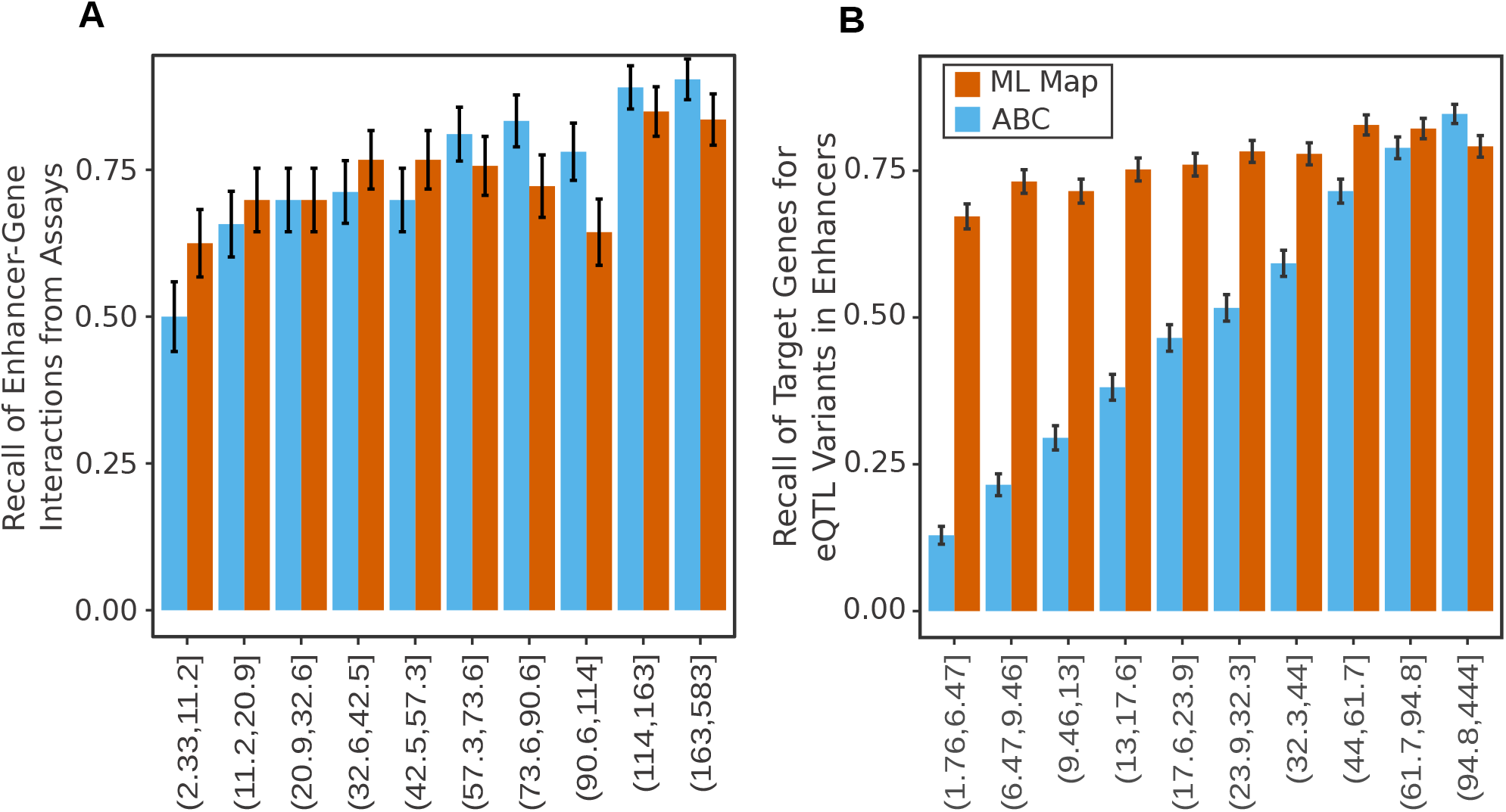
Validation rate versus regulatory element activity, related to Figure 2. The validation rate of the ML map and ABC, stratified by the maximum accessibility (DHS signal) of the validated enhancer. Each bin contains an equal number of enhancers. The max. accessibility (measured in RPKM) was recorded over the collection of cell types in ENCODE (see the *Methods*). (A) Comparative validation rate on enhancers discovered from functional genomics assays (e.g., CRISPR inhibition studies). (B) Validation rate on enhancers uncovered by GTEx. Observe, first, that the enhancers identified by functional genomics assays tend to have greater accessibility than those discovered from GTEx (at least in ENCODE cells). Second, the performance of ABC on the GTEx validations is very sensitive to the magnitude of the enhancer activity, whereas the ML map performs well across the full range of enhancer activities.

**Figure S11.**
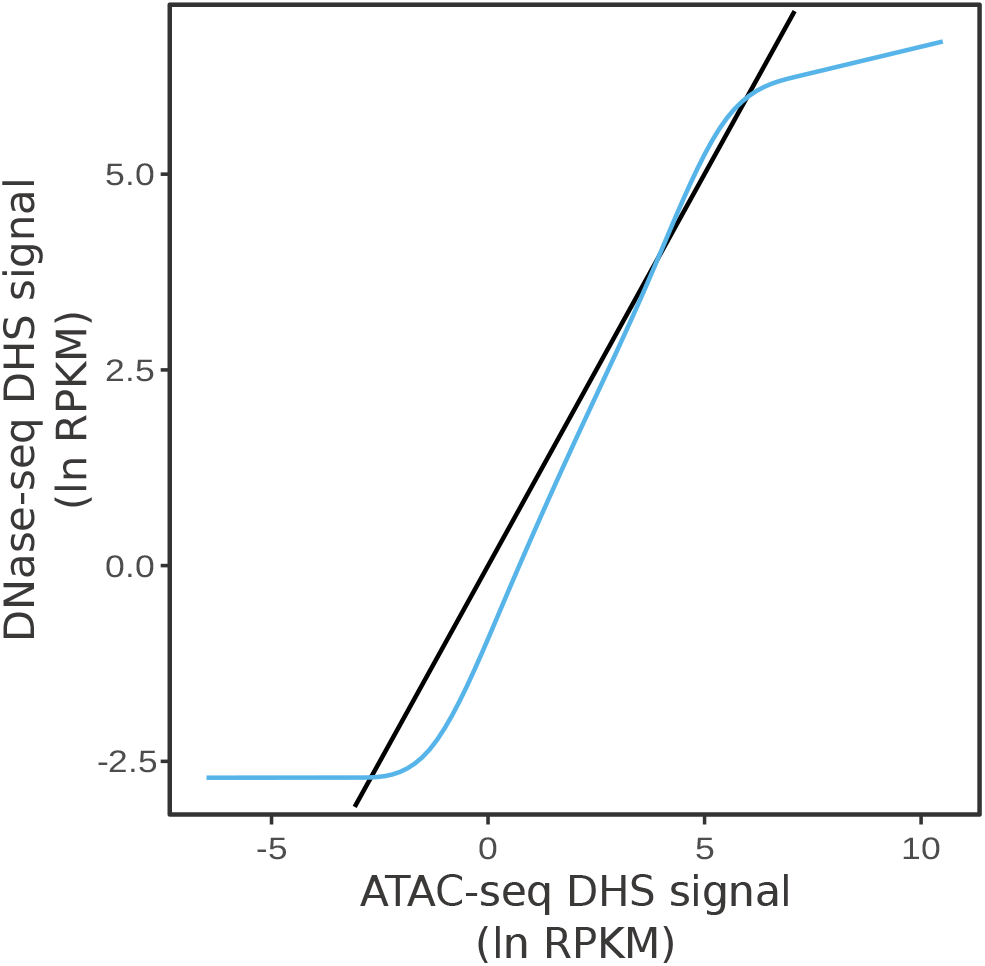
ATAC-seq to DNase-seq signal conversion, related to Figure 2. To include chromatin accessibility data from as many cell types as possible, we used both ATAC-seq and DNase-seq data in the enhancer-to-gene models. We therefore fit a monotonic smoothing spline to transform ATAC-seq signals to the expected DNase-seq data. We chose ENCODE data from the adrenal gland of the same donor to fit the spline, due to the high sequencing depth from both assays in those data. We note that over a large range (roughly −2.5 to 5 ln RPKM), DNase- and ATAC-seq measurements yield very similar signals on average; ATAC-seq is more sensitive outside this range. Black line, the identity *y* = *x*; blue curve, the transformation from ATAC-seq to DNase-seq.

**Figure S12.**
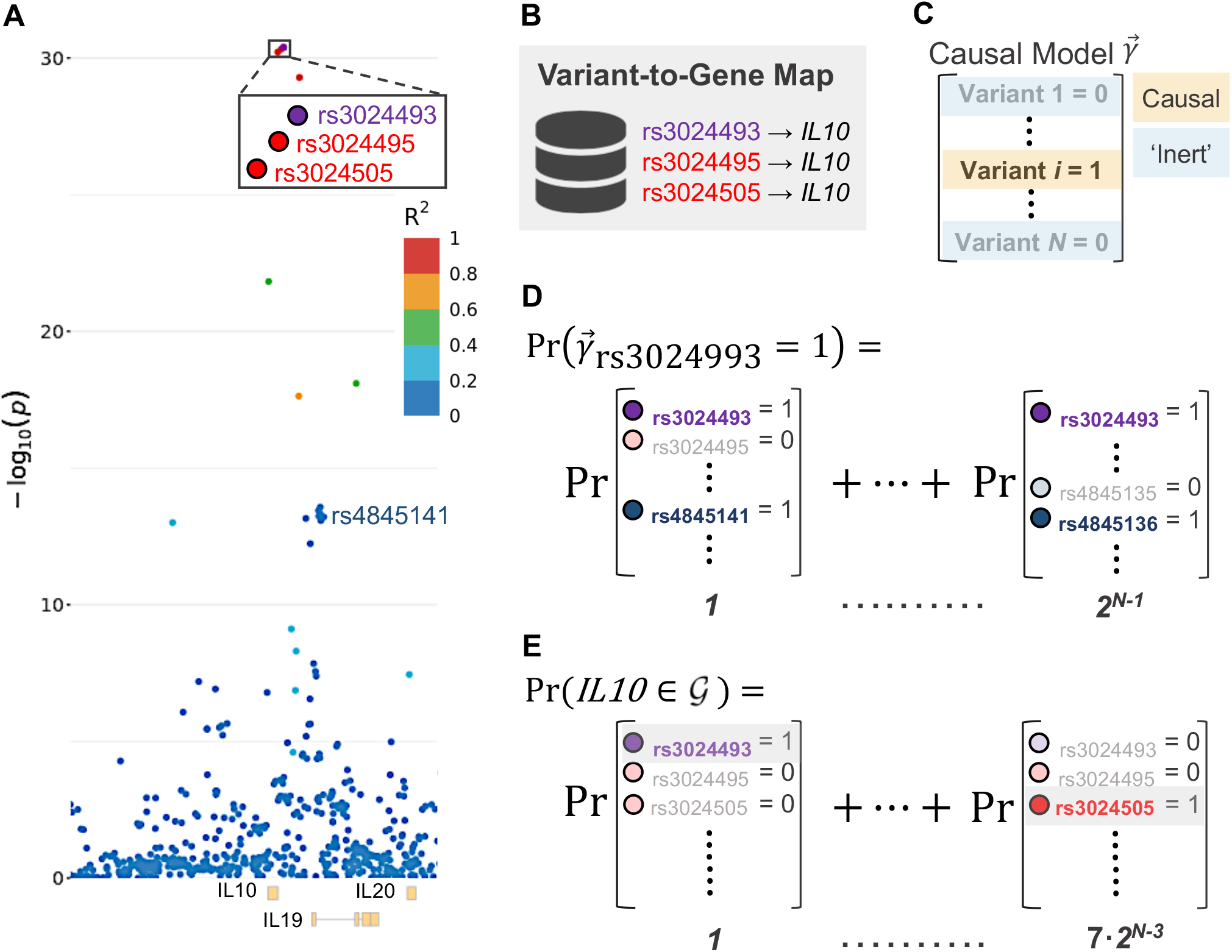
Statistical fine-mapping schematic, related to Figures 3–5. A simplified depiction of statistical fine-mapping, to illustrate how conventional approaches evaluate the posterior probability of association (PPA) for variants, and how we extend this methodology to compute an analogous quantity for *genes*. (A) The association signals for each variant in a genomic locus, from a GWAS of IBD (de Lange et al., 2017). The color of each point (variant) indicates the strength of its Pearson correlation to the lead SNP in the locus, rs3024493 (purple). (B) The three variants with the strongest association signal are all linked to the cytokine gene *IL10* by the variant-to-gene map. (C) A causal model 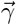, or causal configuration, is an indicator vector that specifies which of the *N* variants in the genome mediate IBD risk. The true (unknown) value of 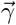 is a complete enumeration of the GWAS variants underlying a phenotypic trait. (D) Theoretically, to compute the probability (PPA) that any variant is causal (here, rs3024993), we first compute the probability of all 2^*N*^ possible causal models that could describe the genetic architecture of the disease. Then, we sum the probability of all models that include the variant (rs3024993) as causal; there are 2^*N*–1^ such models. In practice, implementing that calculation naively (directly) is intractable, so we exploit mathematical features of the problem that dramatically reduce the computational complexity. (E) 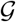 denotes the set of genes that are targeted by at least one causal variant. To compute the probability that any gene (say, *IL10*) is targeted by a causal variant (the gene’s PPA), we sum the probability of all configurations containing at least one causal variant that targets the gene. For instance, if *IL10* is targeted only by the variants rs3024493, rs3024495, and rs3024505, then its PPA could be obtained by summing the probabilities of all models in which at least one of these variants is causal. There are 7 · 2^*N*–3^ such models. As in panel D, we employ a computationally efficient strategy to calculate gene PPAs in practice. Formally, variant and gene PPAs are different marginals taken over the same underlying probability distribution, defined by the space of all 2^*N*^ possible genetic architectures of a trait.

**Figure S13.**
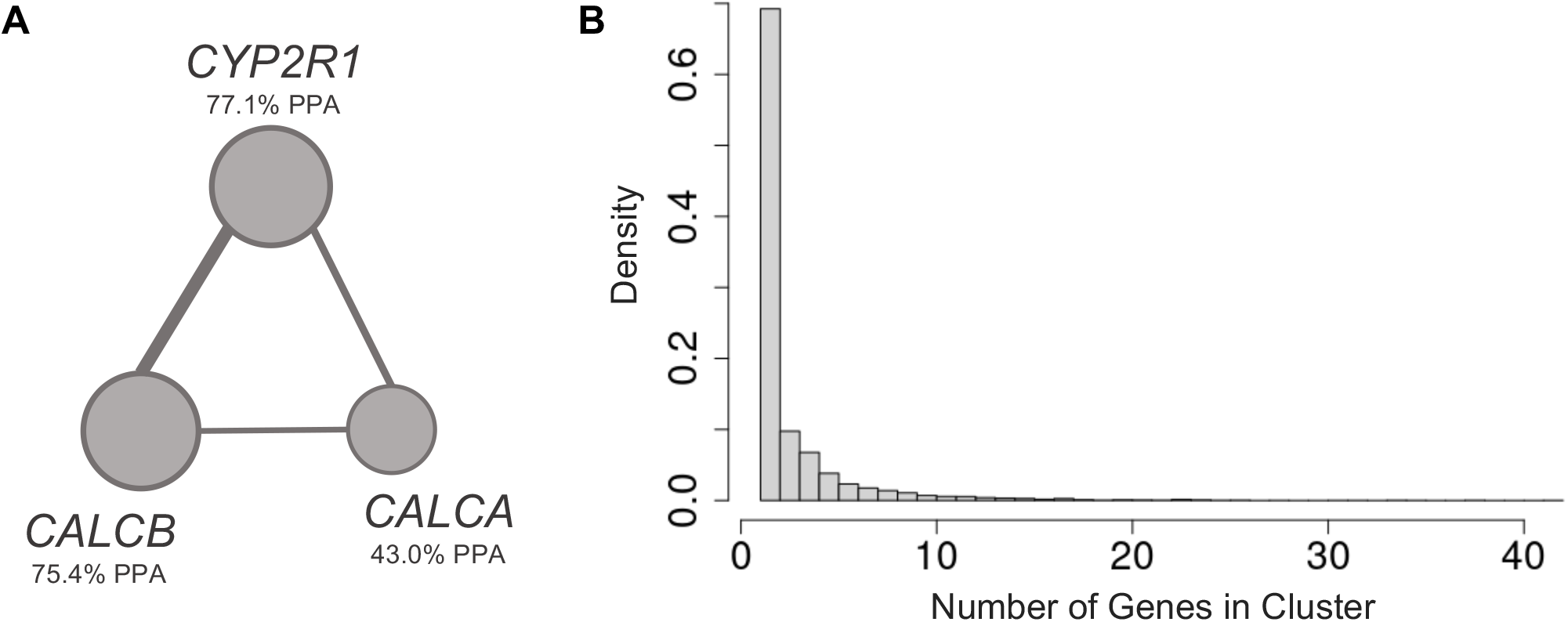
Gene clusters, related to Figure 5. (A) An example gene cluster, from our analysis of self-reported migraine in the UK Biobank (Neale et al., 2021). *CALCA* and *CALCB* encode the *α* and *β* isoforms of CGRP, respectively. Together with its receptor, CGRP is the target of an entire class of migraine therapeutics, while *CYP2R1* is a monooxygenase involved in the metabolic activation of vitamin D. Based on these data, one might prioritize *CALCA* and/or *CALCB* as the likely causal gene(s) in this cluster. (B) The distribution of the number of genes in each gene cluster, taken over all gene clusters computed in all traits we analyzed. The width of each histogram bin is 1, so that the height of the n-th bin indicates the fraction of gene clusters containing *n* genes.

**Figure S14.**
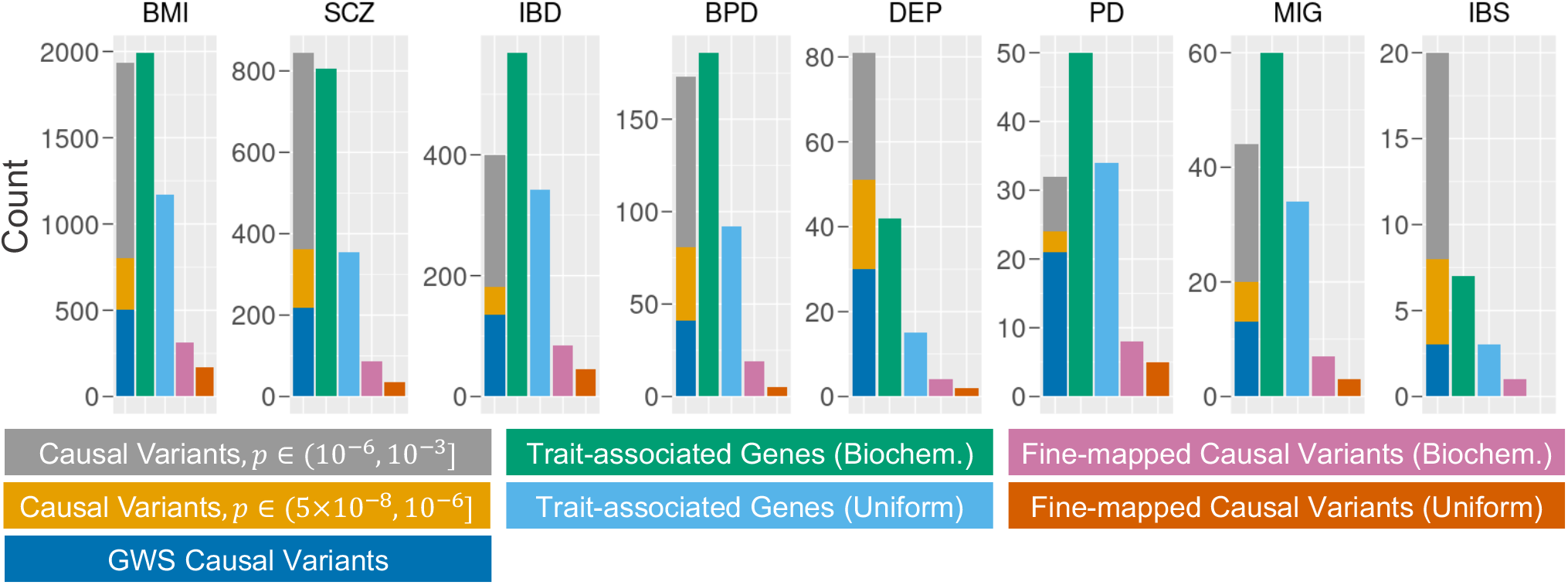
The biochemical prior substantially improves the fine-mapping of causal variants and trait-associated genes, related to Figure 5. Bar plots constructed as in Figure 5, except here we compare the numbers of fine-mapped causal variants and trait-associated genes obtained from the biochemical (purplish-red, bluish-green) versus the uniform (vermilion, sky blue) prior.

**Figure S15.**
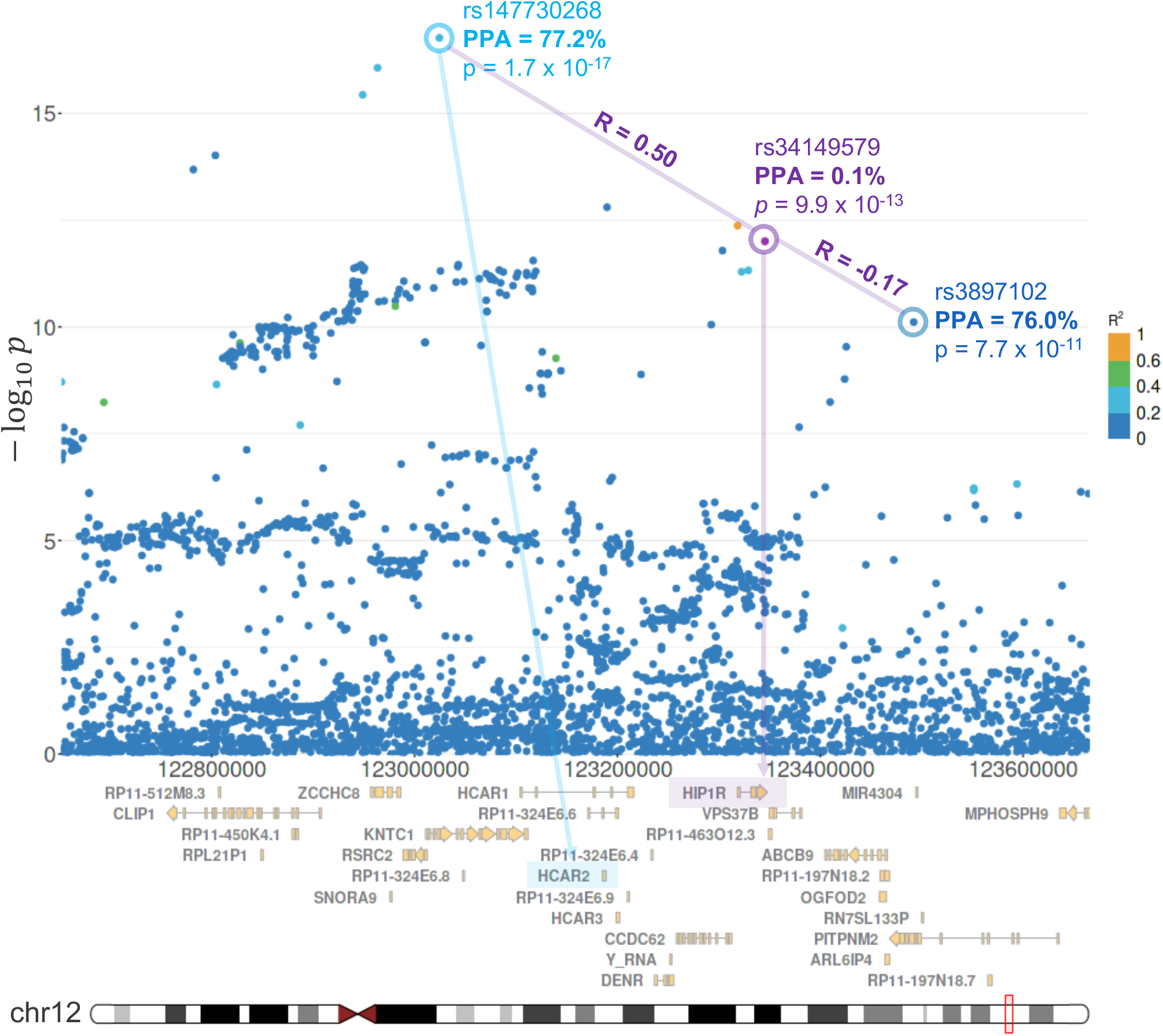
The BMI-associated missense variant in *HIP1R* is not causal, related to Figure 6 and Table 2. The association signals in a locus on chromosome 12, from a GWAS on BMI (Neale et al., 2021) in the UK Biobank (Bycroft et al., 2018). Each point is a genetic variant; its position on the chromosome (GRCh37 coordinates) is given on the x-axis, and its association to BMI is expressed as the negative logarithm of its GWAS p-value, *p*, on the y-axis. *HIP1R* has been implicated in BMI due to the genome-wide-significant association of a missense variant (rs34149579, purple) in the gene (Turcot et al., 2018). However, this variant is almost certainly not causal (PPA = 0.1%); its BMI association is a consequence of its correlations with two causal variants (probably rs147730268 and rs3897102) which flank the gene. The color scale indicates the correlation, *R*^2^, between the minor allele of the missense variant and the minor alleles of the other variants in the locus; the correlations with the putative causal variants are highlighted. rs147730268 is a predicted enhancer variant that we link to the gene *HCAR2* and four others (*DENR*, *KNTC1*, *RSRC2*, *ZCCHC8*). *HCAR2* is a well-known metabolic sensor with a plausible role in body weight regulation; *DENR* is thought to be involved in translation initiation; *KNTC1* is an essential component of the mitotic checkpoint; *RSRC2* has an unknown function which may involve RNA binding; *ZCCHC8* is involved in the degradation of non-functional RNAs, such as eRNAs (Stelzer et al., 2016). Prioritization on the basis of biological rationale might implicate *HCAR2* as a likely causal disease gene.

**Figure S16.**
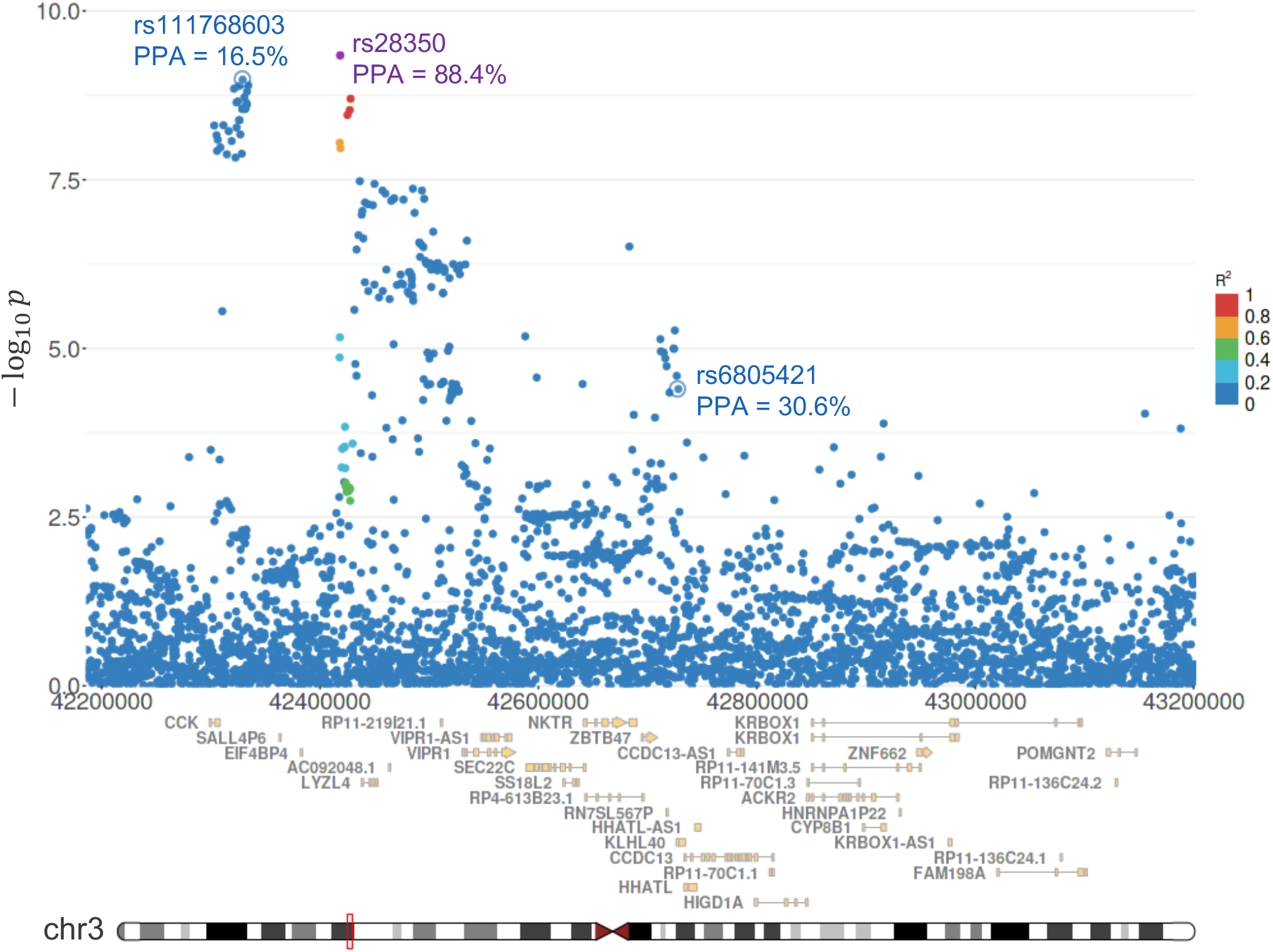
*CCK* has a 99.1% PPA in BMI, related to Figure 6. The association signals in two adjacent LD blocks on chromosome 3 (red box), from a GWAS on BMI (Neale et al., 2021) in the UK Biobank (Bycroft et al., 2018). The data are represented as in Figures 6 and S15. *CCK* is implicated by three independent association signals in this locus. The first is fine-mapped to variant rs28350 (purple), the lead variant. *CCK* is the only protein-coding gene we predict as a target for this variant, via an enhancer more than 110 kbp away from the gene. It is clear from the gene tracks plotted beneath the GWAS data that *CCK* is not the closest gene to that enhancer. The second independent association signal comprises the variant rs111768603 and the tightly correlated variants forming the cluster beneath it. While none of these variants is fine-mapped, many are linked to *CCK* via a proximal enhancer, and their integrated probability mass linked to *CCK* is substantial. Finally, rs6805421 is an unlikely but plausible causal variant, linked to *CCK* via an enhancer more than 420 kbp away from the gene. In the DNase-seq/ATAC-seq data from ENCODE, these three *CCK*-linked enhancers exhibit different patterns of chromatin accessibility across cell types, including the brain, intestinal mucosa, and nerve tissue.

**Figure S17.**
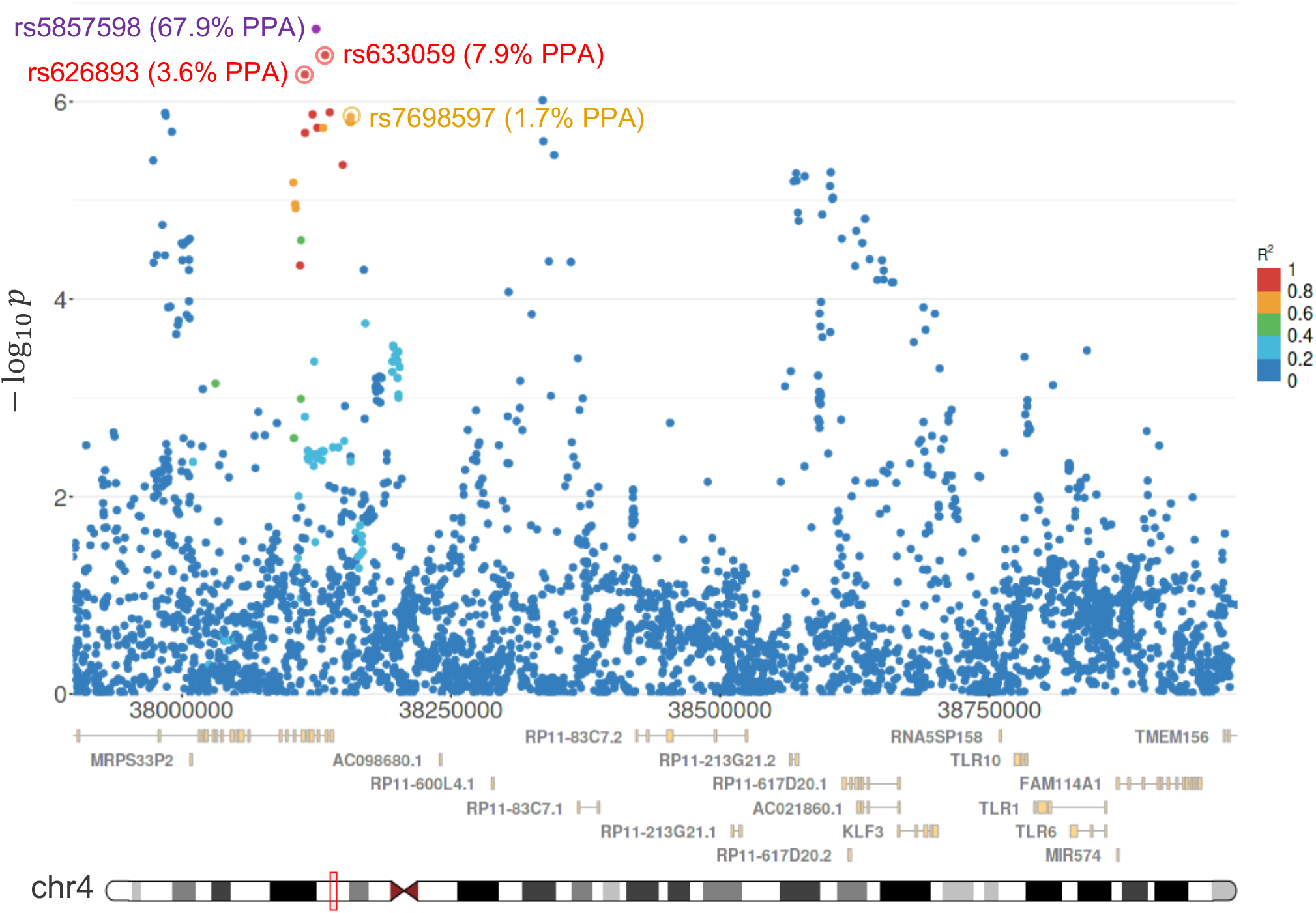
*TLR1* has an 89.0% PPA in IBD, related to Figure 6. The association signals in an LD block on chromosome 4 (red box), from a GWAS meta-analysis of IBD (de Lange et al., 2017). The data are represented as described above. *TLR1* integrates its PPA over several variants, predominantly the lead variant rs5857598 (purple, PPA = 67.9%) and several competing candidate causal variants (red, orange), circled. The lead variant is linked to *TLR1*, over 667 kbp away, via an enhancer with high DHS activity in several types of immune cells.

**Figure S18.**
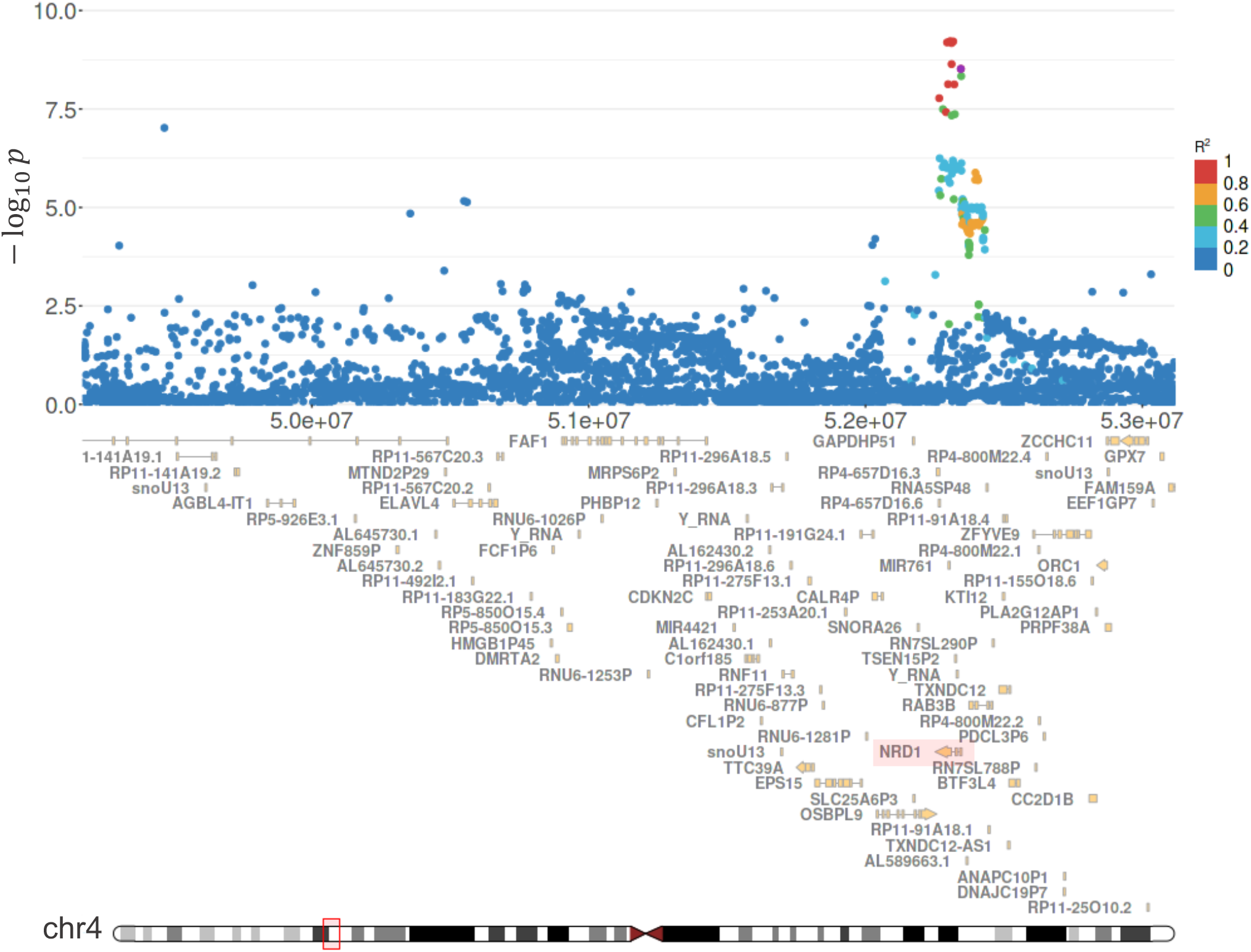
*NRDC* is a likely causal gene in depression, related to Figure 6. Our platform implicates *NRDC*, also named *NRD1*, with a 96.5% PPA from a GWAS of depression (Howard et al., 2019). Shown here are the GWAS data in the LD block containing *NRDC*, which integrates its PPA over a number of putative causal variants in the GWS association signal overlying the 5’ end of the gene. The purple point denotes the variant rs736756, which has the highest PPA (0.328) in the association signal, although it is not the lead variant. *NRDC* is the only gene linked to this association signal.

#### Supplemental tables

**Table S1.** Number of population-specific LD blocks, related to Figure 1. The number of LD blocks stratified across chromosome and super population in the 1000 Genomes Project Phase 3 cohort.

|  | AFR | AMR | EAS | EUR | SAS |
| --- | --- | --- | --- | --- | --- |
| 1 | 565 | 97 | 658 | 465 | 596 |
| 2 | 502 | 76 | 594 | 438 | 548 |
| 3 | 417 | 69 | 514 | 380 | 462 |
| 4 | 411 | 69 | 476 | 359 | 465 |
| 5 | 400 | 58 | 471 | 343 | 433 |
| 6 | 371 | 65 | 423 | 329 | 401 |
| 7 | 358 | 63 | 416 | 320 | 392 |
| 8 | 324 | 46 | 372 | 284 | 341 |
| 9 | 352 | 62 | 387 | 269 | 341 |
| 10 | 343 | 57 | 405 | 291 | 358 |
| 11 | 300 | 49 | 352 | 281 | 341 |
| 12 | 343 | 47 | 400 | 287 | 372 |
| 13 | 265 | 50 | 314 | 236 | 293 |
| 14 | 241 | 37 | 285 | 203 | 262 |
| 15 | 295 | 41 | 326 | 235 | 303 |
| 16 | 267 | 41 | 319 | 209 | 284 |
| 17 | 258 | 49 | 304 | 224 | 299 |
| 18 | 253 | 54 | 304 | 219 | 281 |
| 19 | 223 | 45 | 262 | 186 | 240 |
| 20 | 219 | 45 | 261 | 197 | 230 |
| 21 | 158 | 33 | 151 | 126 | 152 |
| 22 | 172 | 19 | 182 | 130 | 166 |
| *Total* | *7,037* | *1,172* | *8,176* | *6,011* | *7,560* |

**Table S2.** The frequency of the hotspot motif at LD block boundaries suggests that our borders are representative recombination hotspots, related to Figure 1. The percentage of LD block boundaries containing the well-known consensus sequence CCNCCNTNNCCNC within 1 kbp is shown. These results are consistent with previous reports that approximately 40% of recombination hotspots contain this motif (Myers et al., 2008).

| Population | % of LD Boundaries<br>with Hotspot Motif |
| --- | --- |
| AFR | 37.2 |
| AMR | 39.0 |
| EAS | 45.3 |
| EUR | 43.7 |
| SAS | 44.4 |

**Table S3.** The LD modularity objective accounts for sample size, related to Figures 1 and S1. Table summarizing the variables that shape the LD modularity objective for each 1KGP population (see the *Methods*). Sample estimates of the Pearson correlation *r*^2^ between genetic variants are biased; the term (*N* – 1)^−1^ corrects the bias in sample correlations between uncorrelated biallelic variants. The modularity penalty is the sum of the bias correction term and the threshold correlation 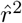. This penalty is subtracted from the sample correlations; only variants with a sample *r*^2^ in excess of the penalty will tend to co-cluster in the same LD block. The statistical underpinnings of LD modularity yield desirable properties. Larger bias corrections are applied to correlation estimates from smaller samples, but the significance threshold *α_N_* for calling correlated variants is implicitly relaxed in smaller samples.

| 1000 Genomes Threshold Characteristics |  |  |  |  |
| --- | --- | --- | --- | --- |
| Super Population | N<sup>o</sup> Haplotypes<br>($N$) | $r^2$ Correction<br>$-[N - 1]^{-1}$ | Modularity Penalty<br>$-([N - 1]^{-1} + \hat{r}^2)$ | $\alpha_N$ |
| African (AFR) | $661 \times 2 = 1322$ | $7.57 \times 10^{-4}$ | $3.26 \times 10^{-3}$ | .0380 |
| American (AMR) | $347 \times 2 = 694$ | $14.43 \times 10^{-4}$ | $3.94 \times 10^{-3}$ | .0981 |
| East Asian (EAS) | $504 \times 2 = 1008$ | $9.93 \times 10^{-4}$ | $3.49 \times 10^{-3}$ | .0606 |
| European (EUR) | $503 \times 2 = 1006$ | $9.95 \times 10^{-4}$ | $3.50 \times 10^{-3}$ | .0608 |
| South Asian (SAS) | $489 \times 2 = 978$ | $10.24 \times 10^{-4}$ | $3.52 \times 10^{-3}$ | .0634 |

**Table S4.** Number of universal LD blocks, related to Figures 1 and S5. The number of LD blocks found by the consensus algorithm over the four unadmixed super populations (AFR, EAS, EUR, SAS), and over all five super populations in the 1KGP Phase 3 cohort.

| Number of Consensus Blocks |  |  |
| --- | --- | --- |
|  | AFR+EAS<br>EUR+SAS | AFR+AMR<br>EAS+EUR+SAS |
| 1 | 178 | 25 |
| 2 | 161 | 11 |
| 3 | 142 | 15 |
| 4 | 119 | 13 |
| 5 | 148 | 18 |
| 6 | 117 | 14 |
| 7 | 119 | 13 |
| 8 | 103 | 8 |
| 9 | 112 | 12 |
| 10 | 114 | 15 |
| 11 | 96 | 6 |
| 12 | 100 | 11 |
| 13 | 92 | 15 |
| 14 | 81 | 12 |
| 15 | 87 | 17 |
| 16 | 99 | 18 |
| 17 | 96 | 13 |
| 18 | 97 | 15 |
| 19 | 74 | 13 |
| 20 | 84 | 14 |
| 21 | 54 | 9 |
| 22 | 53 | 6 |
| *Total* | *2,326* | *293* |

**Table S5.**
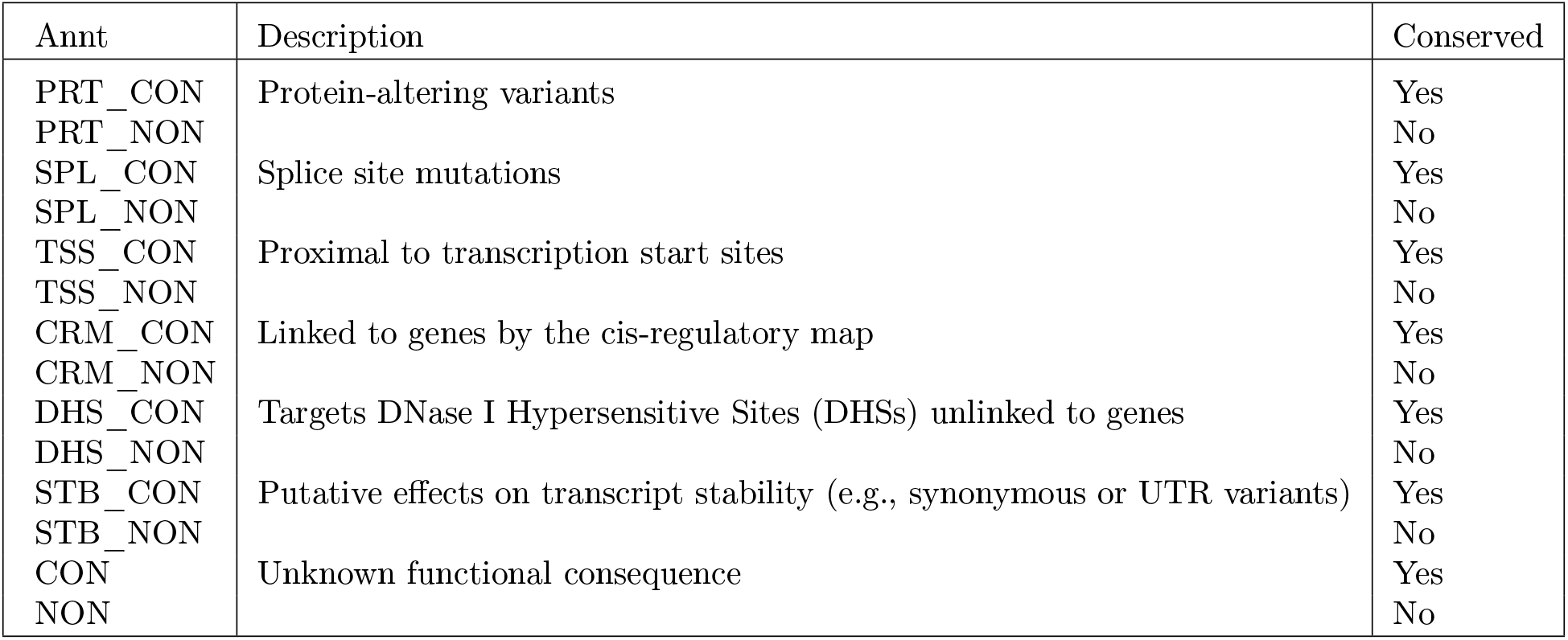
Waterfall logic for variant annotation, related to Figures 3–5. The variant annotations, ranked in order of decreasing severity. Variants are assigned the first (most severe) annotation that matches one of their consequences and their conservation status. Variant consequences are inferred from the Variant Effect Predictor (VEP; McLaren et al., 2016) and our variant-to-gene map. Conservation status was obtained by mapping each variant into conserved elements, computed from phastCons on a genome-wide multiple sequence alignment of 30 mammalian (27 primate) genomes. The phastCons conserved elements were obtained from the UCSC Genome Browser.

**Table S6.** Distance thresholds for scoring recall of known enhancer-gene interactions, related to Figure 2. Maximum genomic distance between an experimentally observed enhancer and a predicted enhancer for the same gene, in order to score the known interaction as predicted (recalled).

| Assay | Maximum enhancer distance |
| --- | --- |
| CRISPR mutagenesis | 300 bp |
| Fine-mapped and validated GWAS variant | 300 bp |
| CRISPRi (KRAB inactivation) | 4600 bp |
| CRISPRa (p300 activation) | 1000 bp |
| CRISPRa (VP16 activation) | 1000 bp |
| Specifically targeted genomic range | 0 bp |
| Fine-mapped eQTL variant | 300 bp |

**Table S7.** GWA studies from which the biochemical prior was learned, related to Figures 3–5. The compendium of GWA studies on which we ran the EM algorithm, to learn the parameters of the biochemical prior distribution. All of these studies passed our QC criteria. Note the recent publication dates.

| Trait | Study Citation | Publication Year |
| --- | --- | --- |
| AUDIT score | Sanchez-Roige et al. (2019) | 2019 |
| Alzheimer’s disease | Jansen et al. (2019) | 2019 |
| Anorexia nervosa | Watson et al. (2019) | 2019 |
| AST concentration in blood | Neale et al. (2021) | 2018 |
| Atopic triad | Watanabe et al. (2019) | 2019 |
| ADHD | Demontis et al. (2019) | 2019 |
| Bipolar disorder (Type I + II) | Mullins et al. (2021) | 2021 |
| Body mass index (BMI) | Neale et al. (2021) | 2018 |
| COPD | Sakornsakolpat et al. (2019) | 2019 |
| Coeliac disease | Neale et al. (2021) | 2018 |
| Depression | Howard et al. (2019) | 2019 |
| Genetic generalized epilepsy | ILAE Consortium (2018) | 2018 |
| Idiopathic pulmonary fibrosis (IPF) | Allen et al. (2020) | 2020 |
| Inflammatory bowel disease (IBD) | de Lange et al. (2017) | 2017 |
| Irritable bowel syndrome (IBS) | Eijsbouts et al. (2021) | 2021 |
| Migraine | Neale et al. (2021) | 2018 |
| Multiple sclerosis (MS) | Neale et al. (2021) | 2018 |
| Parkinson’s disease | Nalls et al. (2019) | 2019 |
| Primary biliary cirrhosis (PBC) | Cordell et al. (2015) | 2015 |
| Recurrent aphthous stomatitis (RAS) | Dudding et al. (2019) | 2019 |
| Schizophrenia | Psychiatric Genomics Consortium (2021) | 2022 |
| TSH concentration in blood | Teumer et al. (2018) | 2018 |

**Table S8.** BMI genes implicated by exome studies and their PPAs computed by our model, related to Table 2 and Figure S15. Left column, the union of all genes implicated in BMI by two association studies of protein-altering variants (Turcot et al., 2018; Akbari et al., 2021). Right two columns, the maximum PPA of each gene in BMI, body fat percentage, or waist-to-hip ratio after adjustment for BMI, computed with the two prior models.

| Gene | Biochemical Prior PPA (%) | Uniform Prior PPA (%) |
| --- | --- | --- |
| *GIPR* | 100.0 | 100.0 |
| *RAB21* | 100.0 | 100.0 |
| *ZFHX3* | 100.0 | 100.0 |
| *GPR151* | 100.0 | 99.7 |
| *SPARC* | 100.0 | 97.1 |
| *KSR2* | 99.7 | 87.9 |
| *PCSK1* | 99.6 | 76.5 |
| *RAPGEF3* | 96.8 | 42.1 |
| *ZBTB7B* | 96.0 | 94.2 |
| *ANKRD27* | 95.5 | 57.0 |
| *ENTPD6* | 94.9 | 84.6 |
| *MC4R* | 90.8 | 61.5 |
| *PRKAG1* | 78.8 | 33.3 |
| *CALCR* | 75.4 | 59.2 |
| *ZFR2* | 74.9 | 48.2 |
| *ROBO1* | 74.2 | 41.7 |
| *UHMK1* | 72.8 | 35.1 |
| *GPR75* | 65.0 | 30.2 |
| *DPP9* | 64.4 | 60.7 |
| *ACHE* | 63.2 | 49.8 |
| *UBR2* | 62.0 | 35.5 |
| *KIAA0586* | 46.0 | 14.4 |
| *ZNF169* | 41.5 | 38.2 |
| *HIP1R* | 40.4 | 38.5 |
| *KIAA1109* | 36.3 | 18.0 |
| *PDE3B* | 22.2 | 12.0 |
| *ANO4* | 10.2 | 6.9 |

#### Supplemental notes

##### Note S1. Comparison of LD partitions to other methods

In order to compare our LD partitions to previous studies of LD, we must first make an important distinction, between haplotype blocks and LD blocks. The haplotype block and the LD block are different concepts, with separate applications in genetics. Haplotype blocks are defined as contiguous genomic regions with little or no evidence of historical recombination. Heuristic methods to identify haplotype blocks already exist (Gabriel et al., 2002b; Wang et al., 2002; Wall and Pritchard, 2003) and are integrated into standard genomics software (Barrett et al., 2005; Chang et al., 2015). However, fine-mapping models must consider *all* nonzero allelic correlations, not merely tightly coupled sites with near-perfect correlation. The existing heuristics to define haplotype blocks are therefore not suitable for delineating LD blocks, because loci separated by historical recombination events may nevertheless exhibit substantial correlation. Moreover, the stringent criteria for defining haplotype blocks exclude much of the genome (Wall and Pritchard, 2003). We therefore forgo a comparison between our method to discover LD blocks and the rich literature on the delineation of haplotype blocks. Instead, we compare our algorithm to two previous approaches that define LD blocks for the purpose of applying multivariate statistical models to GWAS data. These are the algorithms LDetect (Berisa and Pickrell, 2016) and Big-LD (Kim et al., 2018, 2019).

LDetect is a signal processing heuristic that defines approximate LD blocks. Like the method introduced here, LDetect ensures that correlated variants are partitioned into the same LD blocks, which is the requisite condition for genome-wide statistical fine-mapping. Indeed, we find that the blocks defined here share common boundaries with the LDetect blocks (data not shown). The advantage of the approach taken here is that it enables fundamental discoveries about LD in the human genome. For instance, it unveils, in an unbiased way, the existence of a universal partition of the human genome. In addition, the blocks we find are small, which minimizes the computational burden for the statistical calculations. The median size of LDetect blocks is ≥ 1 Mbp across the AFR, EAS, and EUR super populations, compared to a median width < 300 kbp in the LD partitions introduced here. In the EUR super population, for example, LDetect parses the genome into 1,725 LD blocks (Berisa and Pickrell, 2016), whereas we find 6,011 blocks.

Big-LD, on the other hand, is a hierarchical graph clustering algorithm, although it clusters a different graph— according to different criteria—than we do here. The blocks defined by Big-LD are considerably smaller, with substantially higher levels of inter-block correlation, than those identified by our method or LDetect. The authors (Kim et al., 2018) report LD blocks with a median width less than 15 kbp and an average *r*^2^ of 0.155 between adjacent blocks (these results were computed only on Chr 22, but we presume the reported metrics are representative). In contrast, the average *r*^2^ between adjacent blocks in the partitions here is less than 0.0024 among the unadmixed super populations (and less than 0.0035 in AMR). The residual correlations across the Big-LD blocks violate a mathematical assumption of statistical fine-mapping, which requires that the variants in separate blocks be uncorrelated.

##### Note S2. Comparison of the gene fine-mapping model to ABC

Recently, a pioneering GWAS analysis method was reported (Nasser et al., 2021), in which the activity-by-contact (ABC) model was used to link fine-mapped variants in enhancers to target genes. That work is a literal interpretation of the same idea we pursued here. Namely, that the most direct approach for identifying disease genes is simply to identify the causal variants, and then link these to the target genes they affect. In Nasser et al. (2021), the variant fine-mapping and variant-to-gene links were made in separate computational steps: the variants were first fine-mapped and then subsequently linked to genes. The method presented here, which we developed in parallel to the ABC model, is based on the same concept, but we took a more mathematically nuanced approach. By integrating the variant-to-gene links directly into the statistical calculations, rather than deferring those links to a post-processing step, one can learn the prior probabilities that variants are causal. We show that this knowledge substantially improves the power to fine-map variants and detect trait-associated genes. Moreover, the approach we propose allows one to compute the probability that a gene is targeted by a causal variant (the gene PPAs). As we show here, computing gene PPAs profoundly increases the power to detect trait-associated genes, because it does not require first ascertaining the causal variant(s).

The authors of the ABC model describe some limitations of the approach (Nasser et al., 2021). The ABC model focuses on enhancer variants fine-mapped from GWS loci. The method assumes that variants with modest PPAs (≥ 10%) are causal if they fall within ABC enhancers, and sets of putative causal variants were linked to a single gene with the highest ABC score. Lastly, the underlying enhancer-to-gene map achieves its best performance on well-studied cell types and cell lines.

Here, we show how to consider all variants across the genome, even in loci that do not reach genome-wide significance. We propose a variant-to-gene map that is nearly comprehensive with regard to the types of variants it includes; it is not limited to polymorphisms in *cis*-regulatory elements. In addition, the statistical framework we suggest computes rigorous PPAs for variants, taking into account knowledge of whether they fall into various genomic features, such as enhancers. This approach obviates the need to assume that variants in enhancers are causal beyond a PPA threshold. Similarly, we avoid assumptions about the number of causal genes targeted by causal variants. Instead, we cluster groups of genes implicated by the same variants together, and leave the prioritization of putative causal genes as a post-processing step (which we implement with manual curation). Lastly, we put forward an enhancer-to-gene map that accurately predicts the effects of genetic variants on gene expression.

##### S3. Integrating out individual-level genotype data

In order to express the fine-mapping model in terms of publicly available summary statistics, we must rewrite the quantity 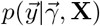 in those terms. We expand the skeletal derivation sketched in Benner et al. (2016). Recall the fundamental linear regression on a quantitative phenotype 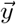:

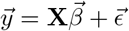

(Recall that we assume **X** has been mean-centered and standardized.) The maximum likelihood estimator (MLE) is the ordinary least squares (OLS) solution of this regression:

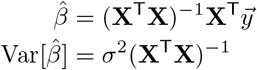

The z-scores computed from GWAS data are evaluated one variant at a time, from single-variant regressions of the form

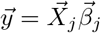

where 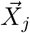 denotes the *j*-th column vector of the genotype matrix **X**, and *j* is the variant index, ranging from 1 to *m*. The z-score for the *j*-th variant is therefore:

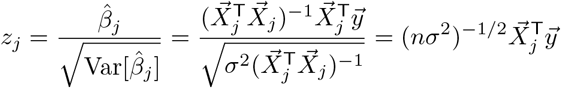

In matrix notation, this becomes

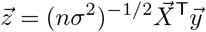

Noting that the variant correlation matrix **R** is none other than *n*^−1^**X**^┬^**X**, we can rewrite 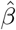 and **X** in terms of available summary statistics:

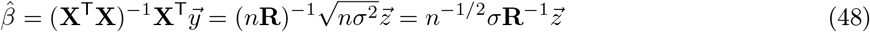

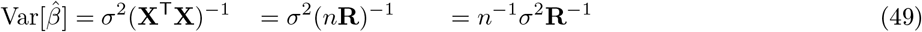

Leveraging the total law of probability for conditional probabilities, we next write

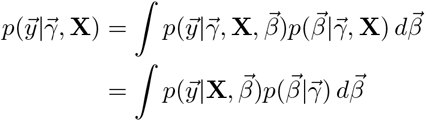

Notice that the probability of observing the phenotypes 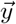, given the genotype matrix **X** and the true vector of effect sizes 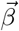, is identical to the probability of observing the MLE 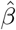 (which is uniquely determined by **X** and 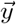):

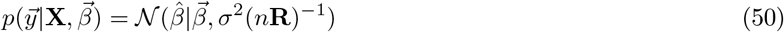

Then, recalling the definition

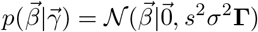

we can write

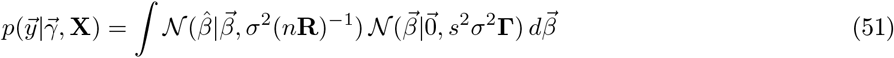

We identify this integral as the convolution of Gaussians. Let *f* and *g* be two independent normal distributions with mean *μ*_1_ and *μ*_2_, and variance 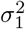 and 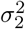, respectively. Then it is well known that the convolution of those distributions is itself a Gaussian, with mean *μ*_1_ + *μ*_2_ and variance 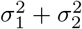:

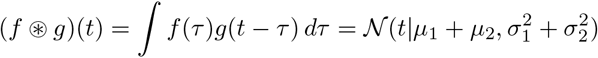

Using this fact, we rewrite equation (51) as a convolution of Gaussian distributions:

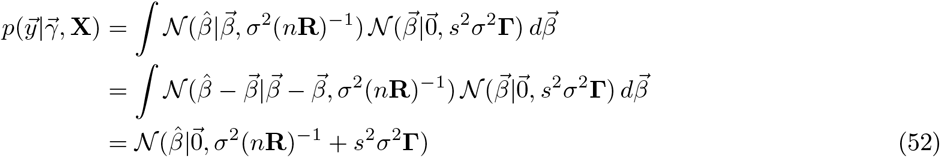

All that remains is a change of variables. Recall equations (48) and (49), and recall the properties of linear transformations of normal random variables. In particular, if we have a random variable *X* following a multivariate normal distribution

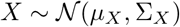

then a linear transformation of that random variable, *Y* = **A***X*, is also a multivariate normal:

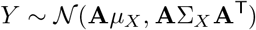

We have the linear transformation

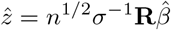

Therefore,

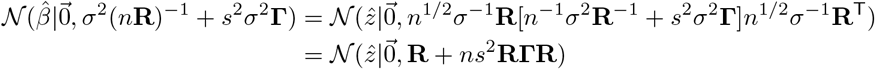

We now have a representation of the probability 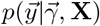 expressed in terms of GWAS summary statistics.

##### Note S4. Equivalence of fine-mapping model for continuous and case-control studies

###### Logistic regression for case-control studies

We show that the fine-mapping formulation derived in Note S3, for quantitative traits, also applies to case-control studies. As with continuous phenotypes, we assume the genotype matrix **X** is standardized, but now 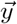 is an indicator vector, such that *y_i_* = 1 if the *i*-th study participant was a case, and *y_i_* =0 otherwise. Case-control studies apply logistic regression to model disease status as a function of genotype. The effect sizes in this logistic regression, denoted here by **λ**, enter via the formula:

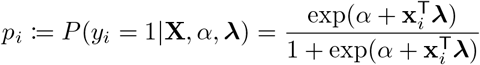

where **x**_*i*_ is the *i*-th row from **X**. The log-likelihood function is:

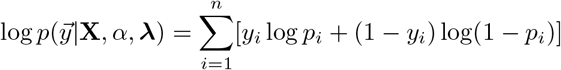

###### Distribution of the MLE

Similar to our treatment of continuous traits, we will find the MLE for the logistic effect sizes λ. We then convert the logistic effect sizes to their linear counterparts, 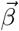. To begin, we leverage a classical result regarding the mean and variance of the MLE. Let 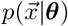 be a distribution with parameter ***θ***. Let ***θ***_0_ be the true value of the parameter and let 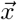 be a sample from 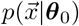. Let 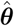 be the MLE of ***θ*** based on the sample 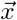. Also, define the log-likelihood function to be 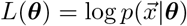. Then:

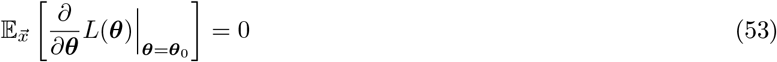

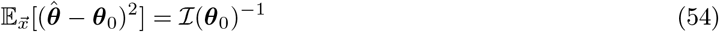

where 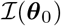 is the information matrix:

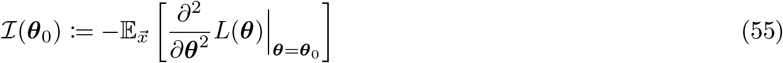

In order to apply this result to our log-likelihood function, a simple calculation, following Pirinen et al. (2013), shows that:

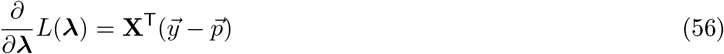

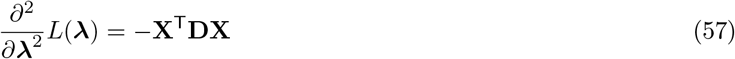

where **D** is the diagonal matrix with *D_ii_* = *p_i_*(1 – *p_i_*). Applying equation (53) to equation (56) yields

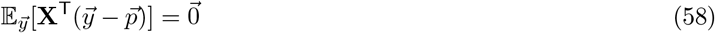

where 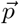 is evaluated at the true effect sizes **λ**_0_. Turning to equation (57), we see that the second partial derivative with respect to the effect sizes does not depend on 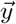. Therefore, the expectation in the definition from equation (55) may be omitted, leaving simply

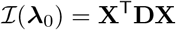

Now, we leverage the fact that effect sizes in GWA studies of complex traits are typically small: the genotype at any particular variant has a small effect on disease risk. We therefore approximate *p_i_* ≈ *p* ∀*i*, where *p* is the proportion of cases. Thus, we obtain the result

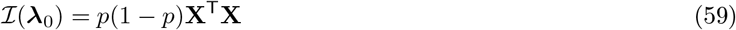

Consequently, from equations (54) and (59), we have

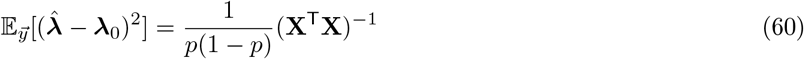

###### Finding the MLE

The MLE of **λ** is obtained at the unique zero of the score function:

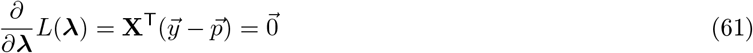

Recall that 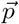 is a function of **λ**; to find the value of **λ** that satisfies equation (61), we follow Pirinen et al. (2013) and consider the Taylor expansion of *p_i_*. Consider a function of the form

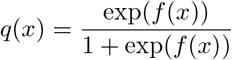

Then, we have the identity

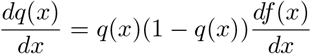

We use this identity to take the Taylor series expansion of *p_i_* as a function of **x**_*i*_ around the mean 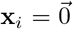 (recall the **X** is mean-centered):

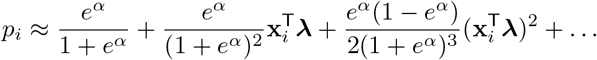

Again, leveraging the small magnitudes of the effect sizes, we approximate the value of *p_i_* with the first two terms of the Taylor expansion. Therefore, we have

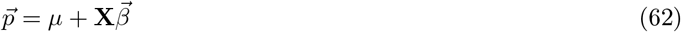

where

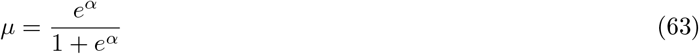

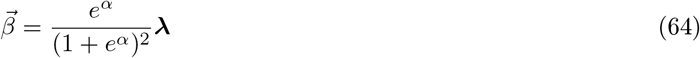

Note that *μ* is simply the proportion of cases. This means that

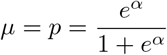

Therefore,

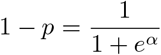

and

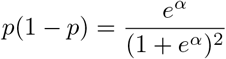

Thus, we now rewrite equation (64) as

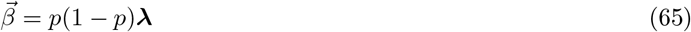

Substituting the expression for 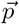, from equation (62), into the MLE solution from equation (61), we have:

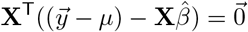

Solving this equation for 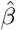 yields

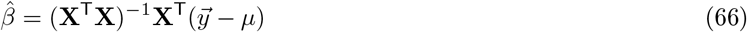

which is identical to the MLE from a linear regression on the mean-centered phenotypes! We are now equipped to estimate the distribution of 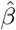. By equation (58), we have:

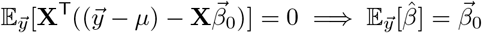

Likewise, by equation (60), and leveraging equation (65), we have:

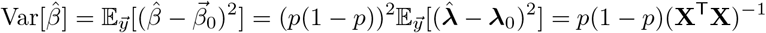

Therefore, we can approximate the distribution of 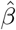 as

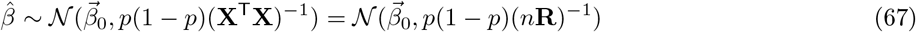

and we write

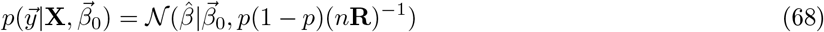

By comparison to equation (50), we see that the probability distribution of 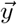 for case-control phenotypes follows precisely the same form as for quantitative traits, with *σ*^2^ = *p*(1 – *p*).

###### Wald statistics from logistic regression

For quantitative traits, the derivation of the fine-mapping model (based on summary statistics) requires a transformation from the effect sizes to z-scores. Here, we show that the same transformation holds for case-control studies via logistic regression. The *i*-th z-score from logistic regression is

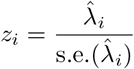

where

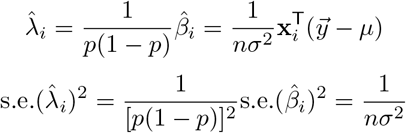

Therefore,

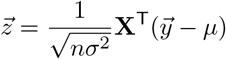

Combining this result with equation (66), we obtain:

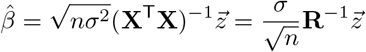

This is precisely the same relationship we obtained for the linear regression case, in equation (48).

##### Note S5. Setting the prior variance on causal effect sizes

Recall the prior on the effect sizes of causal variants, from equation (17). We also have the probability distribution over the phenotypes, expressed in terms of the summary statistics, from equation (18). Note that the variance of the causal effect sizes, *s*^2^, given as input to the model, is expressed in units of the trait variance, *σ*^2^. Moreover, formulation (18) depends only on *s*^2^, not *σ*^2^. For continuous traits, this means that we can specify the causal effects as a fraction of the trait variance, regardless of the value of *σ*^2^. For instance, if we set *s* = 0.05, then we assume 95% of the causal effects explain less than 10% of a standard deviation in the phenotype.

Specifying *s*^2^ relative to the trait variance is undesirable for case-control studies, however, because we have seen that *σ*^2^ = *p*(1 – *p*) for binary traits, where *p* is the proportion of cases (see Note S4). This means that the variance of causal effects would depend on the proportion of cases included in the GWAS, which is unrelated to the strength of the underlying genetic disease mechanisms. Therefore, rather than specifying *s*^2^ on the scale of the linear regression 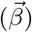, we instead provide *s*^2^ on the scale of the logistic effect sizes, **λ**. Recalling the relationship from equation (65), we have:

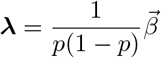

This means that a variance of *s*^2^ on the scale of the linear regression (on which our fine-mapping model is built) corresponds to a variance of *s*^2^[*p*(1–*p*)]^−1^ in the logistic effect sizes. Therefore, to express the causal effects in terms of the logistic regression, we simply multiply the desired variance by *p*(1 – *p*), to cancel out the aforementioned rescaling.

For a case-control study, we might set 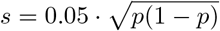, thereby assuming that the logistic effects have variance 0.05^2^. In order to interpret this variance in terms of odds ratios, we need to consider another source of scaling. In our model, WLOG, we presume the genotype matrix **X** is scaled so that all polymorphisms have unit variance. As described in Chen et al. (2015), this convention captures the tendency for variants with larger effect size to have lower minor allele frequency (MAF). Let *v* denote some causal variant tested in a case-control GWAS. The effect size of *v*, as reported by the GWAS (without standardizing **X**), will come from a distribution whose variance, 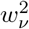, depends on the variant’s MAF. Let 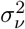 be the variance of *v* in the original (unscaled) genotype matrix. If we let 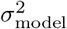 denote the logistic effect size variance in our model, then from Chen et al. (2015), we have:

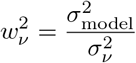

Recognizing that the unscaled genotype of a biallelic variant (coded as {0,1,2} to indicate the number of minor alleles) is a binomial random variable, we have

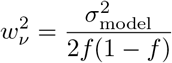

where *f* is the variant’s MAF. Moreover, it is well known that the effect size from logistic regression is the logarithm of the GWAS odds ratio (OR). Therefore, if we set 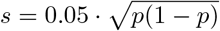, so that *σ*_model_ = 0.05, then 95% of the causal effect sizes will satisfy

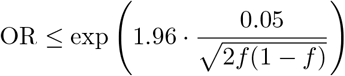

For *f* = 0.5 (a common variant), we have OR ≤ 1.15; for *f* = 0.01 (a relatively rare variant), OR ≤ 2.

##### Note S6. Fast computation of multivariate normal density

In Notes S3–4, we established the key result that

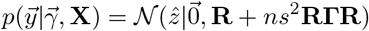

(Without loss of generality, we can assume all variables on the right hand side are specific to a particular LD block.) In Benner et al. (2016), the following additional result is derived:

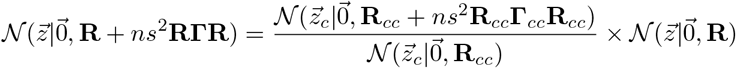

where the 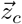, **R**_*cc*_, and **Γ**_*cc*_ denote partitions of each variable on the set of causal variants in a particular model 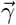. The term 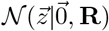 is effectively a constant that cancels out when we compute the probability of any causal model 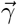. Therefore, within an LD block, 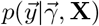 can be computed very efficiently, because it only requires the evaluation of low-dimensional multivariate normal densities (as opposed to the high-dimensional density defined on the thousands of variants in a typical block).

##### Note S7. The EM formulation minimizes cross-entropy at each M step

The cross-entropy loss function is widely used in machine learning, to minimize the Kullback-Leibler divergence between two probability distributions. Here, we show that each M step of our EM algorithm minimizes the crossentropy (i.e., discrepancy) between the prior and posterior probabilities of association for each variant. Therefore, this EM algorithm has a natural interpretation: at each iteration, it sets the prior probabilities to match the current estimate of posterior probabilities as closely as possible.

We begin by revisiting equation (40), which is the function we maximize at each M step. Let us adopt the notation

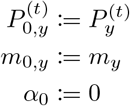

That is, we assign unannotated variants to the annotation index *a* = 0. Then we can simplify the notation in equation (40):

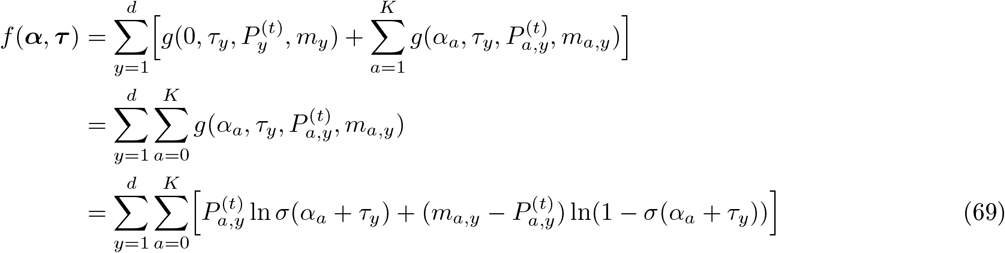

Now, we show that equation (69) is identical to the relevant cross-entropy function. The cross-entropy between two discrete probability distributions *p* and *q*, defined over a sample space 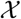, is given by

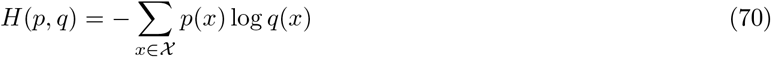

In our setting, each variant, in each trait, is a separate observation; each observation can take one of two values, indicating whether the variant is causal or not. Therefore, 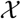 is the set of all possible causal states across all variants under consideration. Furthermore, we take *p* to be the *posterior* probability of the causal status for each variant, estimated at the current EM iteration. In addition, we denote by *q* the *prior* probability of the causal status for each variant, which is the distribution we want to optimize at each iteration. Finally, we define some new notation. Let 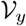 denote the set of variants tested in the GWAS for trait *y*. We also let 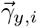 represent the *i*-th element in the true causal indicator vector for trait *y*; take *a*(*i*) to denote the biochemical annotation for the *i*-th variant. Then, the cross-entropy between our posterior and prior distributions may be written as follows:

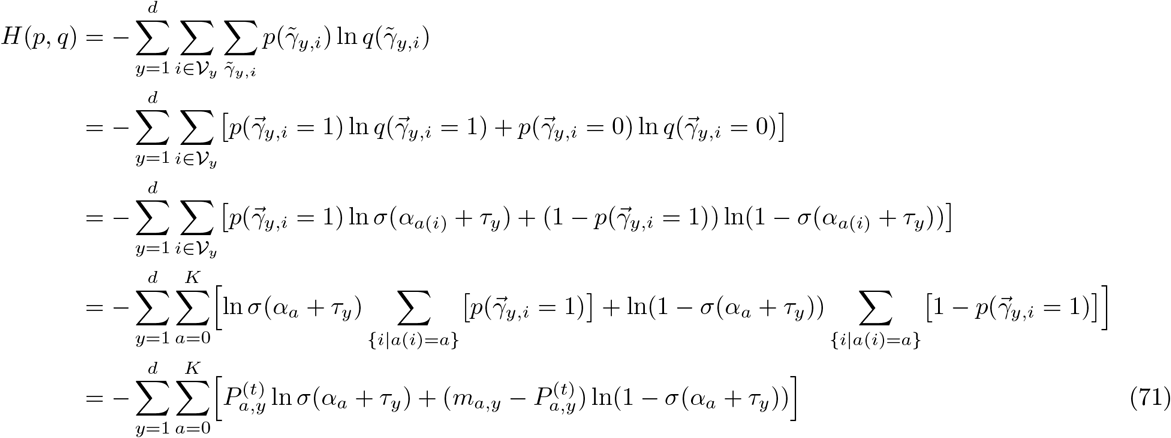

Comparing equations (69) and (71), we see that *f*(***α***, ***τ***) is nothing more than the negative of the cross-entropy. Therefore, maximizing f minimizes the cross-entropy between the posterior and prior probabilities of each variant, at the current EM iteration.

### Supplemental Files

**File S1.** Supplemental file ‘crispr_validation.tsv’ enumerates the known enhancer-gene interactions from CRISPR studies, and whether they are present in the enhancer-to-gene map.

**File S2.** Supplemental file ‘gtex_validation.tsv’ lists the causal variants fine-mapped by the GTEx consortium, together with their purported target genes. For each variant-gene association from GTEx, the file indicates whether the variant is in an enhancer (defined as a DHS with *R*^2^ > 0.84) and whether the interaction is observed in the enhancer-to-gene map.

**File S3.** Supplemental file ‘ENCODE_sample.tsv’ contains the ENCODE accession numbers and metadata for all chromatin accessibility data used to train the enhancer-to-gene map.

**File S4.** Supplemental file ‘cell_types.tsv’ comprises a surjective map from ENCODE-defined life stages and term names to qualitative cell type identifiers.

